# Positive Selection Screen Identifies Natural Product β-Catenin Inactivators

**DOI:** 10.1101/2025.08.27.671140

**Authors:** Matthew W. Boudreau, Vitor F. Freire, Sophie C. Corbett, Lucero Martínez-Fructuoso, Shilpa R. Shenoy, Wenyu Yu, Rohitesh Kumar, Christopher C. Thornburg, Rhone K. Akee, Brian D. Peyser, Qinqin Jiang, Jennifer Splaine, Jamie L. Pfaff, Benjamin C. Chandler, Dinah M. Abeja, Katherine A. Donovan, Jianwei Che, Benjamin L. Lampson, Mariana Cooke, Marcelo G. Kazanietz, Patricia Szajner, Jennifer A. Smith, Vidyasagar Koduri, Tanja Grkovic, Barry R. O’Keefe, William G. Kaelin

**Author notes:** Correspondence (W.G.K.).

## Abstract

Many genetically validated targets in cancer, including the transcription factor β-catenin (β-cat), have historically been viewed as undruggable. Cell-based phenotypic screening of chemical compounds can reveal new biological and pharmacological principles. Natural products are powerful probes because of their superior structural diversity, drug-like properties, and biological activities as compared to unoptimized synthetic compounds. We screened 326,304 natural product mixtures (40,744 extracts and 285,560 fractions derived from them) using mammalian cells expressing an oncogenic version of β-cat fused to a suicide protein. Multiple fractions degraded the β-cat fusion protein or drove it into a compartment where both fusion partners were apparently inactive. The active natural product from one of the latter specifically activates novel, but not classical, protein kinase Cs (PKCs) and thereby relocates β-cat to juxtamembrane vacuolar structures. These findings suggest a path for inactivating oncogenic β-cat and underscore the power of screening natural product collections with robust phenotypic assays.

---

Many genetically validated intracellular targets for various diseases are viewed as difficult to tackle with drug-like molecules because they are believed to lack appropriate hydrophobic pockets (*1, 2*). Examples of such targets in cancer include oncogenic versions of K-Ras, c-Myc, and β-cat, although there has been recent progress toward K-Ras inhibitors that leverage new structural insights as well as inactivation through induced proximity (*3, 4*). Indeed, two K-RAS inhibitors that specifically target K-RAS G12C are now approved (*5*).

The discovery that thalidomide-like drugs (“IMiDs”) are “molecular glues” that reprogram the cereblon ubiquitin ligase to target two otherwise undruggable oncogenic transcription factors, IKZF1 and IKZF3, for degradation (*6, 7*) has spurred interest in identifying additional molecules that can degrade specific target proteins by hijacking particular ubiquitin ligases (*8*). There are many other ways, however, that a small molecule could downregulate a protein of interest (POI). To search for degraders in a mechanism-agnostic fashion, we previously created a cell-based degrader positive selection (i.e. an “up assay”) that uses a bicistronic reporter encoding: 1) the POI fused to a modified deoxycytidine kinase (DCK*) that converts the non-natural nucleoside BVdU to a toxin and 2) green fluorescent protein (GFP) (*9*). GFP facilitates sorting for cells with the desired levels of the fusion protein and the quantification of cells that are both viable and retain the reporter. Genetic or pharmacologic perturbants that lower the abundance of the POI (and hence the POI fusion) promote the survival of GFP-positive viable cells in the presence of BVdU (*9, 10*). “Up assays” are less likely than “down assays” to yield trivial positives that simply interfere with cellular housekeeping functions or are otherwise toxic (*11*).

Mutations that cause the accumulation of active β-cat, including inactivating mutations of the *APC* or *AXIN* genes and activating mutations of *CTNNB1*, which encodes β-cat, are common in many cancers, including colon, gastric, liver, and uterine cancers (*12, 13*). β-cat functions in the Wnt pathway, which plays important roles in stem cell biology, development, and cancer (*12*). A first-in-class β-cat inhibitor that blocks β-cat binding to its partner TCF4 recently entered clinical trials (*14*). Other β-cat inhibitors, including putative degraders, have been described, but they do not appear to be robust (*15*).

To search for novel β-cat degraders, we engineered 293FT cells (which are β-cat-independent) to express an oncogenic β-cat variant (S37C) fused to DCK* (β-cat^S37C^-DCK*) or, as controls, unfused β-cat S37C (β-cat^S37C^) or unfused DCK* (**Fig. 1A**). In earlier whole genome CRISPR screens (*10, 16*) we noted that inactivation of thymidine kinase 1 (TK1) sensitized DCK*-positive cells to BVdU, presumably because TK1 monophosphorylates thymidine, which competes with BVdU, without significantly shifting the sensitivity of DCK*-negative cells to BVdU. We therefore inactivated TK1 in the 293FT cells using CRISPR/Cas9 knockout prior to introducing the different reporters (**Fig. 1B,C**). β-cat^S37C^-DCK*, like β-cat^S37C^, increased Axin 2 levels, a well-established β-cat/Wnt target gene, suggesting that the β-cat moiety in the β-cat^S37C^-DCK* fusion is properly folded and functional (**Fig. 1D**).

**Figure 1.**
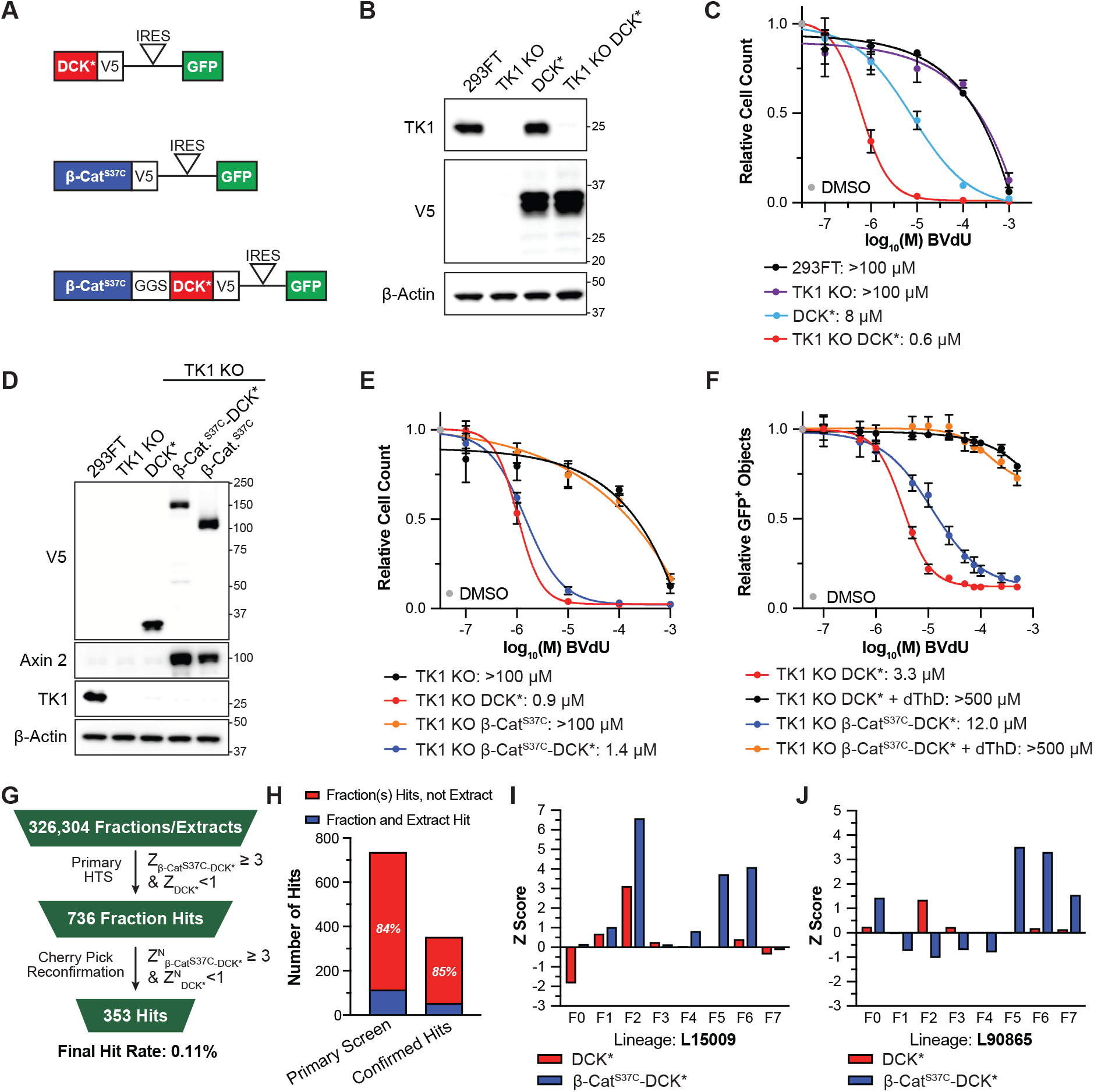
Establishment of β-Cat^S37C^-DCK* Positive Selection System for Natural Product Screening. (**A**) Reporter schematics. DCK*: variant deoxycytidine kinase with S74E/R104M/D133A substitutions. GGS: Gly-Gly-Ser spacer. IRES: internal ribosomal entry site. (**B**) Immunoblot analysis of 293FT cells expressing DCK* with or without TK1 inactivated using CRISPR/Cas9 (“TK1 KO”). (**C**) Relative survival of 293FT cells shown in (**B**) treated with the indicated concentrations of BVdU for 96 hours. n = 3 biological replicates. (**D**) Immunoblot analysis of 293FT cells expressing the reporters shown in (**A**). (**E**) Relative survival of 293FT cells depicted in (**D**) treated with the indicated concentrations of BVdU for 96 hours. n = 3 biological replicates. (**F**) Relative GFP+ objects for 293FT cells depicted in (**D**) treated with DMSO or dThD (final concentration 100 µM) for 24 hours followed by the indicated concentrations of BVdU for 96 hours in 384-well format. n = 3 technical replicates. (**G**) Summary of high throughput screening of natural product extract and fraction library and the hit scoring metrics used. Z: Z-score based on variation amongst experimental wells. Z^N^: Z-score based on the variation amongst the negative (^N^) control wells. (**H**) Number of positively scoring fractions for which corresponding extract did (blue) or did not (red) also score. (**I**,**J**) Z-scores from the primary screen of the L15009 lineage (**I**) and L90865 lineage (**J**).

As expected, the BVdU IC_50_ values for 293FT TK1 KO cells expressing β-cat^S37C^-DCK* or unfused DCK* were 2-3 logs lower than for the parental 293FT TK1 KO cells or 293FT TK1 KO cells expressing unfused β-cat^S37C^ (**Fig. 1E**). The sensitivity of 293FT TK1 KO cells expressing unfused DCK* to BVdU was not altered by cotreatment with the β-cat stabilizer CHIR99021 (**SI Fig. 1A,B**), which disrupts a β-cat phosphodegron by inhibiting the GSK3 kinase, or by coexpression of unfused β-cat^S37C^ (**SI Fig. 1C,D**). Therefore, β-cat signaling does not, *per se*, alter killing by BVdU in the presence of DCK*.

In pilot experiments, we confirmed that CRISPR sgRNAs directed against DCK* and, to a lesser extent, against β-cat, promoted the survival of the 293FT TK1 KO cells expressing β-cat^S37C^-DCK* (**SI Fig. 1E-G**). As expected (*9*), killing of 293FT TK1 KO cells expressing β-cat^S37C^-DCK* or unfused DCK* was reversed by the addition of 100 μM thymidine (dThD) in low-throughput assays and in 384-well plate format (**Fig. 1F**). The Z-prime (Z’) factors for the latter were >0.5 using the assay positive dThD as a surrogate for a specific true positive (**SI Fig. 1H,I**). Consistent with our previous experience with other DCK* fusions, multiple compounds that downregulate proteins by non-specifically interfering with transcription (e.g., actinomycin D), translation (e.g., zotatifin), or protein folding (HSP90 inhibitor: 17-AAG) did not score as hits in the β-cat^S37C^-DCK* reporter cells (**SI Fig. 1J**). BIX-02565, which inhibits translation by inhibiting RSK2, protected both β-cat^S37C^-DCK* and unfused DCK* (**SI Fig. 1J**), although the significance of this is unclear.

Encouraged by these findings, we next conducted a pilot high-throughput screen (HTS) using curated commercial and academic collections of bioactive compounds with known mechanisms of action (See *Methods*) (a total of 2,699 compounds) against the β-cat^S37C^-DCK* and DCK* reporter cells grown in 384-well plates (**SI Fig. 2A**). Experimental compounds were added to the assay plates using acoustic dispensing (day 0). BVdU was added on day one and GFP+ objects per well were measured on day five using a laser scanning imaging cytometer and the values were converted to Z-scores. 11 of the compounds scored in the β-cat^S37C^-DCK* cells and not in the unfused DCK* cells (Z ≥ 3 β-cat^S37C^-DCK*; Z < 1 for DCK*) (**SI Fig. 2A**). Five of the 11 were closely related to compounds that scored in cells expressing unrelated DCK* fusions in our experience and were therefore not studied further.

Of the remaining six compounds, one, **AZ-628**, downregulated both exogenous β-cat^S37C^-DCK* and endogenous β-cat^WT^ (**SI Fig. 2B**), suggesting it was a true positive. Intriguingly, **AZ-628** is a BRAF targeting, type 2 kinase inhibitor (*17*) that structurally resembles a compound dubbed WNTinib that inhibits downstream Wnt signaling, at least in part, by blocking the phosphorylation of EZH2 (*18*). WNTinib did not score in our assay and did not affect β-cat levels (**SI Fig. 2C**,**D**). We did not pursue **AZ-628** further because sensitivity to **AZ-628** does not track with β-cat dependence (in contrast with the good correlation with BRAF dependency) in public databases (DepMap Drug Repurposing Database, **SI Fig. 2E**,**F**) (*19, 20*) and was highly toxic at concentrations just above those used in our screen. These observations might reflect its known polypharmacology. Nonetheless, **AZ-628** might eventually illuminate a tractable path for degrading β-cat and, if so, could be a useful compound for additional structure-activity relationship studies and medicinal chemistry optimization.

The compounds found in nature are far more structurally diverse than synthetic chemicals. Screening natural product mixtures is challenging, however, for multiple reasons (*21*). For example, toxic compounds in such mixtures can cause false positives in “down” assays and false negatives in “up” assays (*11*). Moreover, it can be difficult to isolate and identify the active principle compound(s) from complex natural product mixtures when those mixtures score positively in a screen (*21*). We reasoned that both these challenges could be partially mitigated by marrying the performance characteristics of our screen to the use of partially fractionated natural product mixtures. Prefractioned natural product libraries offer many advantages including: sequestering toxic and nuisance compounds, concentrating minor active compounds, and simplifying downstream chemistry efforts (*22, 23*). The National Cancer Institute (NCI) Program for Natural Product Discovery (NPNPD) has prefractionated natural product organic extracts from the NCI natural product repository using a C8 solid phase extraction column into seven fractions (F1-F7) (*22*). Each unfractionated extract (F0), which is designated with a letter code based on its source (“L” = terrestrial plant; “M” = marine; H = fungal; “K” = marine plant) and a 5-digit unique identifier, is screened together with its seven subfractions (F1-F7) (e.g., “L90865_5” is the fifth fraction, F5, from the terrestrial plant extract L90865). We refer to each extract and its progeny as a lineage.

As an early risk assessment experiment, we screened two NPNPD challenge plates against β-cat^S37C^-DCK* and DCK* cells grown in 384-well plates as described above. These plates include extracts and fractions that either contain known pan-assay interference (PAIN) compounds or are highly pan-toxic in the NCI-60 assay. Our assay performed well, with excellent correlation between replicates and did not yield any positive hits (Z ≥ 3 β-cat^S37C^-DCK*; Z < 1 for DCK*, **SI Fig. 3A**), suggesting that our hit rate moving forward would not be prohibitively high. Excellent replicate-to-replicate correlations (R^2^=0.96-0.98) continued as the HTS began with the first 10 plates of fractionated natural product samples [440 extracts (F0) and 3080 fractions (F1-F7), **SI Fig. 3B,C**]. Thus, the remainder of the HTS campaign was conducted with a single replicate for β-cat^S37C^-DCK* and DCK* cells.

We ultimately screened 326,304 samples (40,744 F0 and 285,560 F1-F7) (**Fig. 1G**). As expected, the Z’ factors were excellent throughout the screen, Z-scores for the two different reporter lines were highly correlated, and most of the samples did not affect BVdU killing in either cell line (Z ≈ 0, **SI Fig 3D-F**). 736 samples corresponding to 695 lineages scored as potential β-cat degraders (Z ≥ 3 β-cat^S37C^-DCK*; Z < 1 for DCK*, **SI Fig 3E**). Excluded from these 736 hits were samples where the F0 scored, but none of the corresponding F1-F7 samples scored. 353 (48%) hits retested positively (here using the negative (N)-control based Z^N^ metric because of the presumed enrichment for true positives) after being cherry-picked and rescreened in duplicate, for a final hit rate of 0.11% (**Fig. 1G**).

The 353 hits, corresponding to 334 unique natural product lineages, originated from 70 different countries/regions (**SI Table 1**) and were derived from terrestrial plants (72.8%), marine sources (25.2%), marine plants (1.1%), and fungal/microbial sources (0.8%) (**SI Fig. 3H**). These percentages broadly mirrored the composition of the screened samples (**SI Fig. 3I**). Hits were derived from a range of taxonomies (**SI Table 1**).

Most of the hits came from the more lipophilic F5 and F6 fractions (**SI Fig. 3J**), likely because scoring in our assay requires cell permeability. For >80% of the hit fractions, the crude extract (F0) did not likewise score as a hit (**Fig. 1H**), perhaps because the F0 contained a compound(s) that masked the activity in the scoring subfraction or due to increased concentration of minor active compounds after separation into the individual fractions. Unsurprisingly, given the single column separation at this point, we noted 16 lineages where adjacent fractions scored. Both these observations were exemplified by the analysis of L15009 and L90865 (**Fig. 1I**,**J**).

322 hit fractions were separated into 22 subfractions (sF1-sF22) by preparative reverse phase HPLC (C18 column), yielding 7,084 subfractions (some hits were not subfractionated due to insufficient supply of the parent extract or other technical considerations) (**SI Fig. 4A**). It is important to note that the exact concentration (µg/mL) for each subfraction was not known and was estimated from an equally distributed mass balance of a 1 mg HPLC injection (see *Methods*). Therefore, subfractions were screened, in duplicate, at two doses (100 nL and 200 nL). Subfraction hits were called positive if they scored at either or both doses (see hit metrics in *Methods*). 435 subfractions, derived from 181 lineages, scored positively and were then counterscreened against 293FT TK1 KO cells expressing DCK*-IKZF1. 398 subfractions, derived from 162 lineages, promoted the survival of β-cat^S37C^-DCK* cells, but not DCK*-IKZF1, and were studied further (**SI Fig. 4A**). Scoring subfractions were well distributed across sF7-sF22 (**SI Fig. 4B**), reflecting the lipophilic nature of most of their parental fractions (**SI Fig. 3J**).

Given the limited amount of material available, we immunoblotted β-cat^S37C^-DCK* cells at a single time point (24 hours) and a single nominal concentration with at least one positive subfraction for each of the 162 extracts that scored positively. 36 lineages had at least one subfraction that modulated β-cat^S37C^-DCK* abundance *and* decreased the expression of the canonical β-cat target Axin 2. This included 24 extracts with subfractions that decreased β-cat^S37C^-DCK* abundance (hereafter referred to as “downregulators”) and, unexpectedly, 12 with subfractions that increased β-cat^S37C^-DCK* abundance (see examples of both in **Fig. 2** and **SI Fig. 4C**). We hypothesized that the latter inactivated the β-cat^S37C^-DCK* fusion protein, including both its β-cat and DCK* moieties, and thus protected cells against BVdU killing despite increasing exogenous β-cat^S37C^-DCK* levels (hereafter called “inactivators”).

**Figure 2.**
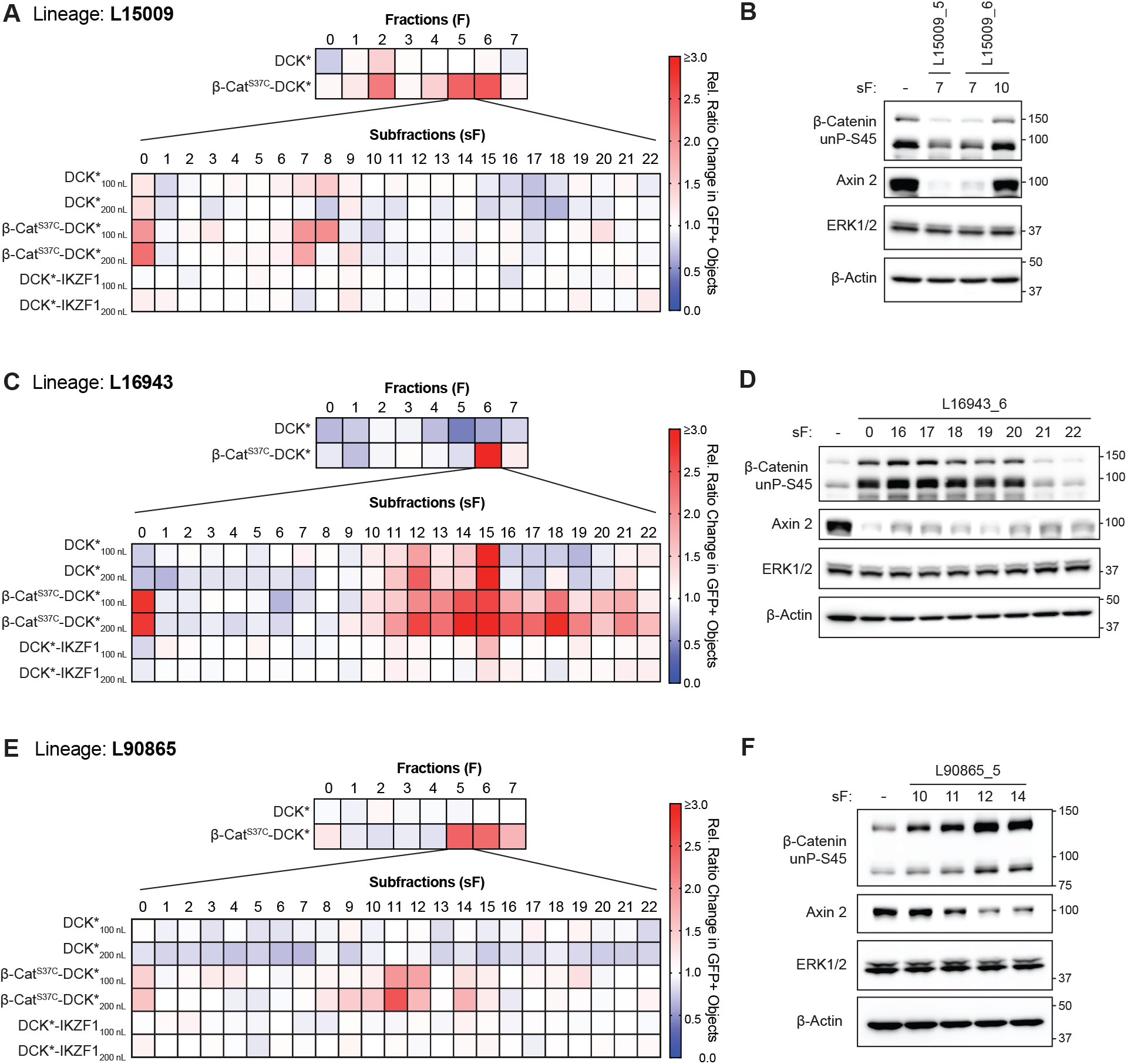
Two Apparent Modes of β-Cat Inactivation Displayed by Natural Product Subfractions. (**A**) Heatmap showing primary and secondary screening data, based on cell survival (GFP+ objects), related to lineage L15009 (fraction L15009_5, see fraction L15009_6 displayed in **SI Fig. 4C**). (**B**) Immunoblot analysis of 293FT TK1 KO β-cat^S37C^-DCK* cells treated with the indicated subfractions from (**A** and **SI Fig. 4C**) for 24 hours. Note differential mobility of exogenous β-cat^S37C^-DCK* and endogenous β-cat. (**C**) Heatmap showing primary and secondary screening data related to lineage L16943. (**D**) Immunoblot analysis of 293FT TK1 KO β-cat^S37C^-DCK* cells treated with indicated subfractions from (**C**) for 24 hours. (**E**) Heatmap showing primary and secondary screening data related to lineage L90865. (**F**) Immunoblot analysis of 293FT TK1 KO β-cat^S37C^-DCK* cells treated with indicated subfractions from (**E**) for 24 hours. For all heatmaps, coloring is representative of the relative GFP+ objects measured in a given test well. 100 nL and 200 nL refers to the transfer volume for subfraction dosing. n = 2 technical replicates for all subfraction data and data are shown as an average. All immunoblot images are representative of 2 biological replicates. “-” = DMSO treatment.

We used our previously described workflow to further isolate pure/semi-pure compounds from the target lineages on a small, testable milligram scale (0.1-1 mg). This enabled preliminary structural annotation based on multiple spectroscopic analyses (*24*). Samples were first tested in the β-cat^S37C^-DCK* and DCK* BVdU killing assays. Selective positives were then tested for β-cat^S37C^-DCK* modulation in western blot assays to prioritize hits worth scaling up for complete structural elucidation and further in-depth exploration.

From the downregulator lineage L15009, we ultimately isolated a known neolignan as a mixture of jatrointelignan A and B epimers (**1**) (**SI Fig. 5A**) (*25*). Compound **1** protected β-cat^S37C^-DCK* cells, but not DCK* cells, against BVdU treatment (**SI Fig. 5B**). The epimeric mixture was unstable and difficult to isolate, leading to a lack of material for follow-up studies, including western blot studies of endogenous β-cat. Active subfractions from L15009 downregulated endogenous β-cat and Axin 2 in SNU398 cells (β-cat^S37C^ hepatocellular carcinoma cell line, **SI Fig. 5C**). For the other downregulator lineages, either the pure compounds we isolated did not validate in secondary assays or we failed to arrive at a pure compound from the limited amount of extract available. The former could reflect false positives or failure to correctly isolate the active principle compound.

We then turned to the putative inactivators. Most of these came from the same plant family, *Euphorbiaceae* (9 out of 12), suggesting that they contain the same or similar compounds responsible for the inactivator phenotype. Surprisingly, multiple inactivators increased the abundance of the DCK* fusion protein (or unfused DCK*) and, more variably, the coexpressed GFP reporter, when tested against a panel of DCK* fusion proteins including DCK* fused to a β-cat^S37C^ C-terminal truncation mutant, suggesting that they non-specifically increase CMV promoter activity (**SI Fig. 6A**). The differential effects between the DCK* fusions and GFP could reflect differences in protein half-life and cap-dependent versus cap-independent translation. Nonetheless, protection by the inactivators was always specific to the cells expressing full-length β-cat^S37C^ fused to DCK* (**SI Fig. 6B-D**). We discovered, using mass spectrometry proteomics of DLD-1 colorectal cancer cells, that a partially purified inactivator from L16943 induced JunB and other AP-1 family members (**SI Fig 6E**). JunB was robustly induced in cells treated with inactivators (**SI Fig. 6F**,**G**). Induction of AP-1 could explain the activation of the CMV promoter, but not the specific protection of full-length β-cat^S37C^ fused to DCK* (*26-31*).

We hypothesized that the inactivators might aggregate or mislocalize β-cat. Treating DLD-1 cells, which have hyperactive β-cat due to an *APC* mutation, with active subfractions from L16943, as well as similar impure samples from L16945 (derived from the root bark of the same plant but not originally included in our screen) relocalized β-cat to what appeared to be large intracellular vacuoles abutting the cell membrane (**SI Fig. 7A**,**B**).

During the purification of the L90865 inactivator subfractions, we isolated sub-milligram quantities of two highly related lathyrane natural products: one containing a free primary alcohol (**2**, also called **lathyranol**) and the other in which that primary alcohol is acetylated (**3**, also called **lathyranol-19-acetate**) (**Fig. 3A,B**). Compound **2**’s primary alcohol is a unique substitution (i.e., on C19 of the geminal dimethylcyclopropane ring) for this class of natural products, which are rarely hydroxylated at this site (*32, 33*). Compound **2**, but not **3**, reduced β-cat activity, as determined by Axin 2 levels, in 293FT cells treated with the β-cat stabilizer CHIR99021, which inhibits GSK3, and mRNA expression of β-cat target genes *AXIN2, LEF1*, and *LGR5* in DLD-1 cells (**Fig. 3C, SI Fig. 7C**,**D**). The inactivators did not induce endogenous β-cat levels, in contrast to exogenous β-cat^S37C^-DCK*, presumably because the latter is driven by the CMV promoter (vide supra). Compound **2**, but not **3**, recapitulated the relocalization of β-cat described above (**Fig. 3D,E** and **SI Fig. 7E**). Of note, compound **2** was stable after 24-hour incubation in cell media with no hydrolysis of acetyl groups observed (**SI Fig 7F**).

**Figure 3.**
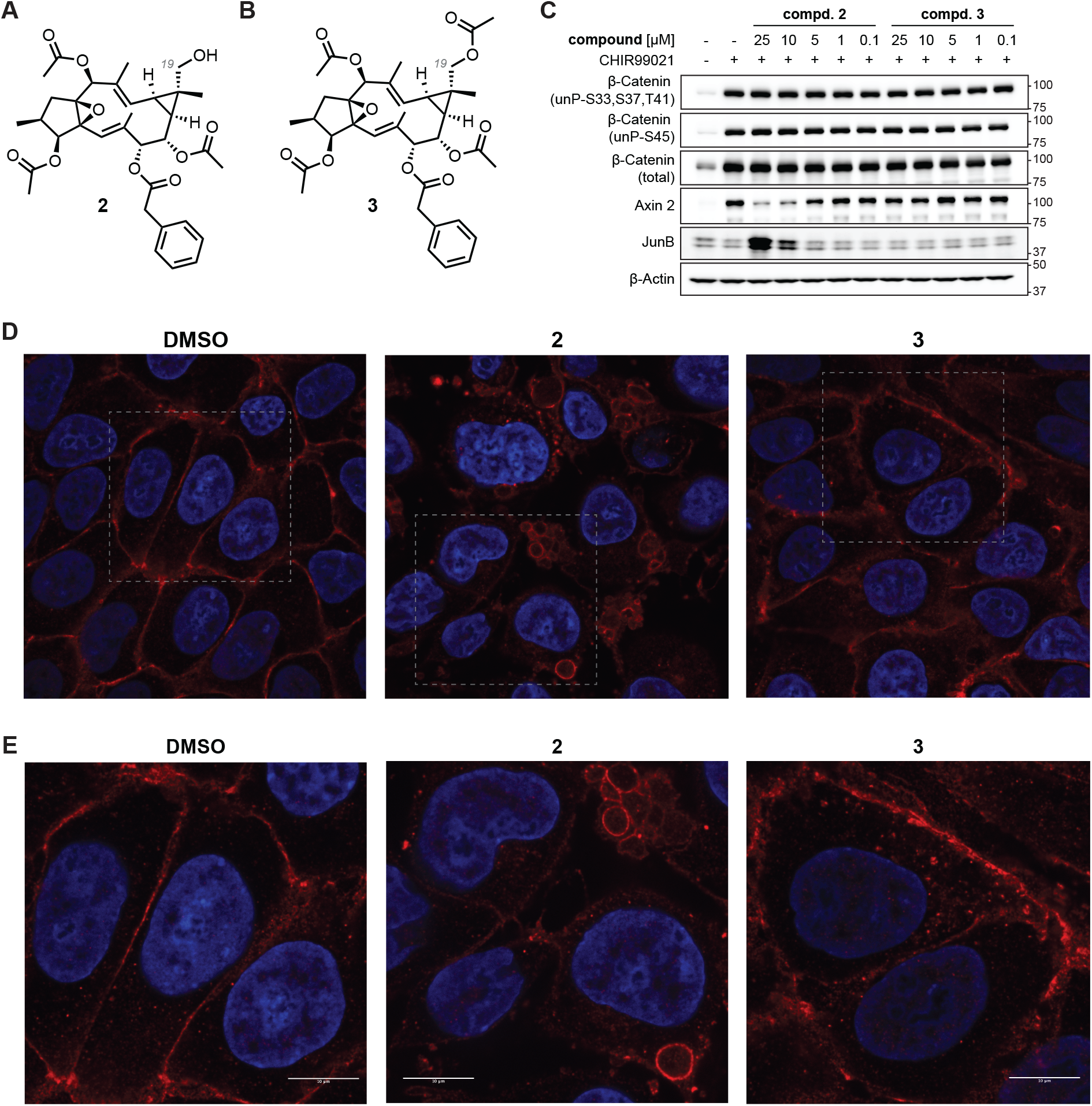
The Active Principle for Inactivator Lineage L90865 (compound 2) Inactivates and Relocalizes β-Cat. (**A**,**B**) Chemical structures of compounds **2** and **3**. The only difference between **2** and **3** is the acetylation of the C19 (grey) alcohol. (**C**) Immunoblot analysis of 293FT cells treated with CHIR99021 (5 µM) and **2** or **3** at the indicated concentrations and incubated for 24 hours. (**D**,**E**) Immunofluorescence images (W1: **D** or SORA enhanced: **E**) of DLD-1 cells treated with **2** or **3** at 25 µM concentration for 24 hours. Blue: DAPI stain, Red: β-Cat. Scale bar = 10 µm.

The structure of compound **2**, together with the requirement for its primary alcohol, led us to ask whether it was a PKC activator, especially since: 1) PKC activators have been isolated from *Euphorbiaceae* before (*32*), 2) other lathyrane natural products resembling compound **2** can activate PKCs (*34*), 3) PKC activation, via ERK and JNK, increases JunB transcription and stability (*28, 35*), 4) PKC activation can increase CMV promoter activity via AP-1 (including JunB) and cap-dependent translation via 4E-BP1 (*26-31*), 5) PKC activation has been reported to cause similar vacuolization (also referred to as ‘budding’) of β-cat (*36, 37*), and 6) PKC can phosphorylate β-cat on Serine 715 (*38, 39*), which is removed by the C-terminal truncation that abrogates protection by the inactivators we tested (**SI Fig. 6B-D**). Regarding the latter, we discovered that the S715A mutation substantially reduced the BVdU protection by 9 of the 12 inactivator lineages, including L90865 extract, fractions, and subfractions that produced compound **2** (**Fig. 4A,B, SI Fig.8A**). The S715A mutation did not, itself, affect reporter expression nor BVdU sensitivity (**SI Fig. 8B**,**C**). Moreover, compound **2** and the panPKC agonist ingenol-3-angelate (ING), but not compound **3**, induced the phosphorylation of multiple PKC substrates within 20 minutes of addition to cells (**Fig. 4C**). This coincided with phosphorylation PKCδ on a site indicative of PKCδ activation (*40, 41*) (**Fig. 4C**). In further support of a role for PKC, both ING and the panPKC activator phorbol-12-myristate (PMA), phenocopied compound **2** with respect to the induction of β-cat^S37C^-DCK*, induction of JunB, and downregulation of Axin 2 in the 293FT TK1 KO β-cat^S37C^-DCK* reporter cells (**SI Fig. 8D**). They also phenocopied compound **2** with respect to endogenous β-cat relocalization and target gene expression in DLD-1 cells (**SI Figs. 7C and 8E**). Relocalization of β-cat by ING occurred within 2-4 hours of treatment (**SI Fig. 8F**).

**Figure 4.**
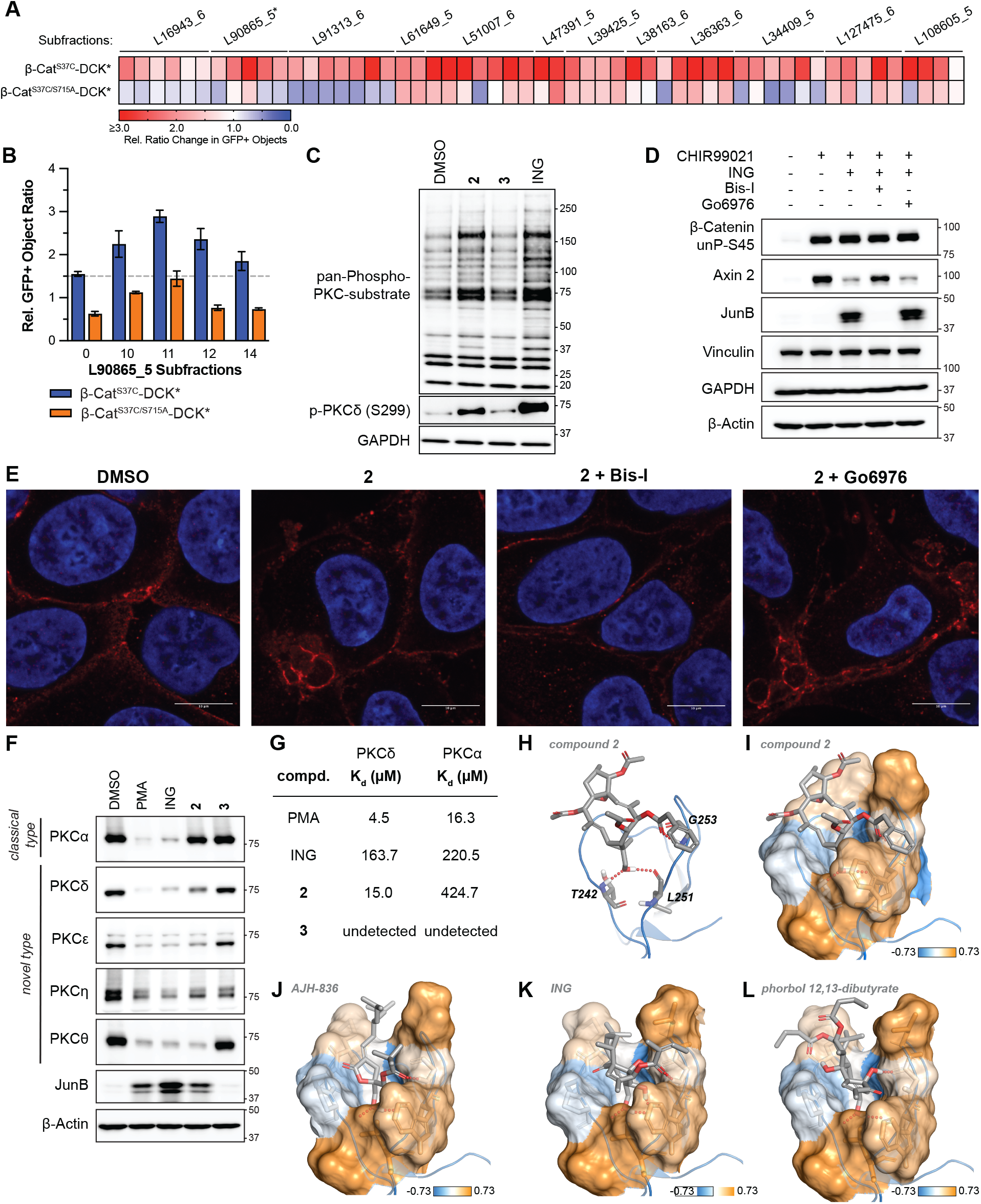
Compound 2 Specifically Activates Novel PKC Family Members. (**A**) Heatmap showing cell survival data (GFP+ objects) for 293FT TK1 KO β-cat^S37C^-DCK* and 293FT TK1 KO β-cat^S37C/S715A^-DCK* cells treated with positive subfractions from the indicated lineages for 24 hours followed by BVdU (50 µM) for 96 hours. Subfractions were dosed at a nominal concentration of 2.5 µg/mL. *: L90865_5 series was dosed at 5 µg/mL. n = 2 technical replicates. (**B**) Relative GFP+ objects for 293FT TK1 KO β-cat^S37C^-DCK* and 293FT TK1 KO β-cat^S37C/S715A^-DCK* cells treated with L90865 subfractions for 24 hours followed by BVdU treatment (50 µM) for 96 hours. n = 2 technical replicates. (**C**) Immunoblot analysis of 293FT cells treated, where indicated, with **2** (25 µM), **3** (25 µM), or ING (1 µM) for 20 minutes. n = 3 biological replicates. (**D**) Immunoblot analysis of 293FT cells treated, where indicated, with CHIR99021 (5 µM), ING (1 µM), Bis-I (1 µM), or Go6976 (0.5 µM) for 24 hours. n = 3 biological replicates. (**E**) Immunofluorescence of DLD-1 cells treated with indicated compounds **2** (25 µM), Bis-I (1 µM), or Go6976 (0.5 µM) for 24 hours. Blue: DAPI stain, Red: β-Cat. Scale bar = 10 µm. (**F**) Immunoblot analysis of 293FT cells treated, where indicated, with PMA (1 µM), ING (1 µM), **2** (25 µM), or **3** (25 µM) for 24 hours. n = 2 biological replicates. (**G**) Summary of the K_d_ values measured by microscale thermophoresis for each of the indicated compounds in the presence of full-length protein PKCδ or PKCα. (**H**) Three-dimensinal representation of the molecular docking of compound **2** to the C1b domain of PKCδ (PDB: 7LF3). Key hydrogen bonds are shown with red dots. (**I-L**) Space-filled models for the simulated binding of compound **2** (**I**) and the reported crystal structures for AJH-836 (**J**, PDB: 7LF3), ING (**K**, PDB: 7KO6), and phorbol 12,13-dibutyrate (**L**, PDB: 7KNJ). Key hydrogen bonds are shown with red dots, and the coloring is based on the Eisenberg consenous hydrophobicity scale (note: a value less than -0.73 is set to -0.73).

RNA sequencing (RNAseq) analysis of DLD-1 cells treated with PMA, ING, or compound **2** exhibited similar changes in gene expression (**SI Fig. 8G, H**). As expected with PKC activators, many genes were up- or downregulated. These changes were specific because the negative control compound **3** caused minimal changes in gene expression (**SI Fig. 8I**). Notably, gene-set enrichment analysis for MSigDB hallmark terms showed that Wnt-β-cat signaling was amongst the top 5 enriched terms when analyzing downregulated genes for PMA, ING, and compound **2** (**SI Fig. 8J-L**).

Notably, however, neither PMA, ING, nor other commercially available lathyrane natural products scored in our β-cat^S37C^-DCK* positive selection assay at any concentration tested, likely due to their toxicity (**SI Fig. 8M-P**). PMA and ING activate both classical and novel PKCs. Bisindolylmaleimide-I (Bis-I), which inhibits both classical PKCs and novel PKCs, but not Go6976, which only blocks the former, reversed the effects of ING with respect to β-cat, JunB and Axin 2 levels in the 293FT TK1 KO β-cat^S37C^-DCK* reporter cells and in 293FT cells treated with CHIR99021 (**Fig. 4D, SI Fig. 9A**). Moreover, Bis-I, but not Go6976, blocked the relocalization of endogenous β-cat by ING and compound **2** in DLD-1 cells (**Fig. 4E, SI Fig. 9B**). These observations suggested that β-cat inactivators, such as compound **2**, do so by activating a novel PKC. Consistent with this idea, doxycycline (DOX)-induced expression of consitutively active versions of either PKCδ or PKCθ, but not their catalytically inactive counterparts, phenocopied compound **2**’s effects on β-cat localization (**SI Fig. 9C-E**). ING still induced the formation of the β-cat cytoplasmic vacuoles in DLD-1 cells in which the classical PKCα was geneticallly inactivated by CRISPR/Cas9 editing (**SI Fig. 9F**,**G**).

Activation of PKC family members characteristically downregulates their apparent abundance in immunoblot assays, due to their decreased stability (*42-44*) or reduced immunoreactivity resulting from epitope phosphorylation (*45*). 293FT cells express multiple PKCs, although PKCα appears to be their primary classical PKC based on protein expression levels (**SI Fig. 9H**) and (*46, 47*)). As expected, PMA and ING activated PKCα and all 4 novel PKC family members in this assay (**Fig. 4F, SI Fig. 9I**). In contrast, the alcohol-containing **2**, but not the acetylated analogue (**3**), specifically activated the novel PKCs while sparing PKCα.

Compound **2**, but not compound **3**, bound directly to full-length human PKCδ, as determined by microscale thermophoresis, and bound to PKCδ with at least 20X higher affinity than to full length PKCα (**Fig. 4G, SI. Fig. 10A**,**B**). As expected, PMA and ING bound to both (**Fig. 4G, SI. Fig. 10A**,**B**). Due to the limited supply of compound **2**, these assays were not repeated in the presence of lipids, which are known to enhance the binding of PMA and ING to C1b domains further (*48*).

Molecular modeling studies suggested that the unique position of the primary alcohol found in compound **2** (on C19, **Fig. 3A**) could adopt a similar hydrogen bond network as known PKC agonists (**Fig. 4H-L**) when docked into the C1b domain of PKCδ (PDB: 7LF3, rat PKCδ, ∼98% conserved with human PKCδ). Molecular dynamic simulations indicated that this hydrogen bond network was stable (**SI Fig. 10C,D**). In sum, the simulations predict a binding mode for compound **2** that traverses the established hydrophobic binding cleft in the C1b domain, a stabilized hydrogen bonding network (Thr242, Leu251, and Gly253), and interaction and stabilization of the active ‘up’ configuration for the novel PKC-specific, diacylglycerol-toggling residue Trp252 (**Fig. 4H-I, SI Fig. 10C**,**D**) (*48*). This model also predicts that the acetylated compound **3** would not bind due to steric incompatibility with the shallow binding site and a lack of a free primary alcohol hydrogen bonding. This *in silico* model of compound **2** binding utilizes the crystal structure of only the C1b domain, which is highly conserved amongst classical and novel PKCs’ C1 domains, and thus does not take into account the rest of the protein structure that likely defines activity in cells (*49, 50*).

Finally, we obtained two synthetic PKC agonists, AJH-836 and YSE-028, that specifically activate the novel PKCs and not classical PKCs (*51, 52*). Both of these compounds are members of the diacylglycerol (DAG) lactone class and bind to the PKC C1 domains (*51, 52*). As expected, they bound selectively to PKCδ compared to PKCα (**SI Fig. 10E**,**F**). Moreover, they phenocopied compound **2** with respect to specific downregulation of novel PKC family members, JunB induction, and β-cat relocalizaton (**SI Fig. 10G**,**H**). Importantly, however, they, like PMA and ING (**SI Fig. 8M**,**N**), did not score in our positive selection assay (**SI Fig. 10I**,**J)**.

## Discussion

PKC activation was historically viewed to be oncogenic, dating back to classical studies with phorbol esters. Nonetheless, there is increasing evidence, including somatic inactivating mutations in cancers and functional studies, to support PKCs as potential tumor suppressors (*43*). The PKC agonist tool compounds PMA and ING have been reported to have anticancer properties, including in colorectal cancer cells (*53*), and, like compound **2**, inhibit β-cat function (*36, 54*). However, PMA and ING appear to be more toxic than **2**, likely in part because they, unlike **2**, also activate classical PKCs. Nonetheless, even DAG lactones that spare classical PKCs failed in our 5 day positive selection assay, perhaps due to off-target effects (*55-58*) or, in the case of YSE-028, susceptibility to cellular esterases (*52, 59*). Compound **2** would therefore be a better starting point for developing PKC agonists for cancer therapy.

Our screen illustrates the complementarity between chemical and genetic screens since the latter typically rely on gene inactivation or quantitative changes in gene expression and hence would not phenocopy PKC activation because activation of PKC requires that it undergo allosteric changes that prevent its autoinhibition (*43*). Indeed, we confirmed that overexpression of neither wild-type PKCδ nor PKCθ relocalized β-cat, in contrast to their constitutively activate variants. It will be informative to determine the breadth of mechanisms responsible for the different β-cat inactivators in our screen. In this regard, it will be important to retest some of the compounds that did not validate in our initial immunoblot assays, exploring different concentrations and timepoints given the 5-day time course of our screen.

In recent decades the pharmaceutical industry has favored target-based *in vitro* chemical screens over cell-based phenotypic chemical screens, largely because it can be difficult to identify the targets of hits emerging from the latter. Nonetheless, cell-based screens offer numerous potential advantages, including the ability to discover new biology, interrogate targets in their native contexts, and prioritize compounds that are bioavailable. Although our mechanistic insights related to compound **2** were hypothesis-based, many powerful biochemical (e.g., based on affinity capture) and genetic approaches (e.g., based on the generation of drug-resistant mutants) have been developed to identify targets for compounds scoring in phenotypic screens (*60*).

Similarly, screening natural product mixtures has largely been abandoned by the pharmaceutical industry (*21, 23, 61*), despite the fact that natural products have remarkable structural diversity and were the source of many important drugs in the past. Moreover, natural products often have superior absorption, distribution, metabolism, and excretion (ADME) properties compared to the compounds found in most synthetic compound libraries. The use of prefractionated natural product mixtures helps mitigate two main concerns regarding natural product screens: confounding effects caused by toxic chemicals in extracts and difficulties isolating the responsible compounds from extracts that score positively (*21, 23*). A major problem that remains relates to the ability to resupply extracts in sufficient quantities for purification and downstream chemical and biological analyses. Resupply is particularly important for molecules, such as compound **2**, that cannot be readily synthesized due to their structural complexity. These problems might be partially mitigated by using natural product collections derived from organisms (e.g., bacteria, fungi, certain plants) that can be easily cultured or cultivated at scale. The use of organisms with biosynthetic gene clusters amenable to genetic manipulation could enable the rapid generation of knockout strains for validation purposes and overproducing strains to enhance yields (*62*).

Drug development success at least doubles when based on a genetically validated target (*63, 64*). Somatic, and rarely germline, mutations have validated many otherwise undruggable targets in cancer. Cancers are often “addicted” to oncogenic proteins compared to normal cells, even when those proteins are active in the latter, providing a basis for a therapeutic window. Nonetheless, drugging the undruggable and finding drugs with broad therapeutic windows remains a challenge. Both challenges might be aided by screening chemical matter, including natural products, using positive selection assays analogous to the one described here.

## Acknowledgements

We thank members of the Kaelin laboratory, Gregory Wyant, and James DeCaprio for helpful discussions and ICCB-L technical staff for assistance in automation and library maintenance. We also thank Professor Hirokazu Tamamura, Institute of Science Tokyo, and Professor Neal Rosen, MSKCC, for gifting compounds YSE-028 and Zotatifin (respectively) for studies herein.

## Data and materials availability

RNAseq data was deposited in the public functional genomics data repository GEO; accession number GSE308354 (https://www.ncbi.nlm.nih.gov/geo/query/acc.cgi?acc=GSE308354). Token for reviewers: utuhugmmjdqbtat. Experimental high-resolution LC-MS and NMR data for compounds **2** and **3** and computational data for optimized conformers for NMR and ECD spectral calculations for compounds **2** and **3** have been deposited in the Harvard Dataverse (dataverse.harvard.edu) and can be found at https://doi.org/10.7910/DVN/1WVXAZ

## References

1. C. V. Dang, E. P. Reddy, K. M. Shokat, L. Soucek, Drugging the ‘undruggable’ cancer targets. Nat Rev Cancer 17, 502–508 (2017).

2. X. Xie et al., Recent advances in targeting the “undruggable” proteins: from drug discovery to clinical trials. Signal Transduct Target Ther 8, 335 (2023).

3. A. R. Moore, S. C. Rosenberg, F. McCormick, S. Malek, RAS-targeted therapies: is the undruggable drugged? Nat Rev Drug Discov 19, 533–552 (2020).

4. C. Weller et al., A neomorphic protein interface catalyzes covalent inhibition of RAS(G12D) aspartic acid in tumors. Science 389, eads0239 (2025).

5. J. M. Riedl et al., Emerging landscape of KRAS inhibitors in cancer treatment. Cancer Cell 44, 471–497 (2026).

6. G. Lu et al., The myeloma drug lenalidomide promotes the cereblon-dependent destruction of Ikaros proteins. Science 343, 305–309 (2014).

7. J. Kronke et al., Lenalidomide causes selective degradation of IKZF1 and IKZF3 in multiple myeloma cells. Science 343, 301–305 (2014).

8. M. Teng, N. S. Gray, The rise of degrader drugs. Cell Chem Biol 30, 864–878 (2023).

9. V. Koduri et al., Targeting oncoproteins with a positive selection assay for protein degraders. Sci Adv 7, (2021).

10. L. Duplaquet et al., Mammalian SWI/SNF complex activity regulates POU2F3 and constitutes a targetable dependency in small cell lung cancer. Cancer Cell 42, 1352–1369 e1313 (2024).

11. W. G. Kaelin, Jr., Common pitfalls in preclinical cancer target validation. Nat Rev Cancer 17, 425–440 (2017).

12. M. J. Parsons, T. Tammela, L. E. Dow, WNT as a Driver and Dependency in Cancer. Cancer Discov 11, 2413–2429 (2021).

13. C. Cui, X. Zhou, W. Zhang, Y. Qu, X. Ke, Is beta-Catenin a Druggable Target for Cancer Therapy? Trends Biochem Sci 43, 623–634 (2018).

14. K. P. Papadopoulos et al., A first-in-human, phase 1/2 trial of FOG-001, a β-catenin:TCF antagonist, in patients with locally advanced or metastatic solid tumors. J Clin Oncol 42, TPS3175 (2024).

15. M. A. McCoy et al., Biophysical Survey of Small-Molecule beta-Catenin Inhibitors: A Cautionary Tale. J Med Chem 65, 7246–7261 (2022).

16. B. L. Lampson et al., Positive selection CRISPR screens reveal a druggable pocket in an oligosaccharyltransferase required for inflammatory signaling to NF-kappaB. Cell 187, 2209–2223 e2216 (2024).

17. R. S. Vijayan et al., Conformational analysis of the DFG-out kinase motif and biochemical profiling of structurally validated type II inhibitors. J Med Chem 58, 466–479 (2015).

18. A. Rialdi et al., WNTinib is a multi-kinase inhibitor with specificity against beta-catenin mutant hepatocellular carcinoma. Nat Cancer 4, 1157–1175 (2023).

19. DepMap., DepMap Public 25Q2. Broad, depmap.org (2025).

20. R. Arafeh, T. Shibue, J. M. Dempster, W. C. Hahn, F. Vazquez, The present and future of the Cancer Dependency Map. Nat Rev Cancer 25, 59–73 (2025).

21. B. A. P. Wilson, C. C. Thornburg, C. J. Henrich, T. Grkovic, B. R. O’Keefe, Creating and screening natural product libraries. Nat Prod Rep 37, 893–918 (2020).

22. C. C. Thornburg et al., NCI Program for Natural Product Discovery: A Publicly-Accessible Library of Natural Product Fractions for High-Throughput Screening. ACS Chem Biol 13, 2484–2497 (2018).

23. A. G. Atanasov, S. B. Zotchev, V. M. Dirsch, T. International Natural Product Sciences, C. T. Supuran, Natural products in drug discovery: advances and opportunities. Nat Rev Drug Discov 20, 200–216 (2021).

24. T. Grkovic et al., National Cancer Institute (NCI) Program for Natural Products Discovery: Rapid Isolation and Identification of Biologically Active Natural Products from the NCI Prefractionated Library. ACS Chem Biol 15, 1104–1114 (2020).

25. J.-Y. C. Zhu, B.; Zheng, Y.-J.; Dong, Z.; Lin, S.-L.; Tang, G.-H.; Gu, Q.; Yin, S., Enantiomeric neolignans and sesquineolignans from Jatropha integerrima and their absolute configurations. RSC Adv. 5, 12202–12208 (2015).

26. P. Angel et al., Phorbol ester-inducible genes contain a common cis element recognized by a TPA-modulated trans-acting factor. Cell 49, 729–739 (1987).

27. W. Bruening, B. Giasson, W. Mushynski, H. D. Durham, Activation of stress-activated MAP protein kinases up-regulates expression of transgenes driven by the cytomegalovirus immediate/early promoter. Nucleic Acids Res 26, 486–489 (1998).

28. R. P. de Groot, J. Auwerx, M. Karperien, B. Staels, W. Kruijer, Activation of junB by PKC and PKA signal transduction through a novel cis-acting element. Nucleic Acids Res 19, 775–781 (1991).

29. A. Maass et al., Rational promoter selection for gene transfer into cardiac cells. J Mol Cell Cardiol 35, 823–831 (2003).

30. V. Kumar et al., Functional interaction between RAFT1/FRAP/mTOR and protein kinase cdelta in the regulation of cap-dependent initiation of translation. EMBO J 19, 1087–1097 (2000).

31. X. Liu et al., Phorbol ester-induced human cytomegalovirus major immediate-early (MIE) enhancer activation through PKC-delta, CREB, and NF-kappaB desilences MIE gene expression in quiescently infected human pluripotent NTera2 cells. J Virol 84, 8495–8508 (2010).

32. M.J. Durán-Peña, J. M. Botubol Ares, I. G. Collado, R. Hernández-Galán, Biologically active diterpenes containing a gem-dimethylcyclopropane subunit: an intriguing source of PKC modulators. Nat Prod Rep 31, 940–952 (2014).

33. F. Vela, A. Ezzanad, A. C. Hunter, A. J. Macías-Sánchez, R. Hernández-Galán, Pharmacological Potential of Lathyrane-Type Diterpenoids from Phytochemical Sources. Pharmaceuticals (Basel) 15, (2022).

34. S. Domínguez-García et al., A novel PKC activating molecule promotes neuroblast differentiation and delivery of newborn neurons in brain injuries. Cell Death Dis 11, 262 (2020).

35. M. Fujihara, M. Muroi, Y. Muroi, N. Ito, T. Suzuki, Mechanism of lipopolysaccharide-triggered junB activation in a mouse macrophage-like cell line (J774). J Biol Chem 268, 14898–14905 (1993).

36. S. Dupasquier et al., Modulating PKCα Activity to Target Wnt/β-Catenin Signaling in Colon Cancer. Cancers (Basel) 11, (2019).

37. M. Duong et al., Protein kinase C ϵ stabilizes beta-catenin and regulates its subcellular localization in podocytes. J Biol Chem 292, 12100–12110 (2017).

38. J. Xue et al., Tumour suppressor TRIM33 targets nuclear β-catenin degradation. Nat Commun 6, 6156 (2015).

39. P. V. Hornbeck et al., PhosphoSitePlus, 2014: mutations, PTMs and recalibrations. Nucleic Acids Res 43, D512–520 (2015).

40. J. Durgan, N. Michael, N. Totty, P. J. Parker, Novel phosphorylation site markers of protein kinase C delta activation. FEBS Lett 581, 3377–3381 (2007).

41. N. Kedei et al., Molecular systems pharmacology: isoelectric focusing signature of protein kinase Cdelta provides an integrated measure of its modulation in response to ligands. J Med Chem 57, 5356–5369 (2014).

42. R. Garg et al., Protein kinase C and cancer: what we know and what we do not. Oncogene 33, 5225–5237 (2014).

43. A. C. Newton, J. Brognard, Reversing the Paradigm: Protein Kinase C as a Tumor Suppressor. Trends Pharmacol Sci 38, 438–447 (2017).

44. N. Isakov, Protein kinase C (PKC) isoforms in cancer, tumor promotion and tumor suppression. Semin Cancer Biol 48, 36–52 (2018).

45. V. O. Rybin, S. F. Steinberg, Immunoblotting PKC-delta: a cautionary note from the bench. Am J Physiol Cell Physiol 290, C750–756 (2006).

46. J. R. van Senten, T. C. Møller, E. V. Moo, S. D. Seiersen, H. Bräuner-Osborne, Use of CRISPR/Cas9-edited HEK293 cells reveals that both conventional and novel protein kinase C isozymes are involved in mGlu(5a) receptor internalization. J Biol Chem 298, 102466 (2022).

47. P. Zong et al., TRPM2 enhances ischemic excitotoxicity by associating with PKCγ. Cell Rep 43, 113722 (2024).

48. S. S. Katti et al., Structural anatomy of Protein Kinase C C1 domain interactions with diacylglycerol and other agonists. Nat Commun 13, 2695 (2022).

49. T. A. Leonard, B. Rozycki, L. F. Saidi, G. Hummer, J. H. Hurley, Crystal structure and allosteric activation of protein kinase C betaII. Cell 144, 55–66 (2011).

50. P. M. Blumberg et al., Wealth of opportunity - the C1 domain as a target for drug development. Curr Drug Targets 9, 641–652 (2008).

51. M. Cooke et al., Characterization of AJH-836, a diacylglycerol-lactone with selectivity for novel PKC isozymes. J Biol Chem 293, 8330–8341 (2018).

52. T. Ishii et al., Synthesis and evaluation of DAG-lactone derivatives with HIV-1 latency reversing activity. Eur J Med Chem 256, 115449 (2023).

53. K. A. Benhadji et al., Antiproliferative activity of PEP005, a novel ingenol angelate that modulates PKC functions, alone and in combination with cytotoxic agents in human colon cancer cells. Br J Cancer 99, 1808–1815 (2008).

54. J. M. Lee et al., RORalpha attenuates Wnt/beta-catenin signaling by PKCalpha-dependent phosphorylation in colon cancer. Mol Cell 37, 183–195 (2010).

55. V. E. Marquez, P. M. Blumberg, Synthetic diacylglycerols (DAG) and DAG-lactones as activators of protein kinase C (PK-C). Acc Chem Res 36, 434–443 (2003).

56. J. Ann et al., alpha-Arylidene Diacylglycerol-Lactones (DAG-Lactones) as Selective Ras Guanine-Releasing Protein 3 (RasGRP3) Ligands. J Med Chem 61, 6261–6276 (2018).

57. M. Cooke et al., Differential Regulation of Gene Expression in Lung Cancer Cells by Diacyglycerol-Lactones and a Phorbol Ester Via Selective Activation of Protein Kinase C Isozymes. Sci Rep 9, 6041 (2019).

58. J. Das et al., Activation of Munc13-1 by Diacylglycerol (DAG)-Lactones. Biochemistry 62, 2717–2726 (2023).

59. E. Elhalem et al., Design, Synthesis, and Characterization of Novel sn-1 Heterocyclic DAG-Lactones as PKC Activators. J Med Chem 64, 11418–11431 (2021).

60. A. M. Freedy, B. B. Liau, Discovering new biology with drug-resistance alleles. Nat Chem Biol 17, 1219–1229 (2021).

61. J. W. Li, J. C. Vederas, Drug discovery and natural products: end of an era or an endless frontier? Science 325, 161–165 (2009).

62. J. R. Davison et al., Genomic Discovery and Structure-Activity Exploration of a Novel Family of Enzyme-Activated Covalent Cyclin-Dependent Kinase Inhibitors. J Med Chem 67, 13147–13173 (2024).

63. E. A. King, J. W. Davis, J. F. Degner, Are drug targets with genetic support twice as likely to be approved? Revised estimates of the impact of genetic support for drug mechanisms on the probability of drug approval. PLoS Genet 15, e1008489 (2019).

64. M. R. Nelson et al., The support of human genetic evidence for approved drug indications. Nat Genet 47, 856–860 (2015).

