## Supplemental Information for "Positive Selection Screen Identifies Natural Product β-Catenin Inactivators"

### This File Includes:

Supplementary Materials and Methods
SI Figs. 1-39
Legends for Table S1
References

### Supplementary Materials and Methods

#### *Cell Culture*

HEK293FT (293FT), DLD-1, SNU398, and RKO cells were originally obtained from American Type Culture Collection (ATCC). 293FT TK1 KO cells were generated by CRISPR-Cas9 editing (see below). 293FT, DLD-1, and RKO cells were grown in DMEM (Gibco, 11995065) supplemented with 10% FBS (GeminiBio, 100-106 or, for experiments involving the addition of doxycycline, GeminiBio, 100-800, which is tetracycline free) and 1% penicillin-streptomycin (Gibco, 15140122). SNU398 cells were grown in RPMI-1640 media (Gibco, 11875093) supplemented with 10% FBS (GeminiBio, 100-106; heat inactivated prior to use) and 1% penicillin-streptomycin (Gibco, 15140122). All cell lines were cultured at 37°C with 5% $CO_2$. Lentiviral infected cells were maintained in their corresponding media plus the desired selection agent (blasticidin: 10-12.5 µg/mL, puromycin: 2-4 µg/mL, or hygromycin: 200-300 µg/mL). Cell lines were occasionally tested for Mycoplasma infection using MycoAlert Mycoplasma Detection Kit (Lonza, LT07-318) or MycoStrip (InvivoGen, rep-mysnc-100).

#### *Chemicals*

The following compounds were purchased: DMSO (Sigma Aldrich, D2650), thymidine (dThD, MedChemExpress, HY-N1150), (*E*)-5-(2-Bromovinyl)-2'-deoxyuridine (BVdU, Fisher Scientific 50-536-044 from Chem-Impex International Inc. MFCD00058585), CHIR99021 (MedChemExpress, HY-10182), Ingenol-3-angelate (ING, Sigma Aldrich, SML1318), Phorbol-12-myristate (PMA, VWR, 80055-400), Bisindolylmaleimide-I (Bis-I, VWR, 80055-904 and MedChemExpress, HY-13867), Go6976 (Selleck, S7119), Euphorbia factor L1 (Selleck, S9410), Euphorbia factor L2 (MedChemExpress, HY-N5001), Euphorbia factor L3 (MedChemExpress, HY-N0562), Euphorbia factor L7A (MedChemExpress, HY-N8121), Euphorbia factor L7B (MedChemExpress, HY-N8118), 17-AAG (MedChemExpress, HY-10211), Actinomycin D (Thermo Fisher Scientific, 11805017), BIX-02565 (MedChemExpress, HY-16104), WNTinib (MedChemExpress, HY-160765), AZ-628 (MedChemExpress, HY-11004), doxycycline (Takara Bio, 631311), AJH-836 (from the Kazanietz-Cooke Laboratory University of Miami), YSE-028 (a gift from Professor Hirokazu Tamamura, Institute of Science Tokyo), Zotatifin (gift from Professor Neal Rosen, MSKCC), DAPI (Cell Signaling, 4083), Blasticidin (Thermo Fisher Scientific, NC9016621), Puromycin (Thermo Fisher Scientific, NC9138068), hygromycin (Life Technologies, 10687010).

#### *Immunoblotting*

Cell pellets were lysed in 1X RIPA buffer (Life Technologies, 89901) supplemented with protease inhibitor cocktail (Roche, 11836170001) and phosphatase inhibitor cocktail (Roche, 4906845001). Cell lysates were vortexed for 10 seconds, incubated on ice for 15 minutes, vortexed for 10 seconds again, and then clarified via centrifugation (16,100 x g) for 15 minutes at 4°C. The supernatants were then transferred to prechilled tubes and the protein concentrations were quantified using the Bradford protein assay (Bio-Rad, 5000006) according to the manufacturer's protocol. Equalized protein samples were then mixed with Laemmli SDS-Sample Buffer (Boston BioProducts, Inc., 6X-reducing, BP-111R) to a 1X concentration and heated to 70°C for 10 minutes. Samples were then either frozen at -80°C for later use or directly loaded and resolved on Novex Wedge Well protein gels (i.e., 4%-20% or 4-12% Novex WedgeWell, XP04205 or XP04125) using SDS running buffer diluted to 1X working concentration in water (10X SDS running buffer: 29 g Tris-Base (Thermo Fisher Scientific, BP1521), 144 g Glycine (Thermo Fisher Scientific, AA43497A5), 10 g SDS (sodium dodecyl sulfate, Sigma Aldrich, L3771) in 1000 mL ultrapure water). Gels were then transferred to nitrocellulose membranes using the Bio-Rad Trans-Blot Turbo Transfer System (Bio-Rad, 1704155 and Bio-Rad, 1704271) with the HIGH MW settings adjusted to achieve a 12–14-minute transfer time. For some immunoblots, especially when a high molecular weight protein was being compared to a low molecular weight protein (e.g.,

**Fig. 1D**), a wet PDVF transfer protocol was used instead wherein gels were transferred to PVDF 0.45 µm pore Immobilon Transfer Membranes (Millipore Sigma, IPVH00010) using a Mini-Trans-Blot electrophoresis system (Bio-Rad, 1703930 and Bio-Rad, 1703932). Transfers were done in precooled 1X transfer buffer (400 mL of methanol (Sigma Aldrich, 34860), 28.8 g glycine (Thermo Fisher Scientific, AA43497A5) 12.1 g Tris Base (Thermo Fisher Scientific, BP1521), 1600 mL of ultrapure water, pH = 8.3) on ice or with a frozen cool pack inside the tank for 2 hours at 54 volts. For most experiments uniform transfer was confirmed using a brief Ponceau Stain (Cell Signaling Technology, 59803) followed by TBST (6.06 g Tris-Base (Thermo Fisher Scientific, BP1521), 17.5 g sodium chloride (Thermo Fisher Scientific, BP35810) 2 mL Tween-20 (Thermo Fisher Scientific, BP337500), pH to 7.6 via addition of hydrochloric acid (37%, Sigma Aldrich, 258148) washing to remove the stain. Membranes were then blocked with bovine serum albumin (2 g in 40 mL of TBST (see above), Gold BioTechnology, A-420-500) for at least 1 hour at room temperature. Primary antibody was then added, and the membranes were incubated overnight at 4°C on a platform rocker. Membranes were then washed three times with TBST (10-20 minutes per wash) followed by incubation for at least 1 hour with the corresponding secondary antibody (horseradish peroxidase (HRP)-conjugated antibody) in TBST. After incubation (room temperature on a platform rocker), membranes were washed at least three times with TBST (20 minutes per wash). Membranes were then incubated in SuperSignal West Pico Plus solutions (1:1, Thermo Scientific, 1863099 and 1863098). Membranes were then chemiluminescent and colorimetric imaged with a Bio-Rad ChemiDoc MP Imaging system (Version 3.0.1.14) and images contrast/brightness/rotation were uniformly adjusted with High/Low/Gamma settings within the Image Lab 6.1 software (Bio-Rad SOFT-LIT-170-9690-ILSMAC-V-6-1). For all immunoblot membranes equal loading was confirmed by blotting for a loading control and/or by Ponceau staining. For some composite figures, different loading controls were used for different panels within the figure because of overlap between the molecular weight of the protein of interest and a loading control protein. For example, in **Figs. 2B, D, F,** β-cat and β-actin were from the same membrane and Axin 2 and ERK1/2 were from the same membrane. Not all the loading controls were included in some figures (e.g. **Fig. 4F**) for space and aesthetic reasons. Some images were also previously taken using the Azure biosystem C600 using the instrument software to get chemiluminescent and colorimetric images.

***Antibodies***

The following antibodies were used: anti-TK1 (Cell Signaling Technology, 8960), anti-V5-HRP (Life Technologies, R96125, 1:9000), anti-β-Actin-HRP (Cell Signaling Technology, 5125S, 1:5000), anti-Axin2 (Cell Signaling Technology, 2151), anti-β-Cat unP-S45 (Cell Signaling Technology, 19807S, 1:2000), anti-β-Cat unP-S33/S37/T41 (Cell Signaling Technology, 8814S, 1:2000), anti-β-Cat total (Cell Signaling Technology, 8480, 1:2000), anti-β-Cat total (Santa Cruz Biotechnology, SC-7963), anti-ERK1/2 (Cell Signaling Technology, 4695), anti-JunB (Cell Signaling Technology, 3753), anti-GAPDH (Cell Signaling Technology, 5174), anti-PKCα (Cell Signaling Technology, 59754), anti-PKCβ (Cell Signaling Technology, 46809), anti-PKCδ (Cell Signaling Technology, 2058), anti-PKCε (Cell Signaling Technology, 2683), anti-PKCη (Abcam, ab179524), anti-PKCθ (Cell Signaling Technology, 13643), anti-Phospho-(Ser) PKC Substrate (Cell Signaling Technology, 2261), anti-Phospho-PKC (pan) (beta II Ser660) (Cell Signaling Technology, 9371), anti-PKCδ (phosphor S299) (Abcam, AB133456), anti-DCK (Abcam, ab151966), anti-GFP (Cell Signaling Technology, 2956), anti-FLAG (Cell Signaling Technology, 14793), anti-vinculin (Sigma-Aldrich, V9131), anti-rabbit-HRP (Thermo Fisher Scientific, NC9611376, secondary antibody), anti-mouse-HRP (Thermo Fisher Scientific, NC9491974, secondary antibody), anti-mouse-AlexaFluor Plus 555 (Life technologies, A32727, 1:500), anti-rabbit-AlexaFluor Plus 488 (Life Technologies, A32731, 1:500). All primary antibodies were used at a 1:1000 dilution unless otherwise stated. All secondary antibodies were used at a 1:10,000 unless otherwise stated.

### ***Plasmids***

*Generation of DCK*, β-cat$^{S37C}$-DCK*, and β-cat$^{S37C}$ IRES GFP Vectors*

pLX304-DCK*-IRES-GFP and pLX304-DCK*-IKZF1-IRES-GFP vectors were described previously (*1*). The β-cat *cDNA* in β-cat-pDONR223 (obtained from Addgene 81734) was PCR amplified (without a stop codon) and with corresponding gateway overhangs (see primers section). The PCR product was gel extracted (Qiagen, 28706), purified, and shuttled into pDONR223 by Gateway BP Clonase Reaction (Thermo Fisher Scientific, 11789020) as per manufacturer's instructions to yield β-cat(no stop)-pDONR223. The β-cat in this plasmid was then shuttled into pLX304-GW-DCK*-IRES-GFP (Addgene, 176291) or pLX304-GW-IRES-GFP (Addgene, 176288) destination vector by Gateway LR recombination (Life Technologies, 11791100) as per manufacturer's instructions. The corresponding pLX304-β-cat-DCK*-IRES-GFP or pLX304-β-cat-IRES-GFP vector was then subjected to site-directed mutagenesis (using single 5' phosphor-primer PCR, see primer section) to make the pLX304-β-cat$^{S37C}$-DCK*-IRES-GFP and pLX304-β-cat$^{S37C}$-IRES-GFP vectors.

β-cat (no stop)-pDONR223 was subjected to site-directed mutagenesis (using single 5' phosphor-primer PCR, see primer section) to yield the corresponding β-cat$^{S37C}$ pDONR223. β-cat$^{S37C}$ was then shuttled into pLX302 (Addgene, 25896) destination vector by Gateway LR recombination (Life Technologies, 11791100) as per manufacturer's instructions yielding a pLX302 β-cat$^{S37C}$ vector.

*Generation of C-Terminal Truncated 1-664 β-cat$^{S37C}$-DCK* Vector*

The β-cat cDNA in β-cat$^{S37C}$ (no stop)-pDONR223 (as a template) was PCR amplified with a 3' PCR designed to eliminate the open reading frame C-terminal to residue 664 (see primer section). The resulting Gateway compatible PCR product was gel extracted (Qiagen, 28706), purified, and shuttled into pDONR223 by Gateway BP Clonase Reaction (Thermo Fisher Scientific, 11789020) as per manufacturer's instructions to yield 1-664-β-cat$^{S37C}$ pDONR223. 1-664-β-cat$^{S37C}$ encoding sequence was then shuttled into pLX304-GW-DCK*-IRES-GFP (Addgene, 176291) destination vector by Gateway LR recombination (Life Technologies, 11791100) as per manufacturer's instructions.

*Generation of β-cat$^{S37C/S715A}$-DCK* Vector*

β-cat$^{S37C}$ (no stop)-pDONR223 was subjected to site directed mutagenesis using an Agilent QuikChange II XL Site-Directed Mutagenesis Kit (Agilent, 200522) according to manufacturer's instructions to make β-cat$^{S37C/S715A}$ pDONR223. The β-cat$^{S37C/S715A}$ cDNA was then shuttled into pLX304-GW-DCK*-IRES-GFP (Addgene, 176291) destination vector by Gateway LR recombination (Life Technologies, 11791100) as per manufacturer's instructions.

All vectors were confirmed by whole plasmid sequencing (Plasmidsaurus, Primordium, or Genewiz).

*CRISPR/Cas9 Plasmids*

A duplex oligonucleotide encoding an sgRNA for TK1 (sgTK1, see guide sequence section) and with *BsmBI* compatible overhangs was subcloned into a pLentiCRISPRv2 (Addgene, 52961) that was modified to express a hygromycin resistance cassette and then cut with *BsmBI*. The resulting vector expresses both the TK1 sgRNA and Cas9. Duplex oligonucleotides encoding sgRNA targeting AAVS1, DCK, and CTNNB1 (sgAAVS1, sgDCK, and sgCTNNB1_1-4, see guide sequence section) were similarly cloned into pLentiGuide (Addgene, 117986), which expresses the inserted sgRNA but without Cas9. All CRISPR related vectors were confirmed by U6 promoter

Sanger Sequencing (Genewiz) or whole plasmid sequencing (Plasmidsaurus, Primordium, or Genewiz).

*Generation of DOX Inducible PKCδ WT/KD/CA and PKCθ WT/KD/CA Vectors*

The PKCδ *cDNA* in pCAGGS-PKCdelta-EGFP (obtained from Addgene 178856) was PCR amplified (without a stop codon) and with corresponding gateway overhangs (see primers section). The PCR product was gel extracted (Qiagen, 28706), purified, and shuttled into pDONR223 by Gateway BP Clonase Reaction (Thermo Fisher Scientific, 11789020) as per the manufacturer's instructions to yield PKCδ WT(no stop)-pDONR223. PKCθ WT(no stop)-pDONR223 was obtained from Addgene (23424). PKCδ WT(no stop)-pDONR223 or PKCθ WT(no stop)-pDONR223 was subjected to site directed mutagenesis using an Agilent QuikChange II XL Site-Directed Mutagenesis Kit (Agilent, 200522) according to manufacturer's instructions to make: PKCδ KD(kinase dead)(no stop)-pDONR223, PKCδ CA(constitutively active)(no stop)-pDONR223, PKCθ KD(no stop)-pDONR223, PKCθ CA(no stop)-pDONR223. PKCδ KD (K378R) and PKCδ CA (R144A, R145A) mutations were previously described (*2*). PKCθ KD (K409R) and PKCθ CA (R145A, R146A) were designed by homology mapping. The corresponding cDNAs were then shuttled into pINDUCER21-puro_Gateway-3xFLAG (Addgene, 172981) destination vector by Gateway LR recombination (Life Technologies, 11791100) as per manufacturer's instructions.

All vectors were confirmed by whole plasmid sequencing (Plasmidsaurus, Primordium, or Genewiz).

*Primers and Guide Sequences*

GateWay compatible β-cat (no stop) (5'->3'):
FWD: GGGGACAAGTTTGTACAAAAAAGCAGGCTTCatggctactcaagctgatttgatgg
REV: GGGGACCACTTTGTACAAGAAAGCTGGGTCcaggtcagtatcaaaccaggccagc

GateWay compatible primer for 1-664 REV (5'->3'):
REV: GGGGACCACTTTGTACAAGAAAGCTGGGTCctcagacattcggaacaaaacagca

GateWay compatible primer for PKCδ WT (5'->3'):
FWD: GGGGACAAGTTTGTACAAAAAAGCAGGCTTCatggcgccgttcctgcgcatcgcc
REV: GGGGACCACTTTGTACAAGAAAGCTGGGTCatcttccaggaggtgctcgaattt

Site Directed Mutagenesis Single Primer to generate β-cat S37C variants (5'->3'):
5'-phospho gtcttacctggactctggaatccattGtggtgccactaccacagc

QuikChange Primers for S715A β-cat generation (5'->3'):
FWD: gatatcgccaggatgatcctgcctatcgttcttttcactctg
REV: cagagtgaaaagaacgataggcaggatcatcctggcgatatc

QuikChange Primers for PKC KD and CA Mutant Generation (5'->3'):

PKCδ -CA Primer 1: CCTGTTTGATGGCTCCGGCGGCGTTCATCGTTGGGAACT
PKCδ -CA Primer 2: AGTTCCCAACGATGAACGCCGCCGGAGCCATCAAACAGG

PKCδ -KD Primer 1: TCTTGAGGGCCCTGATGGCAAAGTACTCTCCTC
PKCδ -KD Primer 2: GAGGAGAGTACTTTGCCATCAGGGCCCTCAAGA

PKCθ -CA Primer 1: CCTGCTTGATGGCACCCGCGGCCTGATGCAAAGCAAAGA
PKCθ -CA Primer 2: TCTTTGCTTTGCATCAGGCCGCGGGTGCCATCAAGCAGG

PKCθ -KD Primer 1:
CATCTTTCTTTAAGGCCCTTATTGCGAAAAATTGATTGGTTTTCTTGAATT
PKCθ -KD Primer 2:
AATTCAAGAAAACCAATCAATTTTTCGCAATAAGGGCCTTAAAGAAAGATG

sgTK1 (5'->3') TTCACCACGCTCTCGGCCAG
sgAAVS1 (5'->3') GGGGCCACTAGGGACAGGAT
sgDCK (5'->3') TCGAAGGGAACATCGCTGCA
sgCTNNB1_1 (5'->3') AAGGTTATGCAAGGTCCCAG
sgCTNNB1_2 (5'->3') ATACTTACGAAAAACTACTG
sgCTNNB1_3 (5'->3') ATGCAATGACTCGAGCTCAG
sgCTNNB1_4 (5'->3') CAACTGGTAGTCCATAGTGA
sgPRKCA (5'->3') AGGAAGGAAACATGGAACTC

#### *Lentivirus Production*
293FT cells (~1 x $10^6$) were seeded into 60mm cell culture dishes (Corning, 353002) in 4 mL of media. The next morning, the media was removed and replaced with 2.5 mL fresh DMEM supplemented with 10 % FBS (no antibiotic added). Cells were then cotransfected with 1µg of the desired lentiviral vector, psPAX2 (Addgene, 12260; 0.75 µg) and pMD2.G (Addgene, 12259; 0.25 µg) using 6 µL of Lipofectamine 2000 (Invitrogen, 11668019) according to manufacturer's protocol. 24 hours after transfection, the media was removed and 4 mL of fresh media (DMEM supplemented with 30% FBS, no antibiotic) was added. The next day, the lentiviral particle-containing media was collected and replaced with 4 mL of fresh media (DMEM supplemented with 30% FBS, no antibiotic). The following day, this media was again harvested and pooled with the previous day's collection, filtered through a 0.45 µm SFCA filter (Westnet Inc., 229766), aliquoted, and ultimately used (see below) or stored at -80°C.

#### *Lentivirus Infection and Stable Cell Line Generation*
Cells were seeded in 6-well plates (5 x $10^5$ cells in 1 mL per well). Then to each well, 1 mL of lentivirus containing-media was added, followed by treatment with polybrene (final concentration: 0.8 µg/mL, Santa Cruz Biotechnology, SC-134220). The plate was then centrifuged at 252 x g for 30 minutes at 30°C. The lentiviral-containing media was removed 24 hours later, and fresh media was added. The cells were allowed to recover and then selected using the appropriate antibiotic: blasticidin: 10-12.5 µg/mL, puromycin: 2-4 µg/mL, or hygromycin: 200-300 µg/mL.

#### *Fluorescence-Activated Cell Sorting (FACS)*
293FT cells stably expressing a given bicistronic vector with (1) a fusion between DCK* and a POI and (2) GFP were grown to confluency. The cells were then harvested, pelleted (via centrifugation at 300 x g for 3 minutes), and then resuspended at ~2 x $10^6$ cells/mL. 1 mL of this cell suspension was then passed through a mesh strainer into a flow cytometry compatible tube (Thermo Fisher Scientific, 352235). The GFP fluorescence of the population was then assessed using an Aria III SORP or Sony MA900 instrument. The gate for GFP-positive cells was determined by the GFP positivity of other DCK* fusion reporter cells that we found earlier had sufficiently high DCK* levels to render cells sensitive to BVdU. The cells were collected in a 15 mL Eppendorf tube, replated in a 15 cm cell culture dish, and expanded.

***Low Throughput BVdU Assay***

293FT or 293FT TK1 KO cells (with a variety of different reporters expressed) were seeded into 6 well plates (5 x $10^4$ cells in ~2 mL of media per well). BVdU diluted media stocks (3X working concentration) were generated by diluting BVdU (1M in DMSO) into media (9 µL stock per 3 mL of media) and then serial dilution for the remaining desired concentrations of BVdU (1:10 for a total of 5 concentrations in total). The next morning (~12-18 hours later) the media was removed and 2 mL of fresh media + 1 mL of BVdU diluted media stocks were added to each well at the indicated concentrations. In some versions, dThD (100 µM final concentration) was added to the media as an assay false positive. The cells were then incubated for 96 hours and then counted by a ViCell XR (Beckman Coulter, 731050). Data were normalized to a DMSO treated control and results are reported as a "Relative Cell Count". $IC_{50}$ curves and values were calculated using GraphPad Prism 10.4.2 Analysis Tool for log(inhibitor) vs. response -- Variable slope (four parameters).

***CHIR99021 Treated Low Throughput BVdU Assay***

293FT TK1 KO DCK* cells were seeded into 6 well plates (5 x $10^4$ cells in 1 mL of media per well) then 1 mL of either CHIR99021 diluted media stock (50 µL of 10 mM DMSO stock into 25 mL) or DMSO alone. Cells were harvested for immunoblot analysis 24 hours later. The other wells had their media removed and 2 mL of fresh media + 1 mL of BVdU diluted media stocks were added and the rest Low Throughput BVdU assay protocol was completed.

***CRISPR/Cas9 Knockout and BVdU Protection Assay***

293FT TK1 KO DCK* or 293FT TK1 KO β-cat$^{S37C}$-DCK* cells were infected with lentiviral particles for the indicated sgRNAs using the *Lentivirus Infection and Stable Cell Line Generation* protocol. Cells were selected with puromycin (~4 µg/mL) for 7 days post-viral infection. Knockout was confirmed by *Immunoblotting*. Then 293FT TK1 KO DCK* or 293FT TK1 KO β-cat$^{S37C}$-DCK* cells expressing the indicated sgRNAs were seeded in 6 well plates (5 x $10^4$ cells in ~2 mL of media per well). The following morning, media was removed and 2 mL of fresh media then 1 mL of BVdU diluted media stocks (BVdU final concentration of 10 µM for DCK* cells or 50 µM for β-cat$^{S37C}$-DCK* cells) were added to the different wells. The cells were incubated for 96 hours and then counted by a ViCell XR (Beckman Coulter, 731050). Cell counts were normalized to fold change from the corresponding sg*AAVS1* control expressing cells.

***BVdU High Throughput Screening (HTS) Assay***

293FT TK1 KO full length β-cat$^{S37C}$-DCK*, C-terminal truncated $^{1\text{-}664}$β-cat$^{S37C}$-DCK*, full length β-cat$^{S37C/S715A}$-DCK*, DCK*-IKZF1 or DCK* cells were seeded into 384-well plates (Corning, 3764) at 450, 450, 450, 300, or 200 cells/well respectively in 30 µL of media via MultiDrop Combi (Thermo Scientific, 5840330) dispensing. Plates were then centrifuged at 218 x g for 10 seconds and cells were allowed to adhere overnight. The next morning, the plates were treated with test samples (e.g., natural product fractions) via acoustic dosing using an Echo 655 liquid handler. Positive (dThD) (column 2) and negative (DMSO) (column 1) controls were then dosed via Hewlett Packard (HP) D300e (Tecan) dispensing, normalized to the amount of DMSO dispensed during the acoustic dispensing phase. In some experiments, a dose response of different compounds of interest were dosed via HP D300e dispensing at this point. Plates were then centrifuged at 218 x g for 10 seconds and incubated. The next day, 10 µL of BVdU media stock solution was added to each well using a MultiDrop Combi Dispenser (Thermo Scientific, 5840330). The concentration of the BVdU media stock solution was calculated to achieve the desired final BVdU concentration (10 µM for DCK*, 50 µM for β-cat$^{S37C}$-DCK*, 50 µM for C-terminal truncated β-cat$^{S37C}$-DCK*, 100 µM for DCK*-IKZF1). Plates were then centrifuged at 218 x g for 10 seconds and incubated for 96 hours. Note: since there is a significant dilution after BVdU media stock solution addition (30 µL to 40 µL), we chose to always refer to the final concentration at day 2 for a given test sample or control even though from day 1 to day 2 of the assay there is a 1.3X concentration as compared to the final concentration. The GFP fluorescence of each well was then quantified using an Acumen eX3/HCI (STP LabTech) laser scanning cytometer. GFP fluorescence was quantified by defining the metric “GFP+ Object” to identify GFP+ cells while excluding general debris.

***BVdU 384-Well Plate $IC_{50}$ Curves***

Using the standard BVdU HTS assay protocol, plates were treated with either dTHd (133 µM on day 1, 100 µM after BVdU media addition) or DMSO, centrifuged (218 x g for 10 seconds), and incubated for 24 hours. The next day, 10 µL of fresh media was added to every well via MultiDrop Combi (Thermo Scientific, 5840330) dispensing and plates centrifuged. Then the indicated doses of BVdU were added via HP D300 dispenser (Tecan) with a final DMSO percentage of 0.35% per well. Plates were centrifuged at 218 x g for 10 seconds and incubated for 96 hours. GFP Objects were then determined by Acumen reading.

***Z-Prime (Z’) Factor Assessments***

Using the standard BVdU HTS assay, cells were treated with either dThD (100 µM final concentration) or DMSO (100 nL total for test wells) via Echo 655 acoustic dispensing. After 24-hour incubation, plates were treated with corresponding BVdU media stock solution (10 µL added), incubated for 96 hours, and GFP+ objects determined by Acumen scanning. The average and standard deviation for dThd-treated or DMSO-treated wells and a Z’ statistic was calculated.

***Known Bioactive Molecule Screen***

Using the BVdU HTS assay, 2,699 compounds at multiple concentrations were assessed for their ability to selectively protect β-cat$^{S37C}$-DCK* from BVdU activity. Compounds tested were from: Harvard Medical School’s ICCB-Longwood Mechanism of Action library, Harvard Medical School’s LINCS project, an academic collection of small molecules targeting deubiquitinating enzymes (*3*), and other commercial libraries available to screen at ICCB-Longwood. After testing, GFP+ objects for every well were normalized utilizing a Z-score calculation (based on the variation amongst test wells). A hit was defined as any compound (at any dose) with a Z-score ≥ 3 in β-cat$^{S37C}$-DCK* cells and a Z-score < 1 in DCK* cells. A small number of compounds that just missed the hit metric criteria were included.

**AZ-628 Treated Cells for Immunoblotting Analysis**

HEK293T TK1 KO β-cat$^{S37C}$-DCK* cells were seeded into 6-well plates (6 x $10^5$ cells in 2.5 mL per well for 24-hour time point plate or 3 x $10^5$ cells in 2.5 mL per well for 48-hour time point plate). The next morning, 500 µL of AZ-628 containing media at (6X desired final concentration) was added to wells (final DMSO %: 0.25%). Cells were harvested at either 24 hours or 48 hours and pellets were used for *Immunoblotting*.

**WNTinib Treated Cells for Immunoblotting Analysis**

HEK293T TK1 KO β-cat$^{S37C}$-DCK* cells were seeded into 12-well plates (1.5 x $10^5$ cells in 800 µL per well). The next morning, 200 µL of diluted media stocks of WNTinib (5X desired final concentration) were added to wells (final DMSO %: 0.25%). Cells were incubated and harvested after 24 hours and pellets were used for *Immunoblotting*.

***Natural Product HTS***

Using the BVdU HTS assay, 326,304 natural product samples were tested using a Beckmann Echo 655 for dispensing (100 nL of 2.5 mg/mL stock). Extracts and fractions were tested at a final concentration of 6.25 µg/mL. After testing, GFP+ object values were converted to a Z-score (based on the variation of experimental values for a given assay plate). A hit was defined as any fraction with a Z score ≥ 3 in β-cat$^{S37C}$-DCK* cells and a Z score < 1 in DCK* cells. A small number of fractions that just missed the hit metric criteria were included. This yielded 736 hit fractions that were the basis for cherry pick reconfirmation assay (see below).

***Cherry Pick Reconfirmation of Primary HTS Fraction Hits***

Using Echo 655 dispensing, 500 nL of each potential hit fraction was transferred to a 384-well plate (Eppendorf Twin.Tec, 951020552), then using a Bravo (Agilent) 10 µL of DMEM media (no FBS) was added to wells and mixed (5 µL, resuspended up and down 3 times). The plate was incubated at room temperature for 10 minutes. 2 µL of diluted hit fractions in media was added to 293FT TK1 KO β-cat$^{S37C}$-DCK* or DCK* cell plates (30 µL media, standard BVdU HTS Assay conditions) was then added to replicate cell plates using an Agilent Bravo. Plates were then centrifuged (218 x g for 10 seconds) and the remainder of the BVdU HTS Assay was conducted. GFP+ object values were then converted to a $Z^N$-score (based on the variation amongst negative (N) control wells on the assay plate). A hit was defined as any fraction with a $Z^N$-score ≥ 3 in β-cat$^{S37C}$-DCK* cells and a $Z^N$-score < 1 in DCK* cells. A small number of fractions that just missed the hit metric criteria were included.

***Subfraction Library Generation***

Selected hit fractions were subjected to further subfractionation at the NCI Natural Products Branch using fully published methods (*4*). The subfraction samples were plated into 96-well plates at a nominal concentration of ~45 µg/well. To each 96-well plate, 45 µL of DMSO was added using a MultiDrop Combi Dispenser (Thermo Scientific, 5840330). The plates were sealed, agitated on a plate shaker (Thermo Scientific Titer Plate Shaker) for 5 minutes, and spun down (218 x g for 10 seconds). Using a Bravo liquid handler, 10 µL was placed into Echo compatible 384-well plates (Labcyte, LP-0200) and the remaining volume was placed into another 384-well plate (Abgene, 0781) and stored at -20°C.

***Subfraction Secondary Screening***

Using the standard BVdU HTS assay, subfractions were tested at two doses: 100 nL (3.3 µg/mL day 1, 2.5 µg/mL at day 2-6) and 200 nL (6.7 µg/mL day 1, 5.0 µg/mL at day 2-6). GFP+ object values were then converted to a relative GFP+ object value (fold protection) by dividing by to the number of GFP+ objects observed for averaged negative control wells. A hit subfraction was defined as having a fold protection ≥ 3 in β-cat$^{S37C}$-DCK* cells, ≤ 1.2 in DCK*, and ≤ 1.5 in DCK*-IKZF1 at any dose tested.

***Cherry Picking Subfraction Hits***

Using a Tecan Evo75, identified hit subfractions were pipetted into small volume, barcoded FluidX tubes (e.g., Azenta, 68-0701-11). The library of subfraction hits in FluidX tubes were then used for non-HTS follow-up studies.

***Treatment of 293FT TK1 KO β-cat$^{S37C}$-DCK* Cells with Subfractions***

293FT TK1 KO β-cat$^{S37C}$-DCK* or SNU398 cells were seeded into 12-well cell culture treated plates (Thermo Fisher Scientific, 0877229) (1.5 x $10^5$ cells in 1 mL or 0.8 mL of media per well respectively). The following morning, wells were treated with the indicated subfraction at the nominal dose that the subfraction scored previously scored. For L16943 and L15009 subfractions, cells were nominally dosed at 3.3 µg/mL. L90865 subfractions were nominally dosed at 6.7 µg/mL. After 24 hours, media was removed, cells trypsinized, harvested, and pelleted for *Immunoblotting*.

***Proteomics with DLD-1 Cells***

DLD-1 cells were seeded in a 6 well plate (3.5 x $10^5$ cells in 2 mL of media per well). The next morning cells were treated with an impure sample related to L16943 lineage (nominal 10 µM) or DMSO for 6 hours. Cell media was then removed and cells washed with PBS, trypsinized, harvested, and pelleted at 4°C (3200 RPM for 3 minutes). Pellets were then washed 2 times with PBS, and the final pellets were frozen at -80°C. Cells were lysed by addition of lysis buffer (8 M Urea (Fisher Scientific, U-15), 50 mM NaCl, 50 mM 4-(2-hydroxyethyl)-1-piperazineethanesulfonic acid (EPPS, Sigma Aldrich, E9502) pH 8.5, Protease (cOmplete mini-eta free, Sigma Aldrich, 11836170001) and Phosphatase inhibitors (PhosStop, Sigma Aldrich, 4906837001)) and homogenization by bead beating (BioSpec) for three repeats of 30 seconds at 2400 strokes/min. The Bradford assay was used to determine the final protein concentration in the clarified cell lysate. Fifty micrograms of protein for each sample was reduced, alkylated and precipitated using methanol/chloroform as previously described (*5*) and the resulting washed precipitated protein was allowed to air dry. Precipitated protein was resuspended in 4 M urea, 50 mM HEPES (Fisher Scientific, BP3101) pH 7.4, followed by dilution to 1 M urea with the addition of 200 mM EPPS, pH 8. Proteins were digested with the addition of LysC (1:50; enzyme:protein, FUJIFILM Wako, NC9242798) and trypsin (1:50; enzyme:protein, Promega, V5113) for 12 h at 37 °C. Sample digests were acidified with formic acid to a pH of 2-3 before desalting using C18 solid phase extraction plates (SOLA, Thermo Fisher Scientific). Desalted peptides were dried in a vacuum-centrifuged and reconstituted in 0.1% formic acid for liquid chromatography-mass spectrometry analysis.

Data were collected using a TimsTOF Pro2 (Bruker Daltonics, Bremen, Germany) coupled to a nanoElute LC pump (Bruker Daltonics, Bremen, Germany) via a CaptiveSpray nano-electrospray source. Peptides were separated on a reversed-phase $C_{18}$ column (25 cm x 75 µm ID, 1.6 µM, IonOpticks, Australia) containing an integrated captive spray emitter. Peptides were separated using a 50 min gradient of 2 - 30% buffer B (acetonitrile in 0.1% formic acid) with a flow rate of 250 nL/min and column temperature maintained at 50 ºC.

To perform diaPASEF, the precursor distribution in the DDA *m/z*-ion mobility plane was used to design an acquisition scheme for Data-independent acquisition (DIA) data collection that included two windows in each 50 ms diaPASEF scan. Data was acquired using sixteen of these 25 Da precursor double window scans (creating 32 windows) that covered the diagonal scan line for doubly and triply charged precursors, with singly charged precursors able to be excluded by their position in the m/z-ion mobility plane. These precursor isolation windows were defined between 400 - 1200 *m/z* and 1/k0 of 0.7 - 1.3 V.s/cm$^2$.

The diaPASEF raw file processing and controlling peptide and protein level false discovery rates, assembling proteins from peptides, and protein quantification from peptides were performed by searching against a Swissprot human database (January 2021) using library free analysis in DIA-NN 1.8 (*6*). Database search criteria largely followed the default settings for directDIA including: tryptic with two missed cleavages, carbamidomethylation of cysteine, and oxidation of methionine and precursor Q-value (FDR) cut-off of 0.01. Precursor quantification strategy was set to Robust LC (high accuracy) with RT-dependent cross run normalization. Proteins with low sum of abundance (<2,000 x no. of treatments) were excluded from further analysis and the resulting data was filtered to only include proteins that had a minimum of 3 counts in at least 4 replicates of each independent comparison of treatment sample to the DMSO control. Proteins with missing values were imputed according to Baek *et al* (*7*) and protein abundances were scaled and significant changes comparing the relative protein abundance of the treatment to DMSO control comparisons were assessed by two-sided moderated t-test as implemented in the limma package within the R framework.

#### *Immunofluorescence*

DLD-1 cells were seeded in 8-well chambered plates (Ibidi, 80826, 1 x 10$^4$ cells in 300 µL of media per well). The next morning, the cells were dosed with the indicated drug in diluted media stocks (50 µL, final DMSO %: 0.5%). 24 hours later, the media was removed and the cells were washed with DPBS (~300 µL, Gibco, 14190094), then fixed with 4% paraformaldehyde (PFA, Life Technologies, J61899.AK; 30-minute incubation). The cells were then washed 2 times with DPBS and then blocked and permeabilized with a 5% BSA (Gold BioTechnology, A-420-500) and 0.3%TritonX-100 (Sigma Aldrich, 648463) in DPBS solution (2 g BSA, 1.2 mL 10% Triton X-100 in 38.8 mL DPBS). The cells were incubated for at least 1 hour, washed 3 times with DPBS, then incubated with desired primary antibodies (1:500, e.g., anti-β-Cat total Santa Cruz Biotechnology, SC-7963) in 5% BSA in DPBS (2 g BSA in 40 mL DPBS) at 4°C for at least 16 hours. The next morning, the cells were washed 3 times with DPBS (~300 µL) and then incubated with the desired secondary antibody (1:500, e.g., anti-mouse-AlexaFluor Plus 555 (Life technologies, A32727) or anti-rabbit-AlexaFluor Plus 488 (Life Technologies, A32731)) for 2 hours at room temperature. The cells were then washed 2 times with DPBS, incubated with a DAPI in DPBS solution for 5 minutes (1 µL of DAPI solution (Cell Signaling, 4083) in 8 mL of DPBS), washed again 2 times with DPBS, and finally left in 300 µL of DPBS. The wells were then imaged on a Nikon Eclipse Ti2 confocal super-resolution microscope (NIS Elements Version 5.42.01 software). Conventional W1 spinning disk images and SORA enhanced images were taken for wells with a 60X oil immersion objective lens and 405 nm or 561 nm lasers. For each individual experiment, the same exposure times and laser powers was utilized for all wells. Images were then processed using ImageJ (Fuji) and brightness and contrast settings were normalized across the experiment. A single Z plane is shown for images herein even though in some instances a full Z-stack was obtained (20-40 images at 0.25 µm steps).

For the experiments involving DOX-inducible expression of PKC isoforms, the DLD-1 cells were seeded in 8-well chambered plates (Ibidi, 80826, 3 x $10^4$ cells in 250 µL of tetracycline negative media per well). The next morning the cells were treated with doxycycline (DOX, Takara Bio, 631311) for 6 hours (added as a diluted, tetracycline negative-media stock to desired final concentration of 100 ng/µL). Immunofluorescence assays were then performed as described above.

***Treatment of HEK293FT Cells (CHIR99021 or PKC downregulation)***

HEK293FT cells were seeded into a 12-well plate (1.0 x $10^5$ cells in 1000 µL per well). The following morning, the cells were treated with diluted drug stocks in media (500 µL, 3X final concentration, final DMSO %: 0.5%) and incubated for 24 hours. Cells were then harvested and pelleted for *Immunoblotting*.

*PKC Biomarker Experiment with **2**, **3**, ING*

Due to the limited supply of **2** and **3**, several modifications. 293FT cells were seeded into 96-well plates (5 x $10^5$ cells in 300 µL of media per well). After 16 hours, media were removed and incubated with 25 µL DMEM culture media with 2.5 µL compounds prepared in DMSO at 25 µM (**2** or **3**) or 1 µM (ING). After 20 min incubation at 37 °C, media were removed and cells were then suspended with 200 µL cold PBS by pipetting and further washed with another 200 µL cold PBS; an additional 1 mL of cold PBS was added into the tube containing cells. Cells were collected by centrifuge at 10,000 rpm for 15 seconds. PBS were carefully removed without disturbing the cell pellet. Cytoplasmic fraction was prepared using the reported protocol with slightly modification (*8*). For each tube, 900 µL lysate buffer (PBS, 0.1% NP40, Sigma-Aldrich I3021), protease inhibitor cocktail (Roche, 11836170001) and phosphatase inhibitor cocktail (Roche, 4906845001)) were added into the cell pellet. The cell suspension was triturated five times on ice with a P1000 micropipette tip. Addition 200 µL lysate buffer was added into the cell lysate and centrifuged at 13,300 rpm for 1 min. The supernatant was carefully collected without touching the pellet (nuclei) and mixed with 4 mL denature buffer (8M urea, ThermoFisher AM9902, and 2M thiourea, Sigma-Aldrich T7875) to avoid protein degradation and dephosphorylation during the following centrifuge concentration (3,500 rpm, 1.5 hour, room temperature) with filter tube (Millipore UFC801024). The supernatant was collected (~40 µL) and protein centration was determined with protein assay kit (Bio-Rad 5000001). 40 µL supernatant was mixed with 20 µL 4X LDS sample buffer (Invitrogen

NP0008) and boiled for SDS-page. For western blot, 5-8 µg samples were loaded into 4-12% protein gel (Invitrogen XP04122BOX)

***DLD-1 RT-qPCR and RNAseq***

DLD-1 cells were seeded into a 12-well plate (1.2 x $10^5$ cells in 1000 µL per well) and allowed to adhere overnight. The next morning, the media was removed and a diluted media stock with the indicated compounds was added to each well (1000 µL, final DMSO %: 0.7%, compounds 2 and 3 = 25 µM, PMA and ING = 1 µM). After 24 hours, cells were collected by scrapping into DPBS and pelleted (after an additional DPBS wash). Pellets were then homogenized using QIAshredder columns (Qiagen, 79654). Total RNA was extracted from the homogenized lysates using the RNeasy Micro Kit (Qiagen, 74004) following the manufacturer's instructions. RNA concentration was measured using a NanoDrop spectrophotometer (Thermo Scientific, ND-8000-GL). For cDNA synthesis, 1 µg of purified RNA was reverse transcribed in a 20 µL reaction using the AffinityScript qPCR cDNA Synthesis Kit (Agilent, 600559), with an equal mix of oligo dT and random primers. The resulting cDNA was diluted 5-fold with ultra-pure $H_2O$ (Invitrogen, 2997886). qPCR reactions were performed in technical triplicate using 384-well plates (Roche, 047297490001) on a LightCycler 480 Instrument II (Roche). Each 7 µL reaction contained 1.8 µL diluted cDNA, 1 µL nuclease-free water, 0.7 µL primer mix (forward and reverse primers, each at the concentration of 5 µM), and 3.5 µL SYBR Green I Master Mix (Roche 04707516001). Relative expression levels were calculated based on $2^{-\Delta\Delta Ct}$ with *GAPDH* as the reference gene.

Primers used (5'->3'):
*AXIN2* FWD CAACACCAGGCGGAACGAA
*AXIN2* REV GCCCAATAAGGAGTGTAAGGACT
*LEF1* FWD AGAACACCCCGATGACGGA
*LEF1* REV GGCATCATTATGTACCCGGAAT
*LGR5* FWD CTCCCAGGTCTGGTGTGTTG
*LGR5* REV GAGGTCTAGGTAGGAGGTGAAG
*CCND1* FWD GCTGCGAAGTGGAAACCATC
*CCND1* REV CCTCCTTCTGCACACATTTGAA
*cMYC* FWD GCTGCTTAGACGCTGGATTT
*cMYC* REV CACCGAGTCGTAGTCGAGGT
*CTNNB1* FWD AGCTTCCAGACACGCTATCAT
*CTNNB1* REV CGGTACAACGAGCTGTTTCTAC
*GAPDH* FWD GGAGCGAGATCCCTCCAAAAT
*GAPDH* REV GGCTGTTGTCATACTTCTCATGG

*DLD-1 RNAseq Information*

Libraries were prepared using Roche Kapa mRNA HyperPrep strand specific sample preparation kits from 200ng of purified total RNA according to the manufacturer's protocol on a Beckman Coulter Biomek i7. The finished dsDNA libraries were quantified by Qubit fluorometer and Agilent TapeStation 4200. Uniquely dual indexed libraries were pooled in an equimolar ratio and shallowly sequenced on an Illumina MiSeq to further evaluate library quality and pool balance. The final pool was sequenced on an Illumina NovaSeq X Plus targeting 30 million 150bp read pairs per library at the Dana-Farber Cancer Institute Molecular Biology Core Facilities. Sequenced reads were aligned to the UCSC hg38 reference genome assembly and gene counts were quantified using STAR (v2.7.3a) (*9*) and Salmon (*10*). Differential gene expression testing was performed by DESeq2 (v1.22.1) (*11*). RNAseq analysis was performed using the VIPER snakemake pipeline (*12*). To annotate enriched pathways (i.e., MSigDB hallmark terms) (*13*), downregulated genes ($log_2FC < -0.75$, $p_{adj} < 0.01$) were input into Enrichr (*14*) utilizing a defined background of genes (all genes measured in our experiment with a sum of counts > 10).

***LC-MS Analysis of Cell Media Treated with Compound 2***

A stock of **compound 2** diluted in cell media prior to the treatment of DLD-1 cells ($T_0$) was frozen at -80°C. The following day after DLD-1 cell treatment and incubation the media was collected ($T_{24hr}$) and frozen at -80°C. These samples were then analyzed by LC-MS High-resolution mass spectra were recorded on an Agilent 1260 Infinity II UHPLC system coupled to an Agilent 6545 QToF equipped with a dual AJS ESI source. A Kinetex $C_{18}$ column (50.0 × 2.1 mm, 1.7 *μ*m, 100 Å Phenomenex) was used. The mobile phase consisted of $H_2O$ + 0.1% Formic Acid (A) and MeCN + 0.1% Formic Acid (B). The gradient used was maintained at 95:5 (A:B) for 0.5 min, from 95:5 to 0:100 (A:B) for 8.0 min, maintained at 0:100 (A:B) for 0.5 min, from 0:100 to 95:5 (A:B) for 0.5 min and then equilibrated in a post-run at 95:5 (A:B) during 1.0 min. The positive mode ESI conditions were 3 kV of capillary voltage, 1.0 kV of nozzle voltage, gas temperature at 300 ˚C and gas flow of 10 L/min. The data was acquired under the following parameters: positive; acquisition time from 0.5 – 9 min; *m/z* range 100–3200; scan time of 0.5 s.

***Molecular Docking Information/Methods***

Compounds **2** and **3** were prepared by LigPrep (Schrodinger suite 2025-1) with NMR and QM defined stereo chemistry. Other parameters were set as default except pH range to be 7.4 +/- 1. Protein (pdbcode: 7LF3) was prepared prior to docking grid generation. Glide SP was used to dock molecules including the native ligand phorbol. Ring structure sampling option was chosen with input ring structure included, and 50,000 poses per ligand were kept for the initial docking phase with 1000 best poses chosen for energy optimization. 50 poses were optimized post docking, and 10 final best poses were inspected visually. The final pose was further optimized together with the protein by "Refine Protein-ligand complex" protocol in Schrodinger suite. 3D figures were produced by Pymol.

***Microscale Thermophoresis (MST) Binding Assays***

Microscale thermophoresis assay (MST) experiments were carried out on a Nanotemper MonolithNT.115 Pico instrument (Nanotemper Technologies GmbH, Munich, GER) using Monolith His-Tag Labeling Kit (Natotemper Cat# MO-L018) as per the manufacturer's protocol. Recombinant his-tagged Protein Kinase C Alpha (PRKCA) Protein (Cat.# abx068748) and His-tagged Protein Kinase C Delta (PRKCD) Protein (Cat.# abx167998) were purchased from Abbexa LLC. Both proteins were reconstituted in ddH2O at their pre-lyophilized concentrations of 8.37 μM and 4.54 μM for PRKCA and PRKCD, respectively, to preserve their original salt concentration. The stock protein solutions were stored in 4 °C and diluted into assay buffer as needed for each compound binding affinity experiment. The assay buffer used in all MST experiments was 1X PBS, 0.008% Pluoronic acid, pH 8, prepared from a 10X PBS, pH 7.4 stock (Quality Biological) and a 5% stock of pluronic acid (Sigma-Aldrich Catalogue No.: P2443).

The potency of the non-covalent binding between the red-tris NTA dye with the his-tagged PKC proteins was first determined to be sure that it was in a $K_d$ range of 10-50 nM. Here each his-tagged PKC protein was prepped in a 16-point 1:1 serial dilution series (high test of 1000 nM), and to which a red-tris NTA dye solution was added for a final concentration of 25 nM dye per well. All mixings were done in wells of a nonbinding plate (384-well clear PS, Greiner Bio-One cat.no#781901). Following a 30-minute incubation at room temperature, the well solutions were loaded (via capillary action) into individual premium capillaries (Nanotemper Cat# No.: MO-K025) and placed into the detection tray chamber of the Monolith instrument. The experiment protocol was conducted via the MO. Control software (Nanotemper) with the LED/excitation set at 40% and the MST power set at medium. Following a capillary scan to confirm proper fluorescence reading, the MST signal was measured per capillary. Here, the $F_{norm}$ values (normalized fluorescence) were calculated as the ratio of fluorescence after to before the IR laser activation of the visual field of the capillary solution was collected in real-time. The $K_d$ was determined in MO.Control and in MO.Affinity Analysis software (Nanotemper) using the $K_d$ fit module. The resulting binding curves were visualized in Spotfire (TIBCO Software) by plotting the raw $F_{norm}$ values against the $log_{10}$ [dilution series] and using a non-linear logistic regression analysis.

Ahead of compound binding experiments, each his-tagged PKC protein (500 nM) was first labeled at a fixed concentration of red-tris NTA dye (50 nM). After a 30-minute incubation at room temperature, the protein:dye solution was centrifuged at 15,000 g for 10 min at 4 °C to remove any unreacted dye. The supernatant/labeled protein was then transferred to a fresh tube. To uniformly control for DMSO in the DMSO solubilized compounds, each compound dilution (high test in the range of 500-1250 μM) was done in ligand buffer (1X PBS, 0.008% Pluoronic acid, 10% DMSO, pH 8) such that upon diluting (1:1) with the pre-labeled PKC protein, the DMSO concentration in all wells were uniformly at 5%. Following a 30-minute incubation at room temperature, the dilution series of protein:compound solution was loaded as described via capillary action. The MST signal was measured per capillary, with the LED/excitation set to autodetect and the MST power at medium. MO.Affinity and Spotfire were employed for data analysis and visualization, respectively.

***Treatment of HEK293FT TK1KO DCK* Reporter Cell Lines with Compounds and L91313 Subfractions***

HEK293T TK1 KO DCK*, DCK*-IKZF1, full length β-cat$^{S37C}$-DCK*, or C-terminal truncated 1-664 β-cat$^{S37C}$-DCK* cells were seeded into 12-well plates (1.5 x $10^5$ cells in 900 μL per well). The following morning, cells were treated with diluted drug stocks in media (100 μL, 10X final concentration, final DMSO %: 0.5%). Final concentrations: 100 μM dThD, 1 μM pomalidomide, 2.5 μg/mL L91313_6_15, 2.5 μg/mL L91313_6_8. After 24 hours, cells were harvested and pelleted for *Immunoblotting*.

***General Chemistry Experimental Procedures***

Optical rotations were measured on a Rudolph Research Analytical AUTOPOL IV automatic polarimeter with a 0.25 dm pathlength cell in MeOH at 25°C. UV spectra were recorded as methanol solutions on a Varian Cary 50-Bio UV/Vis spectrophotometer. ECD (Electronic Circular Dichroism) experiments were recorded as methanol solutions on a J-1500 CD spectropolarimeter using a quartz cell 1 mm path, at room temperature. FTIR spectra were recorded as thin films on a Bruker Alpha II spectrometer. NMR spectra recorded at 25°C on either Bruker Avance III HD spectrometer, equipped with a 5 mm TCI Cryo-Probe Prodigy or a Bruker Avance III spectrometer equipped with a 3 mm TCI cryogenic probe, both operating at a frequency of 600 MHz for the $^1H$ nucleus and 151 MHz for the $^{13}C$ nucleus. For the 3 mm TCI cryogenic probe, all 2D NMR experiments were acquired with non-uniform sampling (NUS) set to 25% using the standard Bruker pulse sequences. For the 5 mm TCI cryogenic probe, all 2D NMR experiments were acquired with non-uniform sampling (NUS) set to 40% for $^1H$–$^1H$ detected experiments or 35% for $^1H$–$^{13}C$ detected experiments using the standard Bruker pulse sequences. Variable temperature NMR spectra were recorded a Bruker Avance III spectrometer, equipped with a 5 mm cryogenic probe operating at a frequency of 500 MHz for the $^1H$ nucleus and 125 MHz for the $^{13}C$ nucleus. Spectra were calibrated to residual solvent signals at $\delta_H$ 2.50 and $\delta_C$ 39.5 for DMSO-$d_6$. NMR FID processing and data interpretation were done using MestReNova software, version 15.0. Semipreparative scale HPLC purification was performed with a Gilson HPLC purification system equipped with a GX-281 liquid handler, a 322-binary pump, and a 172-photodiode array detector. All solvents used for chromatography, UV, and MS were HPLC grade, and the $H_2O$ was Millipore Milli-Q PF filtered.

High-resolution mass spectra were recorded on an Agilent 1260 Infinity II UHPLC system coupled to an Agilent 6545 QToF equipped with a dual AJS ESI source. A Kinetex $C_{18}$ column (50.0 × 2.1 mm, 1.7 *μ*m, 100 Å Phenomenex) was used. The mobile phase consisted of $H_2O$ + 0.1% FA (A) and MeCN + 0.1% FA (B). The gradient used was maintained at 95:5 (A:B) for 0.5 min, from 95:5 to 0:100 (A:B) for 8.0 min, maintained at 0:100 (A:B) for 0.5 min, from 0:100 to 95:5 (A:B) for 0.5 min and then equilibrated in a post-run at 95:5 (A:B) during 1.0 min. The positive mode ESI conditions were 3 kV of capillary voltage, 1.0 kV of nozzle voltage, gas temperature at 300 ˚C and gas flow of 10 L/min. The data was acquired under the following parameters: positive ionization; acquisition time from 0.5 – 9 min; *m/z* range 100–3200; scan time of 0.5 s.

***Computational Chemistry Methods***

Initial conformational analyses for compounds **2** and **3** were performed in gas phase using Maestro (14.4.133) interface of the Schrodinger software package (Schrodinger, 2023.1) with the OPLS-2005 force field. All conformers with relative energy lower than 5 kcal/mol were saved as a .PDB file and fully optimized using DFT at the B3LYP/DGZVP level using PCM Gaussian 16 software. The optimized stable conformers contributing to greater than 1% were further used. For TDDFT computation of the excited states at CAM-B3LYP/dgtzvp level in MeOH, with consideration of the first 20 excitations. The overall ECD curve of compound **2** and **3** was generated by weighing the contributions of each conformer according to their Boltzmann distribution. All ECD spectra were generated using SpecDis 1.71 software. Calculated $^{13}C$ NMR chemical shift data for **3a** and **3b** were obtained through the calculation isotropic magnetic shielding tensors using the gauge-invariant atomic orbital (GIAO) method at the B3LYP/6-31G(d) level. Geometry optimization, ECD spectra and NMR calculations were also calculated using the PCM solvation algorithm included in the Gaussian 16 program.

***Collection, extraction, and isolation for L15009 lineage that led to compound 1***

The stems of *Calliandra confuse* were collected in Belize in 1988 under the contract through the New York Botanical Gardens for the National Cancer Institute. The plant was taxonomically identified by M. J. Balick and a voucher specimen (Q65S535) was deposited at the Smithsonian Institution, Washington DC. The dried ground stems (796 g) were extracted with MeOH/DCM (1:1, v/v) to yield 15.3 g of the crude organic extract (NSC# N15009). A portion of the crude extract (4.0 g) was prefractionated on a $C_8$ SPE column (40 g) to yield seven fractions: 95:5 $H_2O$:MeOH (L15009_1), 80:20 $H_2O$:MeOH (L15009_2), 60:40 $H_2O$:MeOH (L15009_3), 40:60 $H_2O$:MeOH (L15009_4), 20:80 $H_2O$:MeOH (L15009_5), 0:100 $H_2O$:MeOH (L15009_6), 50:50 MeCN:MeOH (L15009_7). Fraction N15009_5 (322.8 mg) was subjected to repeated preparative HPLC separation on a Phenomenex Kinetex C18 (5 µm, 100 Å, 150 × 21.2 mm) column. Mobile phase was composed of $H_2O$ + 0.1% FA (solvent A) and MeCN + 0.1% FA (solvent B). A linear gradient from 30% to 65% of B was applied over 30 min at a flow rate of 9 mL/min with fraction collection at 1 min increment. HPLC fractions 9-11 (116.2 mg) were further separated on a Phenomenex Kinetex C18 (5 µm, 100 Å, 150 × 21.2 mm) column with a linear gradient of 30% B to 40% B over 30 min at a flow rate of 9 mL/min with fraction collection at 0.5 min increment. Fractions 19-22 yielded **1** (7.8 mg) which were identified to be a mixture of known neolignins, jatrointelignan A and B with NMR spectroscopic data in agreement to those previously reported (*15*); HRESIMS *m/z* 605. 2008 $[M + Na]^+$ (calcd. for $C_{31}H_{34}O_{11}Na^+$ *m/z* 605.1993, Δ = 2.49 ppm).

***Collection, extraction, and isolation for L90865 lineage that led to compounds 2 and 3***

The plant *Euphorbia grantii* was collected in Tanzania in June 1996 by the Missouri Botanical Gardens under contract to the National Cancer Institute. The specimen was taxonomically identified by R. E. Gereau, and a voucher (Q66R1558) was deposited at the Smithsonian Institution, Washington, DC. The air-dried leaves were ground and extracted in DCM/MeOH, 1/1 (v/v), to give 4.61 g of the organic extract (NSC # N90865). A portion of the crude extract (3.28 g) was prefractionated on a $C_8$ SPE column (40 g), generating seven fractions (*16*): 95:5 $H_2O$:MeOH (L90865_1), 80:20 $H_2O$:MeOH (L90865_2), 60:40 $H_2O$:MeOH (L90865_3), 40:60 $H_2O$:MeOH (L90865_4), 20:80 $H_2O$:MeOH (L90865_5), 0:100 $H_2O$:MeOH (L90865_6), 50:50 MeCN:MeOH (L90865_7).

Fraction L90865_5 (88 mg) was further subjected to two HPLC chromatographic separation steps. Initial separation was done using a Kinetex $C_8$ column (21.2 × 150 mm, 5 *μ*m, 100 Å, Phenomenex). Mobile phase was composed of $H_2O$ + 0.1% FA (solvent A) and MeCN + 0.1% FA (solvent B). A gradient method from 50 to 75% of B was applied along 25 min at 9.0 mL/min, and fraction collection was performed in 0.333 min increments starting from time = 1.5 min. HPLC fractions 52-54 (1.5 mg) were combined and further purified using a Luna $C_8$ column (10 × 250 mm, 5 *μ*m, Phenomenex). Mobile phase was composed of $H_2O$ + 0.1% FA (solvent A) and MeCN + 0.1% FA (solvent B). A gradient method from 57 to 65% of B was applied along 30 min at 4.0 mL/min, and fraction collection was performed in 0.333 min increments, starting at time 1.0 min. Fraction 52 yielded **3** (1.2 mg, 0.04% of organic extract yield).

Fraction L90865_4 (30 mg) was subjected to an HPLC separation using a Kinetex $C_8$ column (21.2 × 150 mm, 5 *μ*m, 100 Å, Phenomenex). Mobile phase was composed of $H_2O$ + 0.1% FA (solvent A) and MeCN + 0.1% FA (solvent B). A gradient method from 50 to 75% of B was applied over 25 min at 9.0 mL/min, and fraction collection was performed in 0.333 min increments, starting at time = 1.5 min. Fraction 34 yielded **2** (1.0 mg, 0.03% of organic extract yield).

***Compound 2 (lathyranol)***: $[\alpha]_{\mathrm{D}}^{24}$ -7.6 (*c* 0.1, MeOH); UV (MeOH) $\lambda_{max}$ (log $\varepsilon$) 220 (3.26) nm; ECD ($5.5 \times 10^{-5}$ M, MeOH) $\lambda$ ($\Delta\varepsilon$) 225 (+ 1.94) nm; IR (film) $\nu_{max}$ 3420, 2922, 2851, 1737, 1693, 1594, 1369, 1320, 1239, 1207, 1138, 1098, 1030, 840, 804, 621 $cm^{-1}$; $^{1}H$ and $^{13}C$ data in DMSO-$d_6$, see Table below; HRESIMS *m/z* 633.2671 $[M + Na]^+$ (calcd. for $C_{34}H_{42}O_{10}Na^+$ *m/z* 633.2670, Δ = 0.16 ppm).

***Compound 3 (lathyranol-19-acetate)***: $[\alpha]_{\mathrm{D}}^{24}$ -10.9 (*c* 0.05, MeOH); UV (MeOH) $\lambda_{max}$ (log $\varepsilon$) 222 (3.18) nm; ECD($2.5 \times 10^{-4}$ M, MeOH) $\lambda$ ($\Delta\varepsilon$) 225 (+ 0.96) nm; IR (film) $\nu_{max}$ 2920, 2849, 1736, 1690, 1594, 1415, 1375, 1322, 1239, 1209, 1138, 1094, 1028, 842, 803, 724 $cm^{-1}$; $^{1}H$ and $^{13}C$ data in DMSO-$d_6$, see Table below; HRESIMS *m/z* 675.2776 $[M + Na]^+$ (calcd. for $C_{36}H_{44}O_{11}Na^+$ *m/z* 675.2776, Δ = 0 ppm).

NMR data for **Compounds 3** and **2** in DMSO-$d_6$.

| Compound 3[a] | | | Compound 2[b] | | |
|---|---|---|---|---|---|
| Position | $\delta_C$, type | $\delta_H$, mult (*J* in Hz) | Position | $\delta_C$, type | $\delta_H$ mult (*J* in Hz) |
| 1 | 38.8, $CH_2$ | 2.04, m<br>2.22, dd (14.4, 8.8) | 1 | 39.0, $CH_2$ | 2.04, m<br>2.22, m |
| 2 | 28.5, CH | 2.31, ddt (8.8, 8.4, 7.5) | 2 | 28.5, CH | 2.28, m |
| 3 | 77.2, CH | 5.06, d (8.4) | 3 | 77.5, CH | 5.05, br |
| 4 | 71.2, C | - | 4 | 71.7, C | - |
| 5 | 116.1, CH | 4.83, br s | 5 | 115.7, CH | 4.79, br s |
| 6 | 134.1, C | - | 6 | 134.3, C | - |
| 7 | 78.1, CH | 5.03, br s | 7 | 77.5, CH | 5.03, br |
| 8 | 70.6, CH | 4.57, dd (11.1, 1.3) | 8 | 70.6, CH | 4.58, dd (11.2, 1.2) |
| 9 | 25.8, CH | 1.37, dd (11.1, 9.6) | 9 | 23.0, CH | 1.32, dd (11.2, 9.6) |
| 10 | 24.9, C | - | 10 | 27.6, C | - |
| 11 | 23.1, CH | 1.69, dd (11.5, 9.6) | 11 | 21.4, CH | 1.57, br |
| 12 | 118.9, CH | 4.87, d (11.5) | 12 | 118.6, CH | 4.87, d (10.8) |
| 13 | 127.2, C | - | 13 | 126.9, C | - |
| 14 | 72.4, CH | 5.09, s | 14 | 72.6, CH | 5.08, s |
| 15 | 66.6, C | - | 15 | 67.3, C | - |
| 16 | 16.8, $CH_3$ | 0.79, d (7.5) | 16 | 17.2, $CH_3$ | 0.77, d (6.8) |
| 17 | 14.9, $CH_3$ | 1.71, s | 17 | 15.2, $CH_3$ | 1.68, s |
| 18 | 10.8, $CH_3$ | 0.98, s | 18 | 11.0, $CH_3$ | 0.92, s |
| 19 | 72.1, $CH_2$ | 3.81, d (15.5)<br>3.86, d (15.5) | 19 | 68.7, $CH_2$ | 3.09, dd (11.1,5.9)<br>3.29, dd (11.1, 5.9) |
| 20 | 14.3, $CH_3$ | 1.56, s | 20 | 14.3, $CH_3$ | 1.54, br s |
| 21 | 168.8, C | - | 21 | 169.1, C | - |
| 22 | 20.1, $CH_3$ | 2.06, s | 22 | 20.3, $CH_3$ | 2.05, s |
| 23 | 169.7, C | - | 23 | 169.8, C | - |
| 24 | 20.0, $CH_3$ | 1.96, s | 24 | 22.0, $CH_3$ | 1.95, s |
| 25 | 169.2, C | - | 25 | 170.5, C | - |
| 26 | 40.5, $CH_2$ | 3.74, d (15.5)<br>3.81, d (15.5) | 26 | 40.4, $CH_2$ | 3.72, d (15.5)<br>3.82, d (15.5) |
| 27 | 134.1, C | - | 27 | 134.3, C | - |
| 28/32 | 129.2, CH | 7.32, m | 28/32 | 129.5, CH | 7.31, m |
| 29/31 | 128.1, CH | 7.33, m | 29/31 | 128.3, CH | 7.33, m |
| 30 | 126.6, CH | 7.26, m | 30 | 126.8, CH | 7.27, m |
| 33 | 169.4, C | - | 33 | 170.1, C | - |
| 34 | 20.3, $CH_3$ | 1.99, s | 34 | 20.4, $CH_3$ | 1.98, s |
| 35 | 170.0, C | - | OH | - | 4.65, t (5.9) |
| 36 | 20.3, $CH_3$ | 2.03, s | | | |

[a]NMR data acquired at 60 °C. [b]NMR data acquired at 25 °C

### Structural Elucidation for Compounds 2 and 3

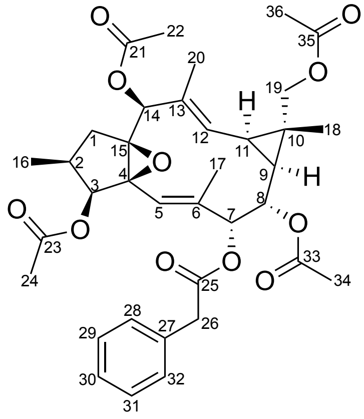

3

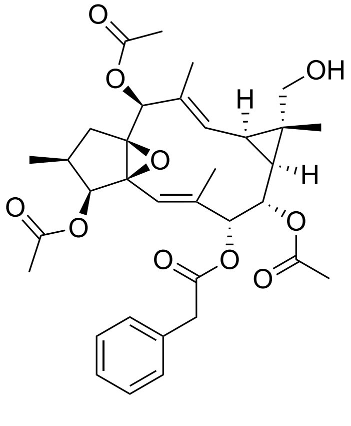

2

Compound **3** was isolated as an optically active clear oil ($[\alpha]_D$ -10.9, *c* 0.05, MeOH) and showed a sodium adduct $[M + Na]^+$ at *m/z* 675.2776 by HRESIMS, corresponding to the molecular formula $C_{36}H_{44}O_{11}$. The $^1$H NMR spectrum of compound **3** in DMSO-$d_6$ showed 24 resonances comprised of five sp$^2$-hybridized methines, seven sp$^3$-hybridized methines, three diastereotopic methylene pairs, and eight methyls. The $^{13}$C NMR spectrum accounted for all 34 carbons in the molecule and showed five carbonyls, five other non-protonated carbons, two sp$^2$-hybridized methine resonances, four oxymethines, three sp$^3$-hybridized methines, three methylenes, two sp$^2$-hybridized methyls, four acetyl methyls, and two other methyls. Interpretation of 2D data allowed the assignment of four acetoxy and one phenylacetate substituents, with the data for the remaining atoms in the molecular formula ($C_{20}H_{25}O$) consistent with a diterpenoid core structure. The 11-membered lathyrane-type macrocyclic ring was established by contiguous $^1$H–$^1$H COSY and $^1$H–$^{13}$C HMBC correlations, and comparison of NMR chemical shifts with other related analogues in the literature (*17-19*). $^1$H–$^{13}$C HMBC correlations from H-3, H-8, and H-14 to carbonyl resonances $\delta_C$ 168.8, 169.4, and 170.0 respectively, implied acetoxy group substitutions at C-3, C-8, and C-14. A deshielded methylene group at $\delta_H$ 3.81/3.86 and $\delta_C$ 72.1 showed $^1$H-$^{13}$C HMBC correlations to resonances associated with the cyclopropyl ring as well as the last remaining acetyl group. This, together with an absence of one methyl resonance in the prototypical lathyrane scaffold suggested acetate group addition on one of the methyl groups in the *gem*-dimethylcyclopropane subunit. At 25°C, the resonance associated with an olefinic methine at $\delta_H$ 4.83, $\delta_C$ 116.1 and an oxymethine at $\delta_H$ 5.03, $\delta_C$ 78.1 were broad and made the correct assignment of the C-5, C-6, and C-7 in compound **3** challenging. Variable-temperature NMR experiment in DMSO-$d_6$ at 25, 30, 40, 50, and 60°C showed sharpening and better resolution of signals at $\delta_H$ 4.83 and 5.03 with increasing temperature (**SI Fig. 15**). At 60°C $^1$H-$^{13}$C HMBC correlations from the sp$^2$-hybridized methine resonance at $\delta_H$ 4.83 ($\delta_C$ 116.1, CH-5) to C-3 and C-4 confirmed the placement of the olefinic methine on C-5 and correlations from H-7 to the carbonyl resonance at $\delta_C$ 169.2 placed the phenylacetate group on C-7 to complete the planar structure of compound **3**. Relative configuration of compound **3** was defined by coupling constant analysis, $^1$H–$^1$H NOESY correlations (**SI Fig. 11, 12**) and comparison of literature values with that of related compounds with defined absolute configurations. The *E* geometry of the $\Delta^5$ and $\Delta^{12}$ double bonds was assigned based on lack of NOESY NMR spectrum cross peaks between H-5 to H-17, and H-12 to H-20, while the observed $^1$H-$^1$H NOESY correlations from H-5 to H-3, H-17 to H-7 and H-14 to H-20 corroborated the assignment. The cyclopentane ring $^1$H–$^1$H NOESY correlations between H-1$\alpha$, H-2, H-3, H-15, and H-14 showed them to be co-facial and arbitrarily assigned as $\alpha$-oriented, while correlations from H-1$\beta$ and H-16 and positioned them on the $\beta$ face of the molecule. The *cis*-configuration of H-2 and H-3 was further supported by the measured $^{3}J_{HH}$ coupling constant of 8.4 Hz. The oxymethine at H-8 was observed as a doublet of doublets ($J$ = 11.1, 1.3 Hz) with a $^{3}J_{HH}$ coupling constant to H-7 of 1.3 Hz suggesting a *syn*-orientation between H-7 and H-8 and a $^{3}J_{HH}$ coupling constant to H-9 of 11.1 Hz suggesting the *anti*-orientation between H-7 and H-9, in agreement with that observed for other ingol analogues whose structures were confirmed *via* XRD (*17*). Protons at H-9, H-11, and H-19 on the cyclopropane ring showed NOESY cross-peaks and were assigned as *α*-oriented, with the *cis*-configuration of H-9 and H-11 further supported by the measured $^{3}J_{HH}$ coupling constant of 9.6 Hz. Protons at H-8, H-12 and H-18 were assigned as *β*-oriented due to their NOESY cross-peaks. $^{13}$C NMR chemical shifts were calculated for **3** to define the relative configuration of non-protonated carbons at C-4 and C-15 as either 4*S**,15*S** (isomer **3a**) or 4*R**,15*R** (isomer **3b**) (**SI Fig. 37-39**). Optimized conformers were used to calculate the magnetic shielding tensors using gauge-invariant atomic orbital (GIAO) method at B3LYP/6-31G(d) level. The data for both isomers were scaled using empirical factors for B3LYP hybrid functional and 6-31G(d) basis set in DMSO. Comparison between experimental data for **3** with that calculated for isomers **3a** and **3b** were compared by four different methods, namely: DP4+ (*20*), mean absolute error (MAE), mean squared error (MSE) and root mean square error (RMSE). For all the statistical parameters (*21*), isomer **3a** (4*S**,15*S**) showed to be the best fit for compound **3**. Together, these data defined the relative configuration of compound **3** to be (2*S**,3*S**,4*S**,7*R**,8*S**,7*R**,8*S**,9*S**,10*S**,11*R**,14*S**,15*S**).

Comparison between $^{13}C$ NMR chemical shifts for the experimental data of the naturally occurring **3** and GIAO DFT calculated isomers **3a** and **3b**.

| Position | **3** $\delta_{Exp}$ (ppm | **3a** $\delta_{Calc}$ (ppm) | **3b** $\delta_{Calc}$ (ppm) |
|---|---|---|---|
| 1 | 38.8 | 37.3 | 35.9 |
| 2 | 28.5 | 30.3 | 32.8 |
| 3 | 77.2 | 79.8 | 79.9 |
| 4 | 71.2 | 73.2 | 69.2 |
| 5 | 116.1 | 118.4 | 116.4 |
| 6 | 134.1 | 136.0 | 137.7 |
| 7 | 78.1 | 81.9 | 77.1 |
| 8 | 70.6 | 73.1 | 74.7 |
| 9 | 25.8 | 28.3 | 26.8 |
| 10 | 24.9 | 26.5 | 27.0 |
| 11 | 23.1 | 24.8 | 25.5 |
| 12 | 118.9 | 121.9 | 122.0 |
| 13 | 127.2 | 127.0 | 131.8 |
| 14 | 72.4 | 75.4 | 76.6 |
| 15 | 66.6 | 70.3 | 71.9 |
| 16 | 16.8 | 15.7 | 10.9 |
| 17 | 14.9 | 13.2 | 12.6 |
| 18 | 10.8 | 10.0 | 10.4 |
| 19 | 72.1 | 75.8 | 75.1 |
| 20 | 14.3 | 11.9 | 14.6 |
| 21 | 168.8 | 168.6 | 168.8 |
| 22 | 20.1 | 17.5 | 18.2 |
| 23 | 169.7 | 169.8 | 169.2 |
| 24 | 20.0 | 17.4 | 18.2 |
| 25 | 169.2 | 170.1 | 169.8 |
| 26 | 40.5 | 39.9 | 41.8 |
| 27 | 134.1 | 131.2 | 131.6 |
| 28 | 129.2 | 126.1 | 125.3 |
| 29 | 128.1 | 124.9 | 125.2 |
| 30 | 126.6 | 123.7 | 123.7 |
| 31 | 128.1 | 125.0 | 125.1 |
| 32 | 129.2 | 126.4 | 125.7 |
| 33 | 169.4 | 169.3 | 169.0 |
| 34 | 20.3 | 17.9 | 17.8 |
| 35 | 170.0 | 169.9 | 169.6 |
| 36 | 20.3 | 17.6 | 18.0 |
| DP4+ | | 100% | 0% |
| $R^2$ | | 0.9982 | 0.9974 |
| MAE | | 2.06 | 2.38 |
| MSE | | 5.41 | 7.90 |
| MAPE | | 4.96 | 5.60 |
| RMSE | | 2.33 | 2.81 |

Compound **2** showed a sodium adduct $[M + Na]^+$ at *m/z* 675.2776 by HRESIMS, corresponding to the molecular formula $C_{36}H_{44}O_{11}$. The $^1H$ NMR spectrum of compound **2** was similar to that of compound **3**, with the exception of loss of resonances for one acetyl group and a presence of a new exchangeable resonance at $\delta_H$ 4.65. The exchangeable resonance showed a $^1H$-$^1H$ COSY and a $^1H$-$^{13}C$ HMBC correlation to a diastereotopic methylene pair at $\delta_H$ 3.09/3.29 and $\delta_C$ 71.7, as well as an additional $^1H$-$^{13}C$ HMBC correlation to a non-protonated carbon at C-10, suggesting loss of the acetyl substitution at C-19. The placement of the primary alcohol on C-19 was further confirmed by $^1H$-$^{13}C$ HMBC correlations from H-19 to C-8, C-9, and C-10. ROESY NMR spectrum cross-peaks between H-1α, H-2, H-3, H-15, and H-14 ad H-9, H-11, and H-19 showed them to be co-facial and arbitrarily assigned as *α*-oriented, while correlations from H-1*β* and H-16 and from H-7 to H-9 positioned them on the *β* face of the molecule. The absolute configuration of compounds **2** and **3** was established *via* comparison of the calculated and experimental ECD spectra using time-dependent density functional theory (TDDFT) at the CAM-B3LYP/dgdzvp level. The experimental ECD spectrum of **2** and **3** in MeOH showed similar trends and magnitude ratios with that of the calculated spectrum for the (2*S*,3*S*,4*S*,7*R*,8*S*,7*R*,8*S*,9*S*,10*S*,11*R*,14*S*,15*S*) enantiomers (**SI Fig. 36**) to complete the stereochemical assignment of compounds **2** and **3** as depicted.

### Supplementary Figures

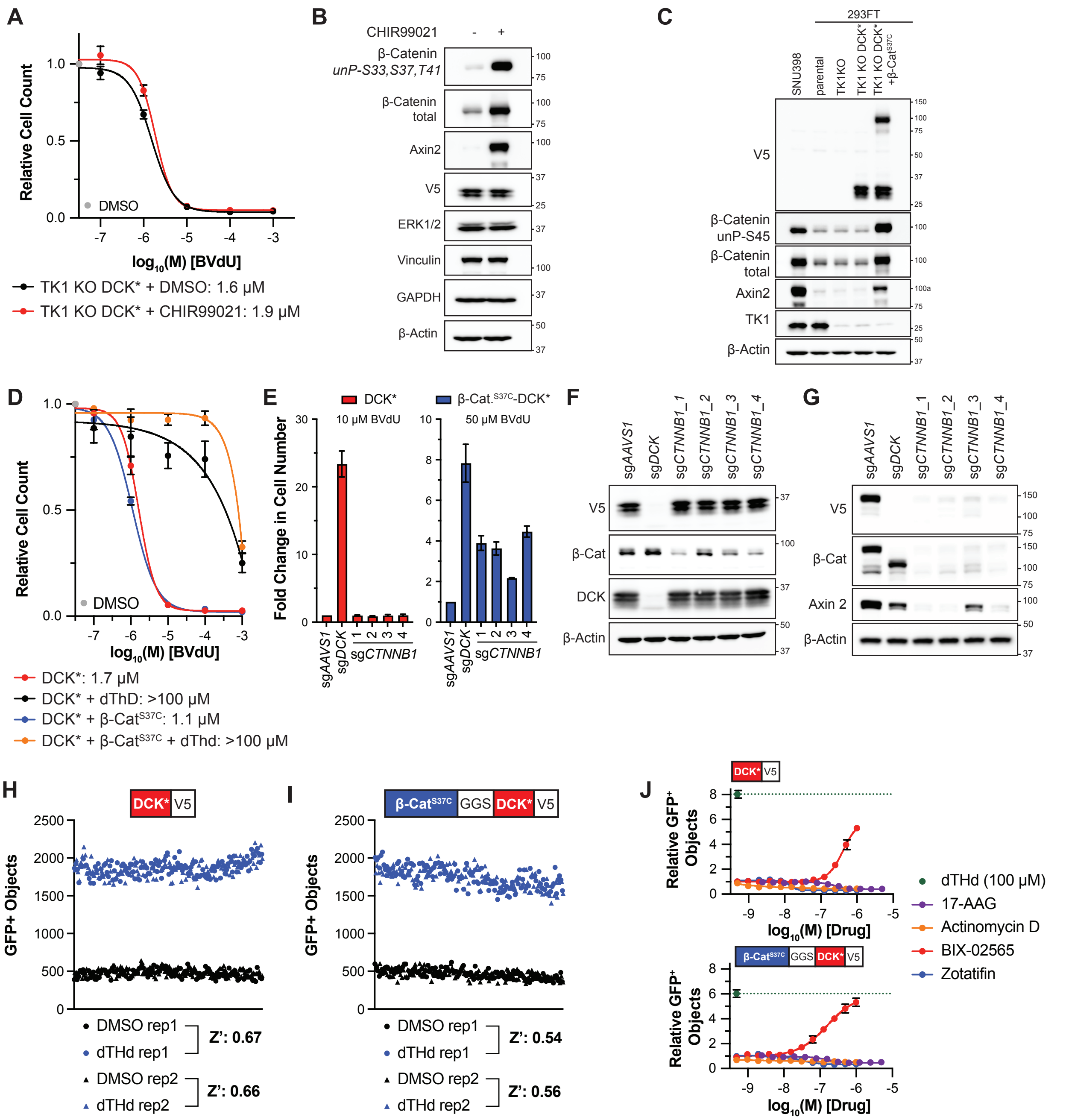


**SI Figure 1. BVdU Sensitivity is Not Affected by β-Cat Activity and β-Cat$^{S37C}$-DCK* System Performance in 384-Well Format.**
(**A**) Relative survival of 293FT TK1 KO DCK*cells treated with CHIR99021 (10 μM) or DMSO for 24 hours followed by indicated BVdU concentrations for 96 hours. n = 3 biological replicates. (**B**) Immunoblot analysis of cells treated as in (**A**) after the initial 24 hours. (**C**) Immunoblot analysis of 293FT TK1 KO cells expressing, where indicated, V5-tagged versions of DCK*, oncogenic β-Cat$^{S37C}$, or both. SNU398 hepatocellular carcinoma cells, which harbor an endogenous β-Cat$^{S37C}$ mutation and parental 293FT cells were included as controls. (**D**) Relative survival of 293FT cells as in (**C**) after treatment with the indicated BVdU concentrations for 96 hours. n = 3 biological replicates. (**E**) Fold change in cell number of 293FT TK1 KO

DCK* cells or 293FT TK1 KO β-Cat$^{S37C}$-DCK* cells stably expressing Cas9 and the indicated CRISPR sgRNAs and then treated with BVdU (10 µM for DCK* cells and 50 µM for β-Cat$^{S37C}$-DCK* cells) for 96 hours. n = 3 biological replicates. *AAVS1* targeting guide (sg*AAVS1*) is the negative control and data is set relative to sg*AAVS1* control cells. (**F**,**G**) Immunoblot analysis of 293FT TK1 KO DCK* cells (**F**) and 293FT TK1 KO β-Cat$^{S37C}$-DCK* (**G**) stably expressing Cas9 and the indicated CRISPR sgRNAs. (**H**,**I**) Z-prime calculation and cell survival (GFP+ objects) analysis of 293FT TK1 KO DCK* (**H**) or 293FT TK1 KO β-Cat$^{S37C}$-DCK* (**I**) that were grown in 384-well plates, treated with DMSO or dThD (final concentration: 100 µM), and then BVdU for 96 hours (10 µM for DCK* cells and 50 µM for β-Cat$^{S37C}$-DCK* cells). n = 2 technical replicates. (**J**) Relative GFP+ objects for 293FT TK1 KO DCK* or β-Cat$^{S37C}$-DCK* cells treated with the indicated compounds and concentrations for 24 hours, followed by BVdU (10 µM for DCK* cells and 50 µM for β-Cat$^{S37C}$-DCK* cells) for 96 hours. n = 3 technical replicates.

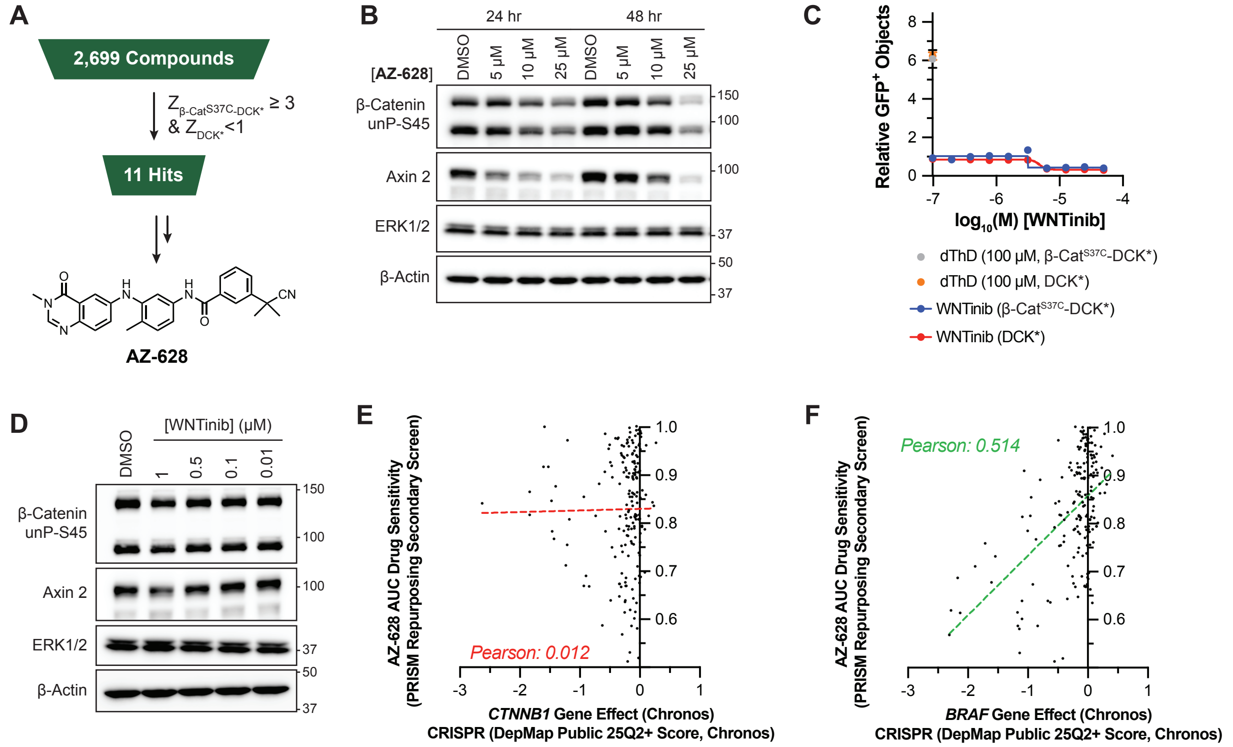


**SI Figure 2. β-Cat$^{S37C}$-DCK* Positive Selection Screening of Known Bioactive Compounds Identified AZ-628.**

(**A**) Summary of known bioactive small molecule screen that identified AZ-628. (**B**) Immunoblot analysis of 293FT TK1 KO β-Cat$^{S37C}$-DCK* cells treated with AZ-628 at the indicated concentrations for 24 or 48 hrs. n = 3 biological replicates. (**C**) Relative GFP+ objects for 293FT TK1 KO DCK* or β-Cat$^{S37C}$-DCK* cells treated with the indicated concentrations of WNTinib for 24 hours followed by BVdU for 96 hours. n = 2 for technical replicates. (**D**) Immunoblot analysis of 293FT TK1 KO β-Cat$^{S37C}$-DCK* cells treated with WNTinib at the indicated concentrations for 24 hours. n = 2 biological replicates. (**E**,**F**) Correlations between sensitivity (Area Under the Curve, AUC) of cells lines to AZ-628 with *CTNNB1* dependence (**E**) or *BRAF* dependence (**F**) based on publicly available data in DepMap (DepMap Public 25Q2 Chronos and PRISM Repurposing Secondary Screen). The Pearson correlation is displayed with a poor correlation in red and a good correlation in green.

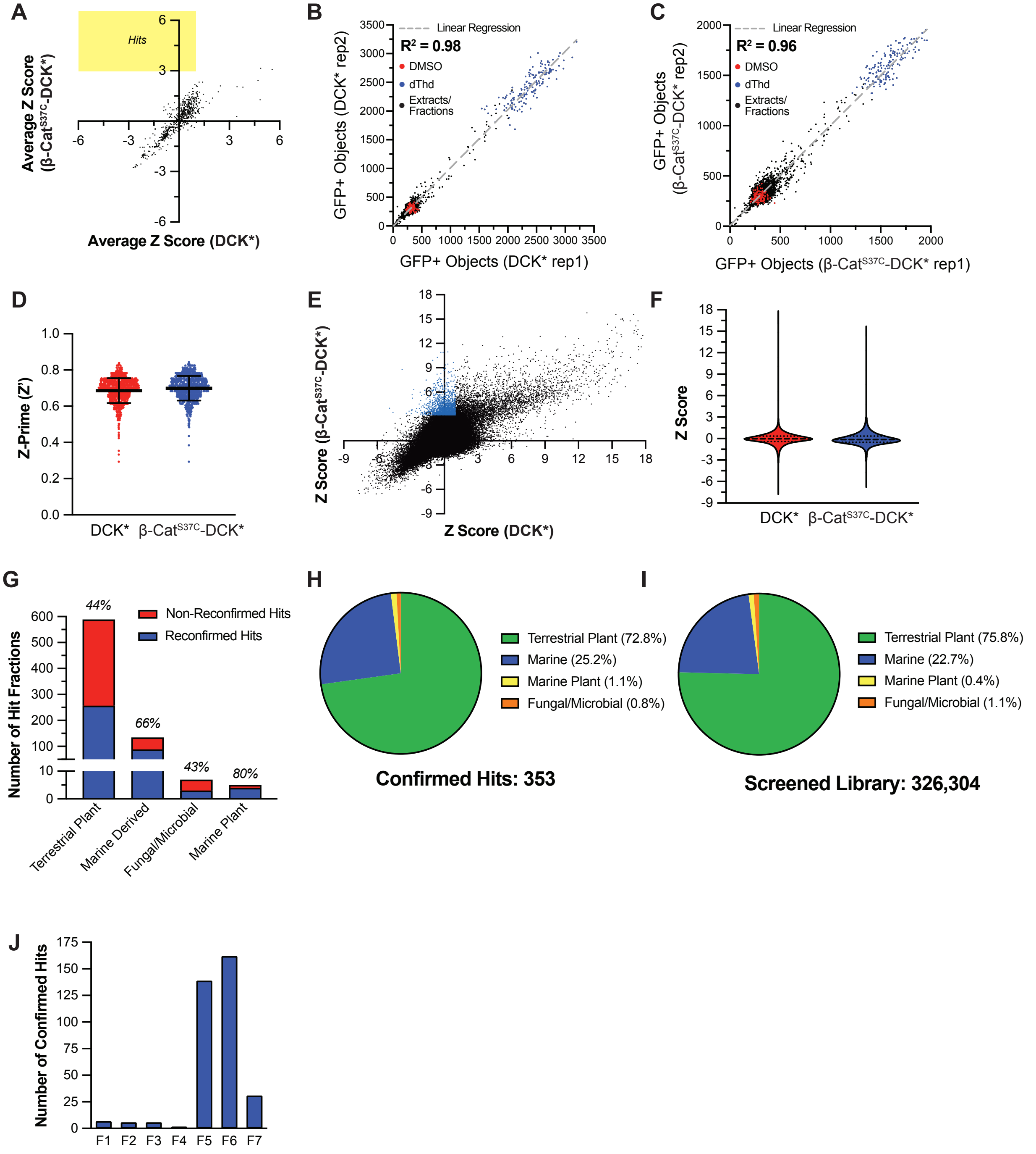


**SI Figure 3. Positive Selection β-Cat$^{S37C}$-DCK* Screening of Prefractionated Natural Product Library.** (**A**) Average Z-scores observed when testing "challenger" natural product plates. n = 2 technical replicates. Hits (yellow box) are defined as Z ≥ 3 in 293 FT TK1 KO β-Cat$^{S37C}$-DCK* cells and Z < 1 in 293FT TK1 KO DCK* cells. (**B**,**C**) Replicate to replicate correlations for 293FT TK1 KO DCK* cells (**B**) and 293FT TK1 KO β-Cat$^{S37C}$-DCK* cells (**C**) when treated with either controls (DMSO in red or dThD in blue) or natural product extracts/fractions (in black) followed by BVdU for 96 hours. The linear regression line is plotted in gray and the correlation's $R^2$ is displayed ($R^2 > 0.95$ is considered optimal). (**D**) Z-Prime (Z') factors calculated during the primary screen of natural product extracts and fractions. Z' > 0.3 is considered acceptable for screening. (**E**) Z-scores for all natural product extracts and fractions tested. Hits are identified in blue (Z ≥ 3 in β-

Cat$^{S37C}$-DCK* cells and Z < 1 in DCK* cells). (**F**) Violin plots displaying the Z-scores for the natural product primary screen. (**G**) Graph showing total number of hits and cherry pick reconfirmation likelihood as a function of hit source. (**H**,**I**) Pie chart of the various sources for confirmed hits in the screen (**H**) compared to the screened library (**I**). (**J**) Histogram of confirmed hits and their corresponding fraction number. Fractions were purified utilizing a reverse phase approach, thus increasing fraction number is correlated with increasing lipophilicity.

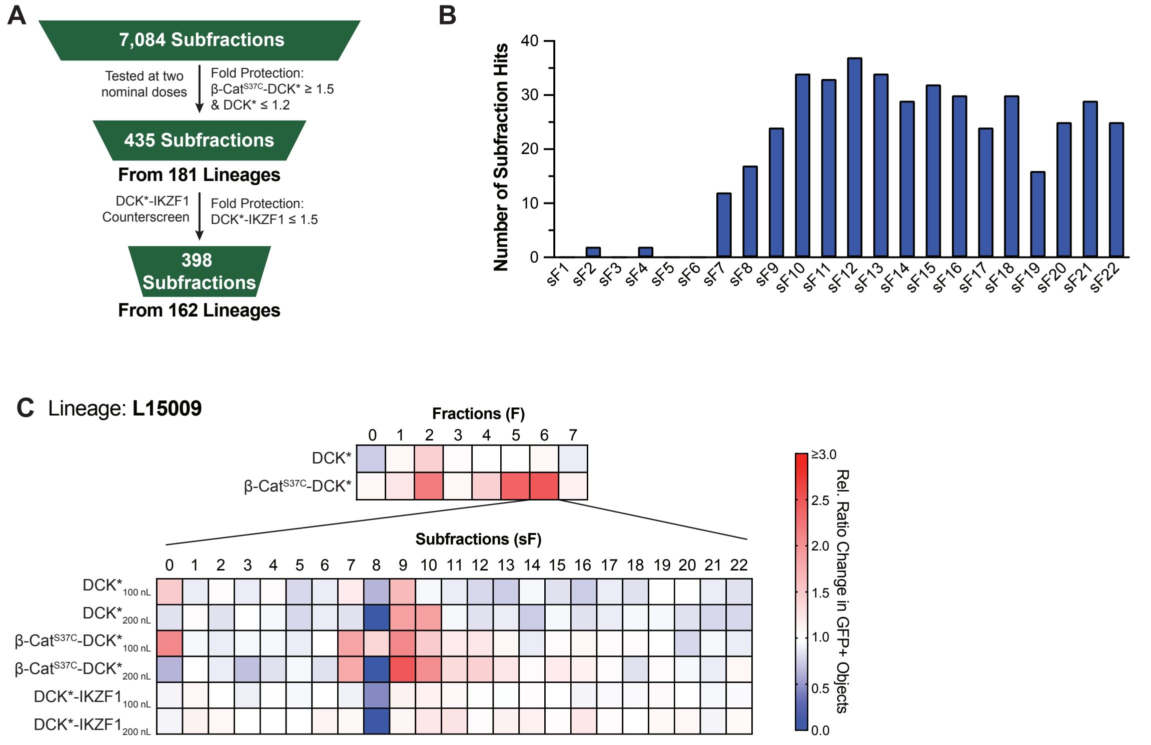


**SI Figure 4. Summary of Subfraction Screening Results.**
(**A**) Summary of subfraction screening, counterscreening, hit metrics, and the number of subfraction hits and lineages. Fold protection: relative increase in GFP+ objects as compared to negative control. (**B**) Histogram depicting the number of hits for each subfraction number. Subfractions were generated utilizing a reverse-phase HPLC step, so increasing subfraction number is correlated with increasing lipophilicity. (**C**) Heatmap showing primary and secondary screening data, based on cell survival (GFP+ objects), related to lineage L15009 (fraction L15009_6). Coloring is representative of the relative GFP+ objects measured in a given test well. 100 nL and 200 nL refers to the transfer volume for subfraction dosing. n = 2 technical replicates for subfraction data and the data are shown as an average.

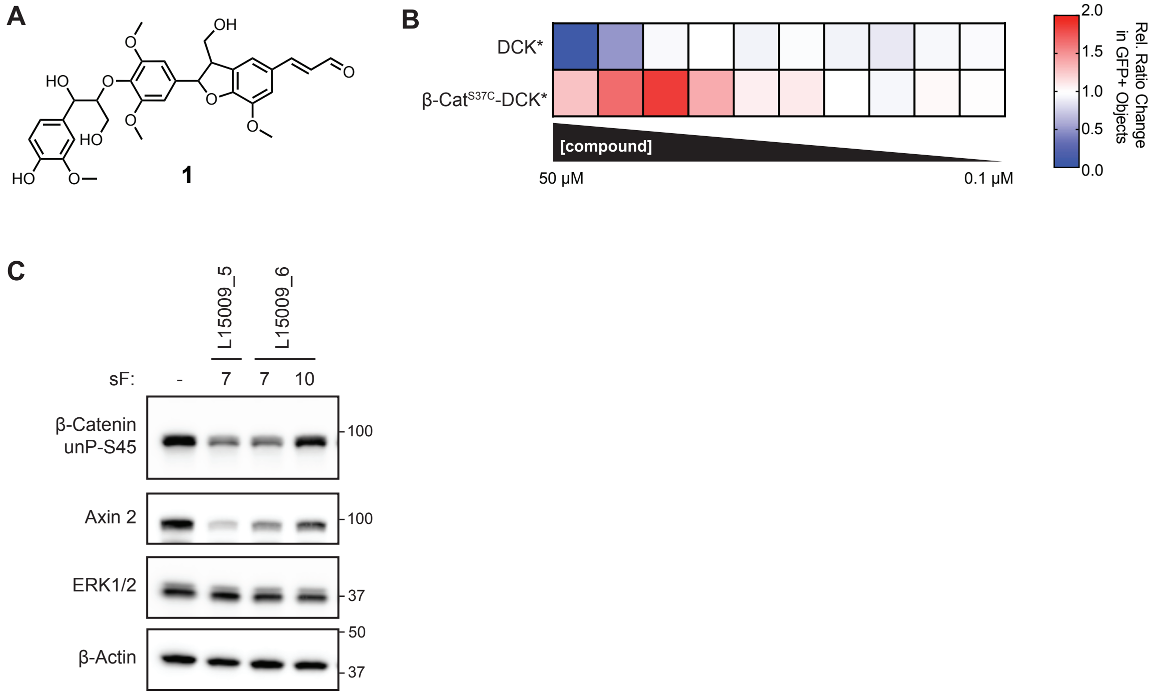


**SI Figure 5. Compound 1 is the Active Principle Compound for L15009 Downregulator Lineage.** (**A**) Chemical structure of compound **1**, a neolignan natural product. (**B**) Heatmap of relative increase in GFP+ objects for 293FT TK1 KO DCK* and 293FT TK1 KO β-Cat$^{S37C}$-DCK* cells utilizing the BVdU assay. n = 3 technical replicates and shown as an average. (**C**) Immunoblot analysis of SNU398 β-Cat$^{S37C}$ mutant hepatocellular carcinoma cells treated with L15009 subfractions for 24 hours. n = 2 technical replicates given limiting material.

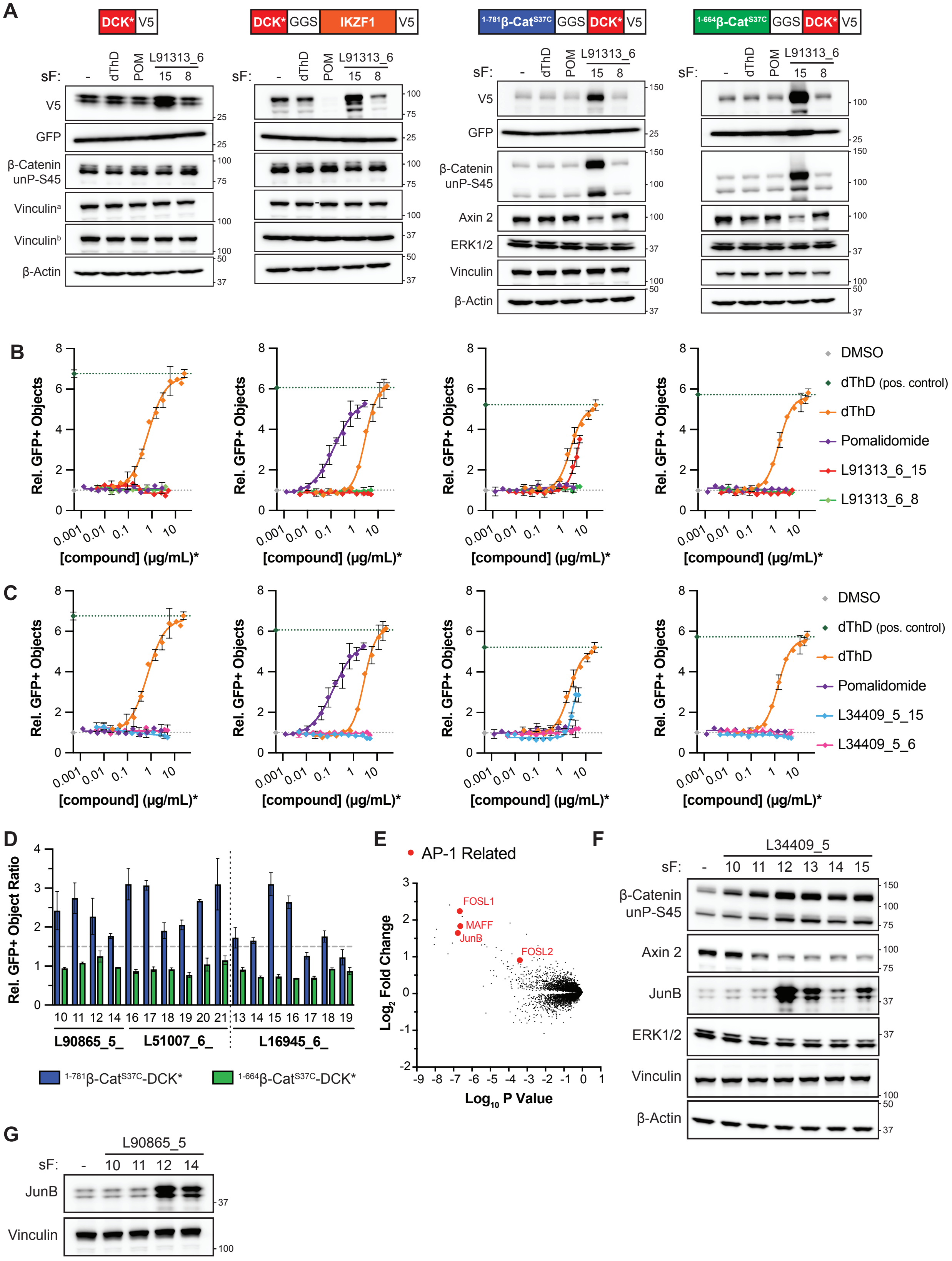
A
DCK* V5
DCK* GGS IKZF1 V5
1-781β-CatS37C GGS DCK* V5
1-664β-CatS37C GGS DCK* V5
sF: - dThD POM L91313_6 15 8
V5
GFP
β-Catenin unP-S45
Vinculin^a
Vinculin^b
β-Actin
Axin 2
ERK1/2
Vinculin
B
Rel. GFP+ Objects
[compound] (μg/mL)*
DMSO
dThD (pos. control)
dThD
Pomalidomide
L91313_6_15
L91313_6_8
C
L34409_5_15
L34409_5_6
D
Rel. GFP+ Object Ratio
10 11 12 14 16 17 18 19 20 21 13 14 15 16 17 18 19
L90865_5_
L51007_6_
L16945_6_
1-781β-CatS37C-DCK*
1-664β-CatS37C-DCK*
E
AP-1 Related
FOSL1
MAFF
JunB
FOSL2
Log2 Fold Change
Log10 P Value
F
L34409_5
sF: - 10 11 12 13 14 15
β-Catenin unP-S45
Axin 2
JunB
ERK1/2
Vinculin
β-Actin
G
L90865_5
sF: - 10 11 12 14
JunB
Vinculin

**SI Figure 6. Inactivator Subfractions Specifically Protect Full-Length β-Cat$^{S37C}$-DCK* cells from BVdU.**

(**A**) Immunoblot analysis of the indicated 293FT TK1 KO DCK* reporter cells treated with DMSO, dThD (assay positive, 100 μM), POM (pomalidomide, IKZF1 degrader, 1 μM), active inactivator subfraction (L91313_6_15, 2.5 μg/mL), or inactive inactivator subfraction (L91313_6_8, 2.5 μg/mL) for 24 hours. Vinculin$^{a}$: loading control for GFP blot. Vinculin$^{b}$: loading control for V5 blot. (**B**,**C**) Relative GFP+ objects for the indicated 293FT TK1 KO DCK* reporter cells treated with the indicated compounds/subfractions over the concentration range shown. Cells were treated with compounds/subfractions for 24 hours, then with BVdU for 96 hours (10 μM of DCK* cells left most panel, 100 μM for DCK*-IKZF1, 50 μM for both full-length 1-781 β-Cat$^{S37C}$-DCK* and C-terminal truncated 1-664 β-Cat$^{S37C}$-DCK* cells). *: For subfractions, the concentration (μg/mL) is nominal. n = 3 technical replicates. (**D**) Relative GFP+ objects for 293FT TK1 KO cells expressing full-length (1-781) β-Cat$^{S37C}$-DCK* or C-terminal truncated 1-664 β-Cat$^{S37C}$-DCK* cells treated with inactivator subfractions for 24 hours (5 μg/mL) followed by BVdU (50 μM) for 96 hours. n = 2 for technical replicates. (**E**) Proteomic analysis of DLD-1 cells treated with an impure sample derived from L16943 (nominal concentration of 10 μM) for 6 hours. AP-1 transcription factors that were among the top 40 proteins with an increased abundance (relative to DMSO control) are highlighted in red. n = 3 biological replicates. (**F**) Immunoblot analysis of 293FT TK1 KO β-Cat$^{S37C}$-DCK* cells treated with inactivator subfraction hits from L34409_5 (3.3 μg/mL) for 24 hours. (**G**) Immunoblot analysis of β-Cat$^{S37C}$-DCK* cells treated with inactivator subfraction hits from L90865_5 (6.7 μg/mL) for 24 hours. The samples analyzed in (**G**) were also immunoblotted to produce **Fig. 2F**. "-" = DMSO treatment.

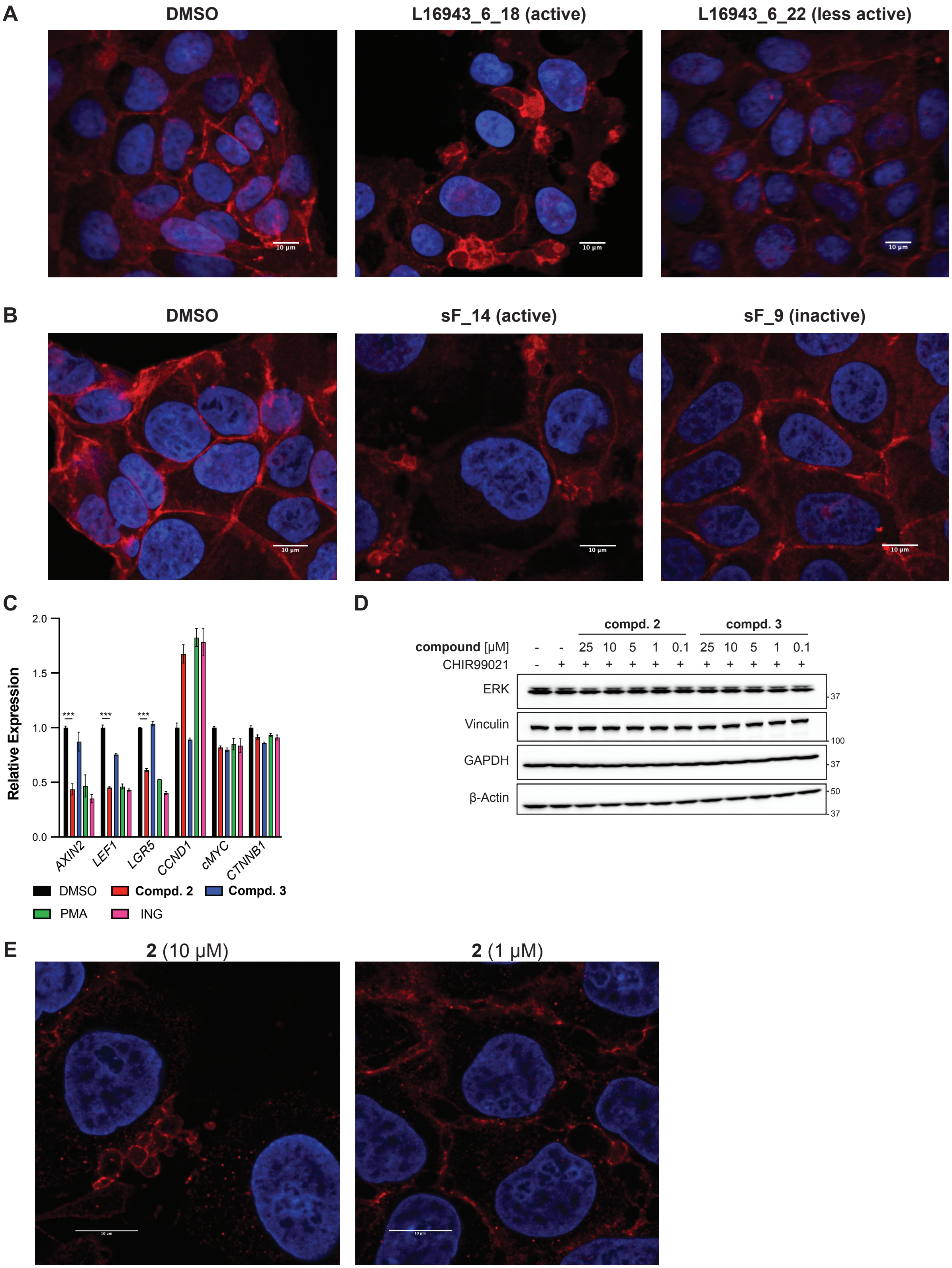
A
DMSO
L16943_6_18 (active)
L16943_6_22 (less active)
B
DMSO
sF_14 (active)
sF_9 (inactive)
C
Relative Expression
2.0
1.5
1.0
0.5
0.0
***
***
***
AXIN2
LEF1
LGR5
CCND1
cMYC
CTNNB1
DMSO
Compd. 2
Compd. 3
PMA
ING
D
compd. 2
compd. 3
compound [µM] - - 25 10 5 1 0.1 25 10 5 1 0.1
CHIR99021 - + + + + + + + + + + +
ERK
Vinculin
GAPDH
β-Actin
37
100
37
50
37
E
2 (10 µM)
2 (1 µM)

**SI Figure 7 (cont.)**

**F**

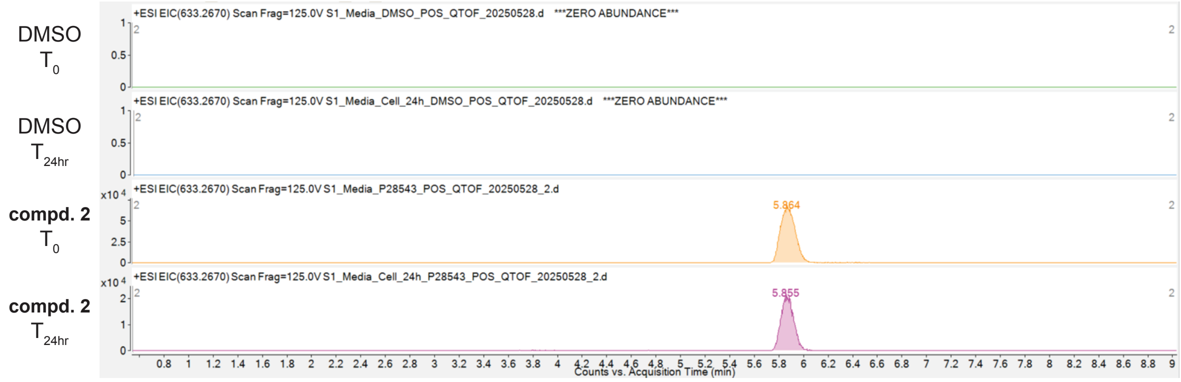


**SI Figure 7. Multiple β-Cat Inactivator Lineages Contain Compounds that Alter β-Cat Localization in APC mutant DLD-1 Colorectal Cancer Cells.**
(**A,B**) Immunofluorescence images of DLD-1 cells treated with the indicated the **L16943** subfractions (3.3 µg/mL) or DMSO for 24 hours. For (**B**), a semi-purified derivative of a L16945 lineage subfraction was used (sF_14) compared to inactive derivative in the same series (sF_9). Blue: DAPI stain, Red: β-Cat. Scale bar = 10 µm. (**C**) Relative mRNA expression of the indicated genes as measured by RT-qPCR of DLD-1 cells after 24-hour incubation with **compound 2** (25 µM), **compound 3** (25 µM), PMA (1 µM), ING (1 µM), or DMSO. n = 3 technical replicates, ***: $p < 0.001$ (Student unpaired T-Test). (**D**) Further immunoblot loading controls associated with **Fig. 3C**. (**E**) Immunofluorescence images of cells treated with compound **2** at the indicated concentrations for 24 hours. The data in (**E**) were generated as part of the experiment that produced the data depicted in **Fig. 3D,E** and shows the full dose-dependency of **2**. Blue: DAPI stain, Red: β-Cat. Scale bar = 10 µm. (**F**) LC-MS analysis of compound **2** after incubation with DLD-1 conditioned cell culture media at time 0 ($T_0$) or after 24 hours ($T_{24hr}$) with. Note: **P28543** = compound **2**.

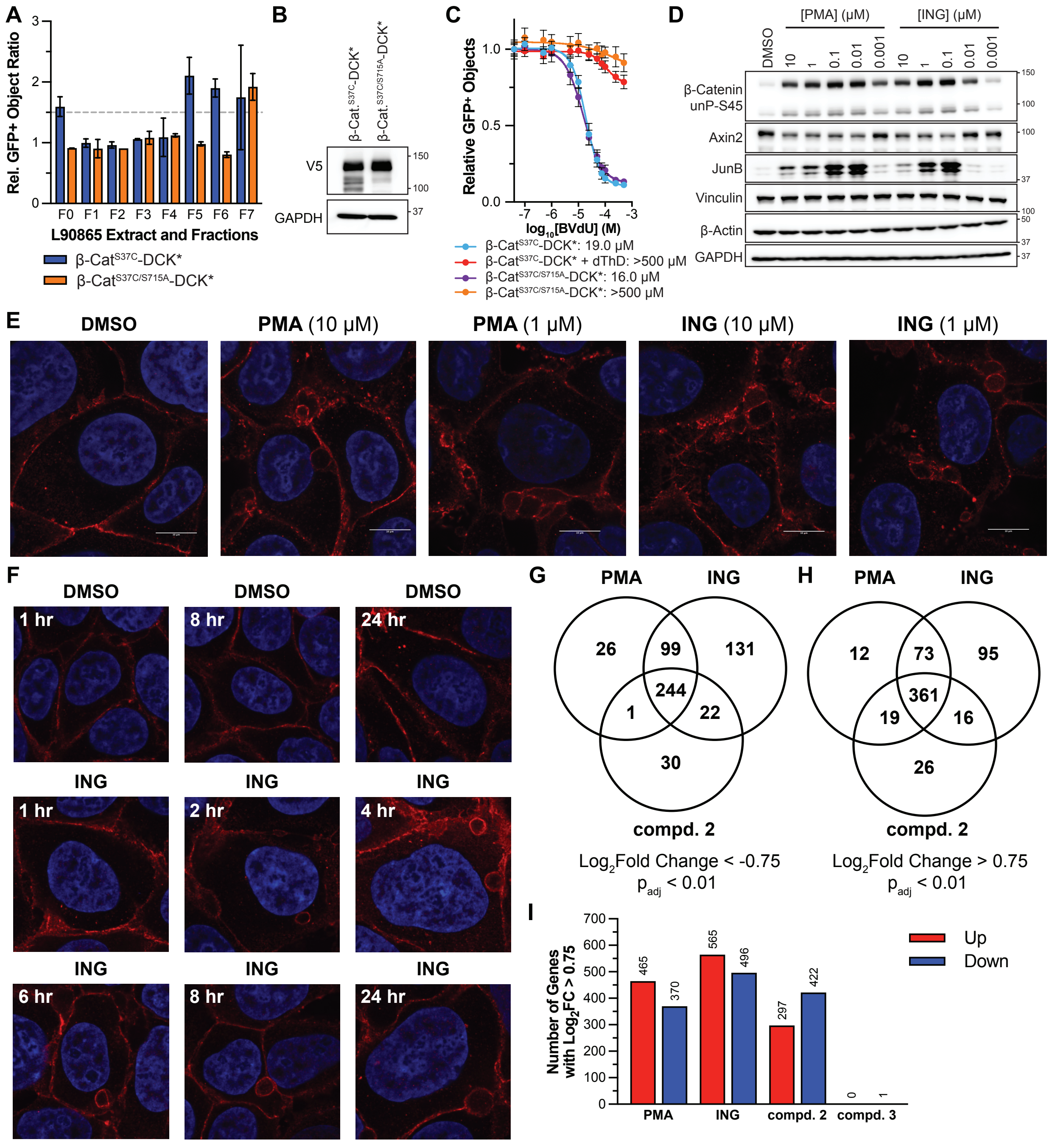
A
Rel. GFP+ Object Ratio
F0 F1 F2 F3 F4 F5 F6 F7
L90865 Extract and Fractions
β-Cat$^{S37C}$-DCK*
β-Cat$^{S37C/S715A}$-DCK*
B
β-Cat.$^{S37C}$-DCK*
β-Cat.$^{S37C/S715A}$-DCK*
V5
GAPDH
150
100
37
C
Relative GFP+ Objects
$\log_{10}$[BVdU] (M)
β-Cat$^{S37C}$-DCK*: 19.0 μM
β-Cat$^{S37C}$-DCK* + dThD: >500 μM
β-Cat$^{S37C/S715A}$-DCK*: 16.0 μM
β-Cat$^{S37C/S715A}$-DCK*: >500 μM
D
[PMA] (μM)
[ING] (μM)
DMSO
10 1 0.1 0.01 0.001
β-Catenin unP-S45
Axin2
JunB
Vinculin
β-Actin
GAPDH
E
DMSO
PMA (10 μM)
PMA (1 μM)
ING (10 μM)
ING (1 μM)
F
DMSO
1 hr
8 hr
24 hr
ING
1 hr
2 hr
4 hr
6 hr
8 hr
24 hr
G
PMA
ING
compd. 2
26
99
131
244
1
22
30
$\mathrm{Log_2}$Fold Change < -0.75
$p_{adj}$ < 0.01
H
PMA
ING
compd. 2
12
73
95
361
19
16
26
$\mathrm{Log_2}$Fold Change > 0.75
$p_{adj}$ < 0.01
I
Number of Genes with $\mathrm{Log_2}$FC > 0.75
465
370
565
496
297
422
0
1
PMA
ING
compd. 2
compd. 3
Up
Down

### SI Fig. 8 (Cont.)

**J** Treatment: PMA

| Rank | MSigDB Hallmark Term | P-value | Combined Score |
|---|---|---|---|
| 1 | Cholesterol Homeostasis | 1.1e-13 | 451.69 |
| 2 | Notch Signaling | 3.8e-05 | 116.96 |
| 3 | Estrogen Response Early | 1.7e-08 | 98.05 |
| 4 | Estrogen Response Late | 7.3e-08 | 86.26 |
| 5 | Wnt-beta Catenin Signaling | 1.6e-04 | 74.48 |

**K** Treatment: ING

| Rank | MSigDB Hallmark Term | P-value | Combined Score |
|---|---|---|---|
| 1 | Cholesterol Homeostasis | 7.6e-14 | 391.14 |
| 2 | Estrogen Response Late | 7.6e-11 | 132.26 |
| 3 | Estrogen Response Early | 1.1e-07 | 127.50 |
| 4 | Notch Signaling | 1.9e-04 | 72.75 |
| 5 | Wnt-beta Catenin Signaling | 7.6e-04 | 45.23 |

**L** Treatment: Compound 2

| Rank | MSigDB Hallmark Term | P-value | Combined Score |
|---|---|---|---|
| 1 | Cholesterol Homeostasis | 3.5e-06 | 106.96 |
| 2 | Wnt-beta Catenin Signaling | 4.8e-05 | 106.56 |
| 3 | Notch Signaling | 1.6e-04 | 100.88 |
| 4 | Androgen Response | 4.3e-05 | 60.86 |
| 5 | Estrogen Response Early | 1.9e-05 | 48.72 |

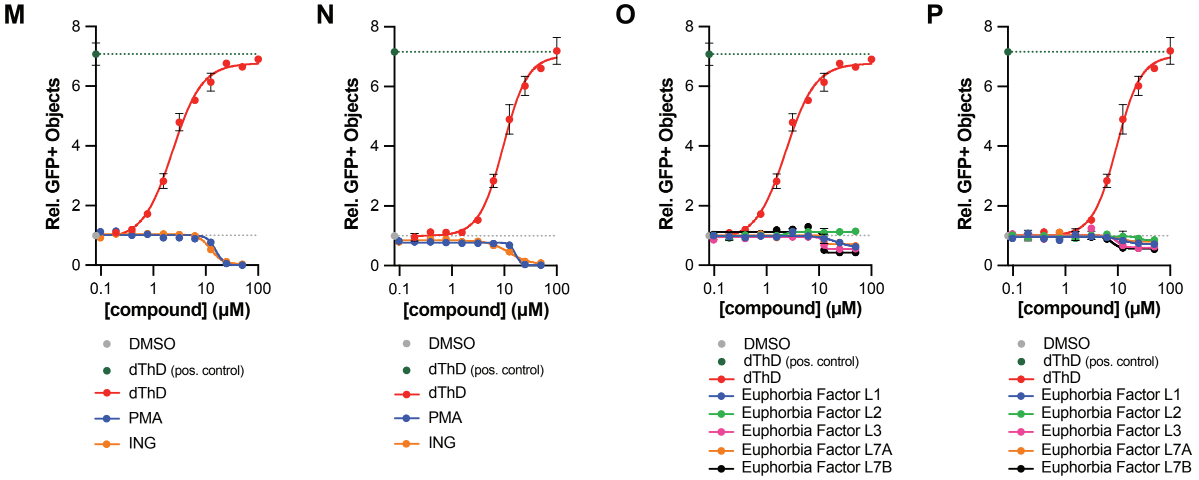


**SI Figure 8. PKC Activators Phenocopy β-cat Inactivators.**
(**A**) Relative GFP+ objects for 293FT TK1 KO β-cat$^{S37C}$-DCK* and 293FT TK1 KO β-cat$^{S37C/S715A}$-DCK* cells treated with L90865 extract and fractions for 24 hours followed by BVdU treatment (50 μM) for 96 hours. n = 2 technical replicates. (**B**) Immunoblot analysis of 293FT TK1 KO β-cat$^{S37C}$-DCK* and 293 FT TK1 KO β-cat$^{S37C/S715A}$-DCK* cells used for BVdU assays. (**C**) Relative GFP+ objects for 293FT cells as in (**A**) treated with DMSO or 100 μM dThD for 24 hours followed by indicated concentrations of BVdU for 96 hours. n = 10 technical replicates. (**D**) Immunoblot analysis of 293FT TK1 KO β-cat$^{S37C}$-DCK* cells treated with the indicated concentrations of PMA or ING for 24 hours. n = 3 biological replicates. (**E**) Immunofluorescence of DLD-1 cells treated with the indicated concentrations of PMA or ING for 24 hours. Blue: DAPI stain, Red: β-Cat. Scale bar = 10 μm. (**F**) Immunofluorescence of DLD-1 cells treated with the indicated compounds for the indicated times. Blue: DAPI stain, Red: β-Cat. (**G-L**) RNAseq analysis of DLD-1 cells treated with the indicated compounds for 24 hours. Cells were treated with either PMA (1 μM), ING (1 μM), compound **2** (25 μM), or compound **3** (25 μM). (**G**,**H**) Venn diagram of genes that decreased (**G**) or increased (**H**) in expression as compared to DMSO control. (**I**) Plot of the number of genes up- or downregulated ($|log_2FC| < 0.75$, $p_{adj} < 0.01$). Data is from 2 technical replicates. (**J-L**) Gene set enrichment analysis (MSigDB Hallmark 2020) of downregulated genes ($log_2FC < -0.75$, $p_{adj} < 0.01$) for cells treated with the indicated compounds. (**M-P**) Relative GFP+ objects of 293FT TK1 KO DCK* cells (**M**,**O**) or of 293FT TK1 KO β-cat$^{S37C}$-DCK* cells (**N**,**P**) treated with the indicated compounds for 24 hours followed by BVdU (10 μM for DCK* cells and 50 μM for β-Cat$^{S37C}$-DCK* cells) for 96 hours. n = 3 for technical replicates.

A

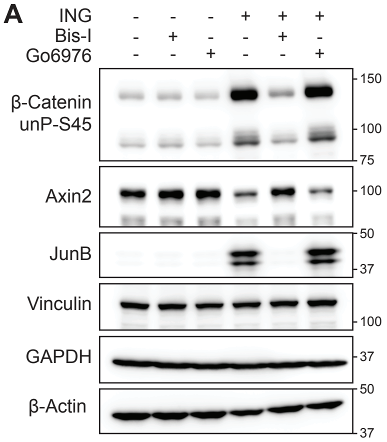


B

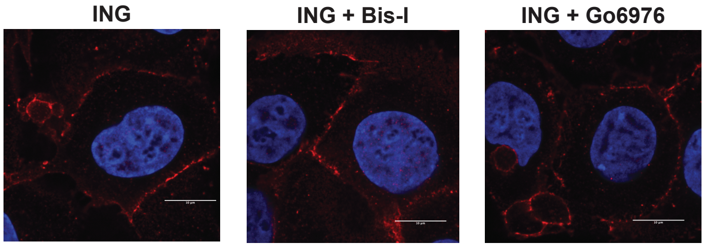


C

**PKCδ**

| - | Name | Amino Acid Change |
|---|---|---|
| WT | Wild-Type | n/a |
| KD | Kinase-Dead | K378R |
| CA | Constitutively Active | R144A, R145A |

**PKCθ**

| - | Name | Amino Acid Change |
|---|---|---|
| WT | Wild-Type | n/a |
| KD | Kinase-Dead | K409R |
| CA | Constitutively Active | R145A, R146A |

D

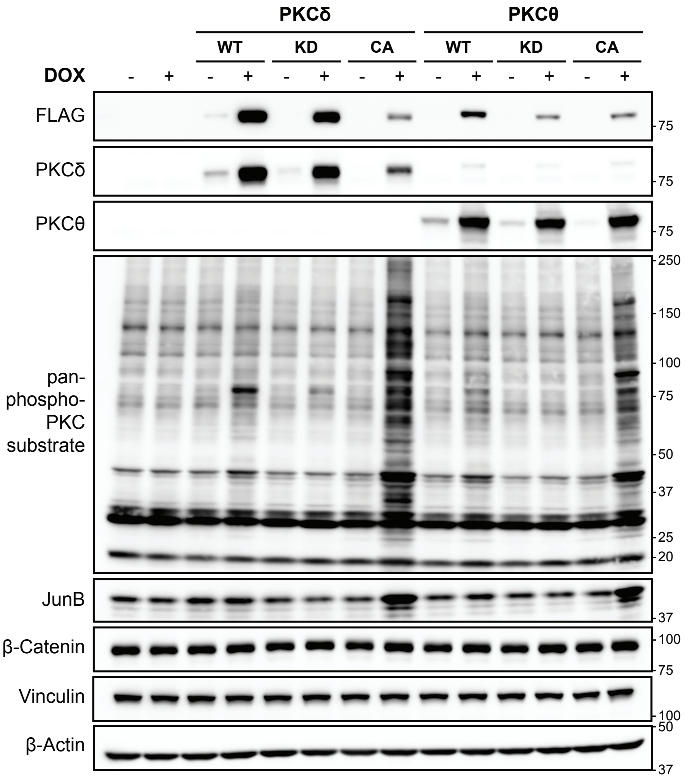

### SI Figure 9 (cont.)

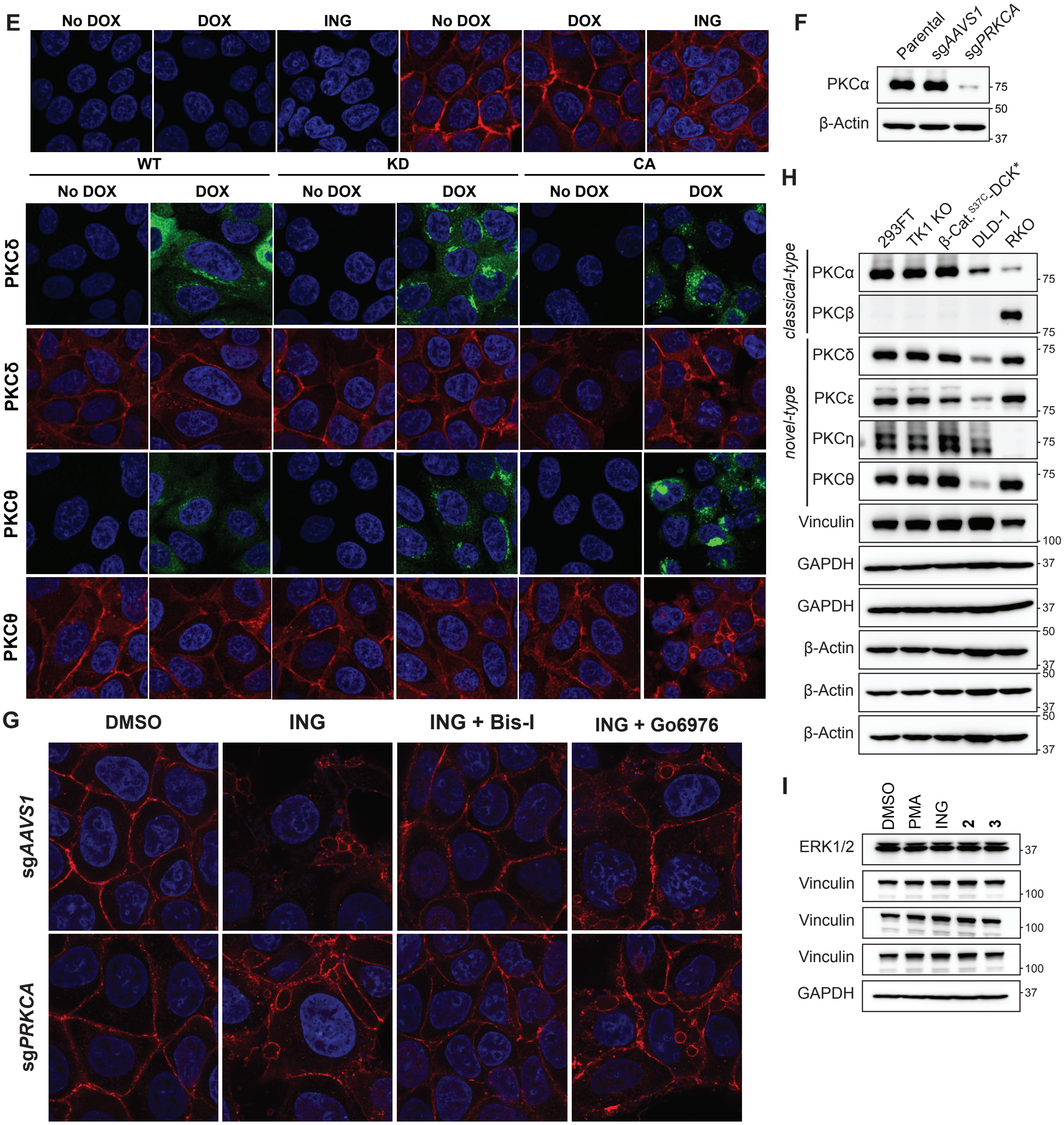


**SI Figure 9. Expression of Active Novel PKCs Phenocopy β-cat Inactivators.**
(**A**) Immunoblot analysis of 293FT TK1 KO β-cat$^{S37C}$-DCK* cells treated, where indicated, with ING (1 μM), Bis-I (1 μM), or Go6976 (0.5 μM) for 24 hours. n = 3 biological replicates. (**B**) Immunofluorescence imaging of DLD-1 cells treated, where indicated, with ING (1 μM), Bis-I (1 μM), or Go6976 (0.5 μM) for 24 hours. These cells were a part of the same experiment that generated the images shown in **Fig. 4E**. (**C**) Summary of amino acid changes for wild-type (WT), kinase-dead (KD), or constitutively active (CA) PKCδ and PKCθ variants. (**D**) Immunoblot analysis of DLD-1 cells expressing the indicated doxycycline-inducible PKCδ or PKCθ variants. The exogenous PKCs have a FLAG tag. Images are representative of 3 biological replicates. (**E**) Immunofluorescence imaging of DLD-1 cells (as in **D**). Images are representative of 3 biological replicates. Blue: DAPI stain, Green: FLAG, Red: β-Cat. (**F**) Immunoblot analysis of DLD-1 cells stably expressing Cas9 and the indicated CRISPR sgRNAs. (**G**) Immunofluorescence imaging of DLD-1 sg*AAVS1*

or sg*PRKCA* cells treated, where indicated, with ING (1 μM), Bis-I (1 μM), or Go6976 (1 μM) for 24 hours. (**H**) Immunoblot analysis of 293FT, 293FT TK1 KO, 293FT TK1 KO β-cat$^{S37C}$-DCK*, DLD-1, and RKO colorectal cancer cells. (**I**) Further immunoblot loading controls for experiment shown in **Fig. 4F**.

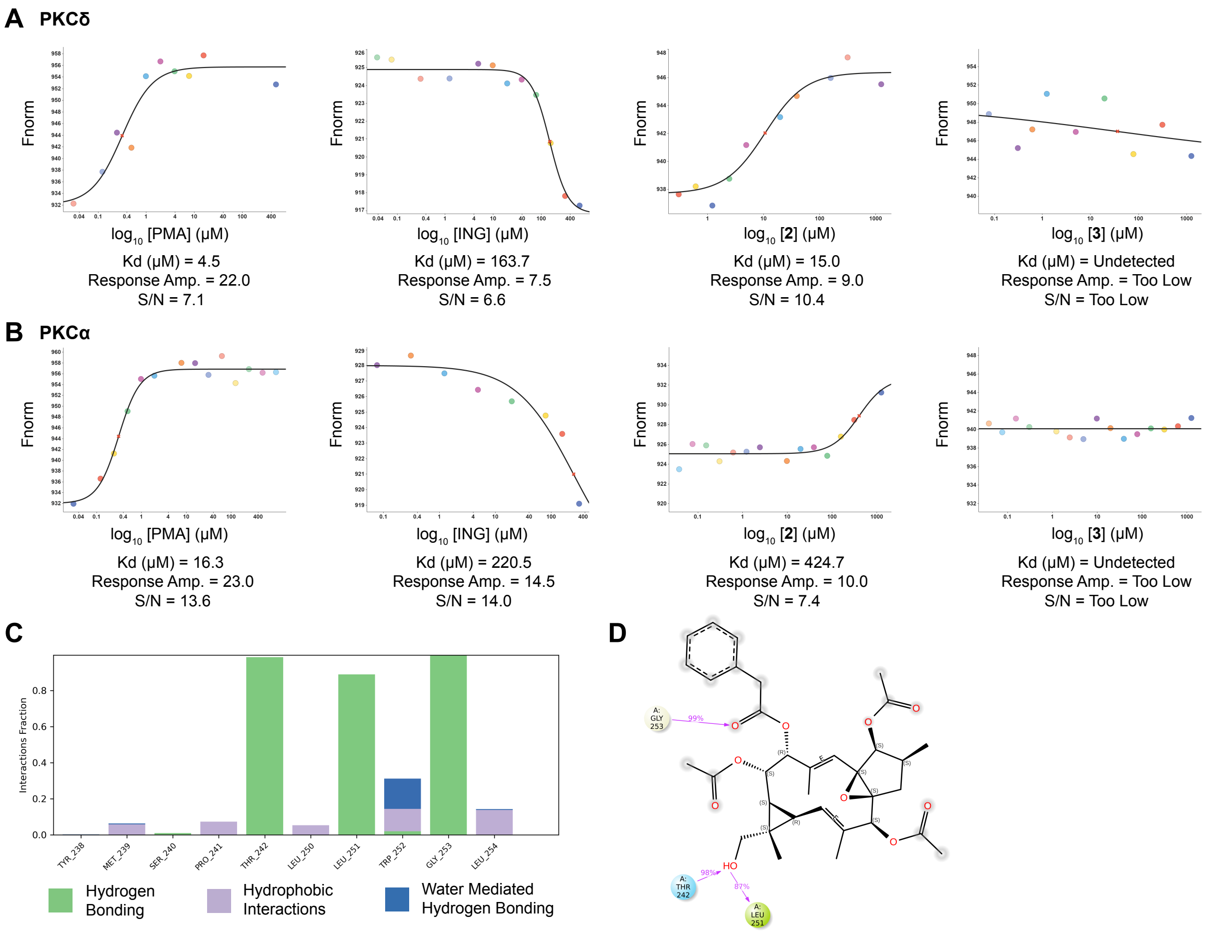

A PKCδ
Fnorm
log10 [PMA] (μM)
Kd (μM) = 4.5
Response Amp. = 22.0
S/N = 7.1
log10 [ING] (μM)
Kd (μM) = 163.7
Response Amp. = 7.5
S/N = 6.6
log10 [2] (μM)
Kd (μM) = 15.0
Response Amp. = 9.0
S/N = 10.4
log10 [3] (μM)
Kd (μM) = Undetected
Response Amp. = Too Low
S/N = Too Low
B PKCα
log10 [PMA] (μM)
Kd (μM) = 16.3
Response Amp. = 23.0
S/N = 13.6
log10 [ING] (μM)
Kd (μM) = 220.5
Response Amp. = 14.5
S/N = 14.0
log10 [2] (μM)
Kd (μM) = 424.7
Response Amp. = 10.0
S/N = 7.4
log10 [3] (μM)
Kd (μM) = Undetected
Response Amp. = Too Low
S/N = Too Low
C
Interactions Fraction
TYR_238
MET_239
SER_240
PRO_241
THR_242
LEU_250
LEU_251
TRP_252
GLY_253
LEU_254
Hydrogen Bonding
Hydrophobic Interactions
Water Mediated Hydrogen Bonding
D

### SI Figure 10 (cont.)

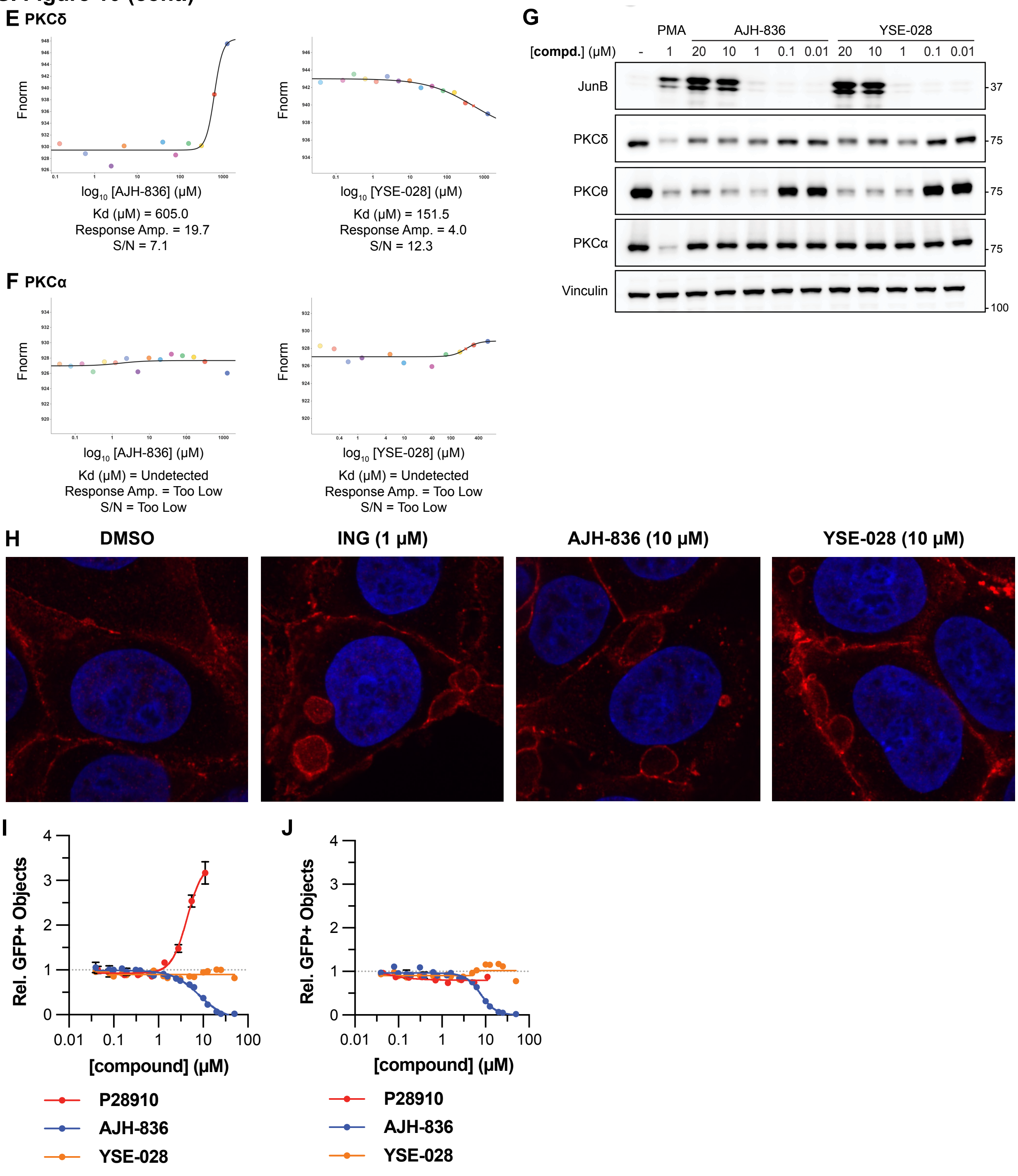


**SI Figure 10. Compound 2 Binds PKCδ With Higher Affinity Than PKCα.**
(**A**,**B**) MST curves and values ($K_d$, Response Amplitude (Amp.), Signal-to-Noise (S/N)) for the indicated compounds (PMA, ING, **2**, or **3**) for binding to PKCδ (**A**) and PKCα (**B**). (**C**) Plot of interactions fraction for amino acid residues within the C1b domain of PKCδ (PDB: 7LF3) that have contacts with compound **2** during in silico docking and molecular dynamic simulations. (**D**) Two-dimensional representation of the molecular modeling of compound **2** bound to the C1b domain of PKCδ (PDB: 7LF3). Key hydrogen bonds are shown with purple arrows. (**E**,**F**) MST curves and values ($K_d$, Response Amplitude (Amp.), Signal-to-Noise (S/N)) for the indicated compounds (AJH-836 or YSE-028) for binding of PKCδ (**E**) and PKCα (**F**).

(**G**) Immunoblot analysis of 293FT cells treated with the indicated compounds and doses for 24 hours. n = 3 biological replicates. (**H**) Immunofluorescence imaging of DLD-1 cells treated with the indicated compounds and doses. Images are representative of 3 biological replicates. Blue: DAPI stain, Red: β-Cat. (**I**,**J**) Relative GFP+ objects of 293FT TK1 KO β-cat$^{S37C}$-DCK* cells (**I**) or of 293FT TK1 KO β-cat$^{S37C/S715A}$-DCK* cells (**J**) treated with the indicated compounds for 24 hours followed by BVdU (50 μM) for 96 hours. n = 3 for technical replicates. **P28910** is an impure natural product sample derived from the L34409 extract lineage (a β-cat inactivator). Since **P28910** is impure and the active principle's molecular weight is unknown, the concentration is nominal and an estimation of its concentration was used (based on an assumption of a molecular weight of 600 g/mol).

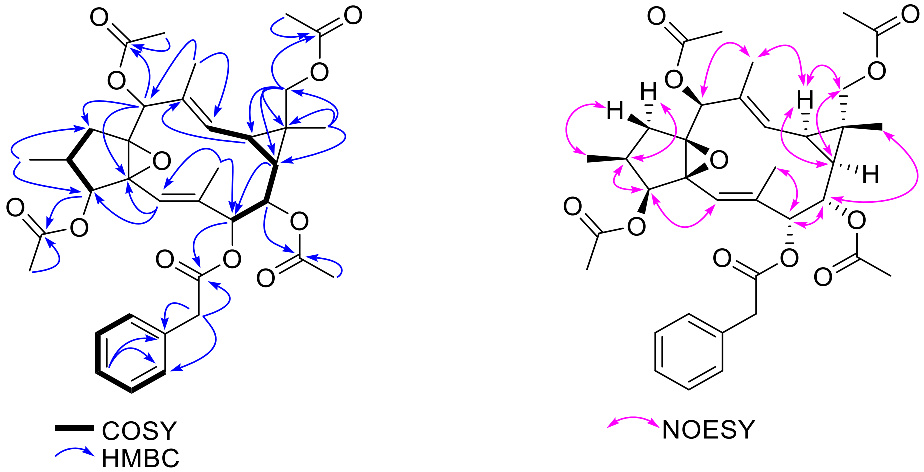


**SI Figure 11.** Key $^{1}H$–$^{1}H$ COSY, $^{1}H$–$^{13}C$ HMBC and $^{1}H$–$^{1}H$ NOESY correlations for compound **3**.

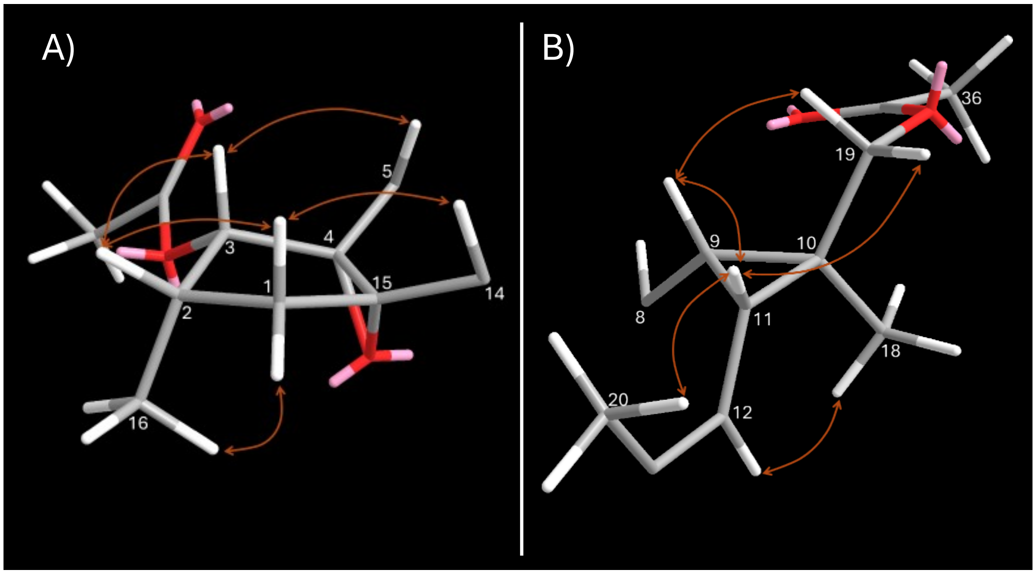


**SI Figure 12.** Key NOE correlations observed in (a) five membered-ring and (b) cyclopropane-ring of compound **3**.

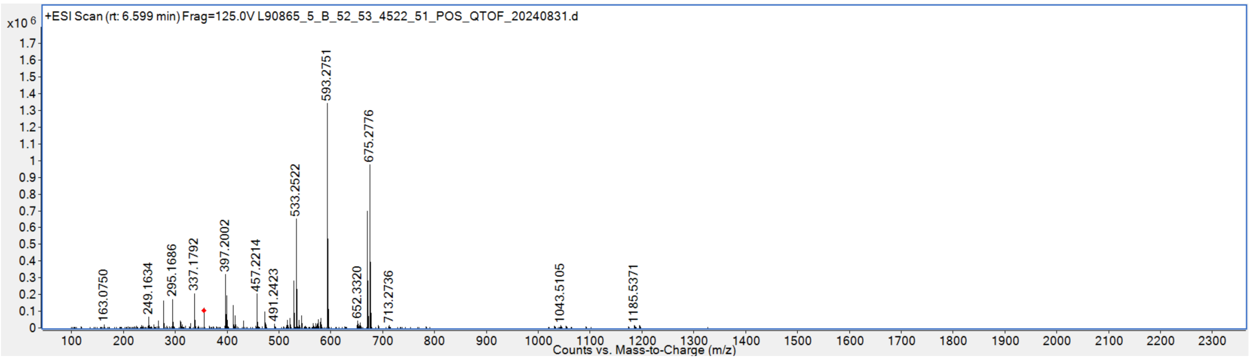


**SI Figure 13.** HRESIMS for compound **3**.

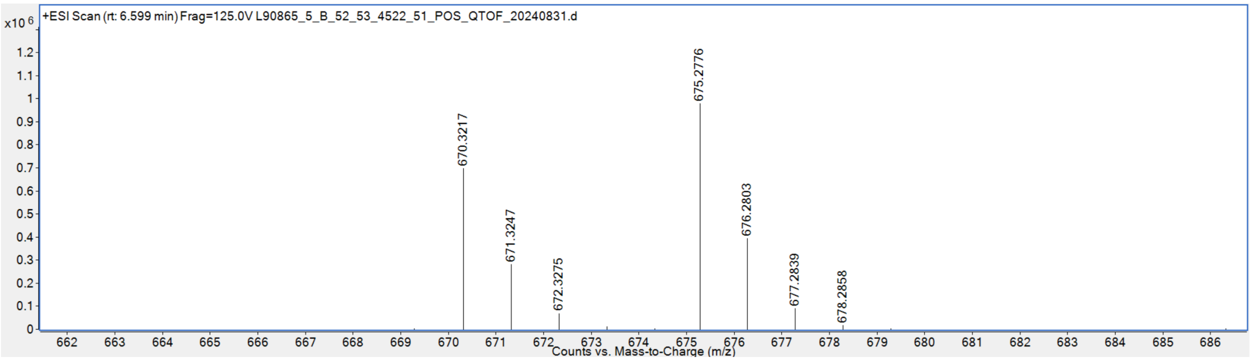


**SI Figure 14.** Expansion of HRESIMS for compound **3**.

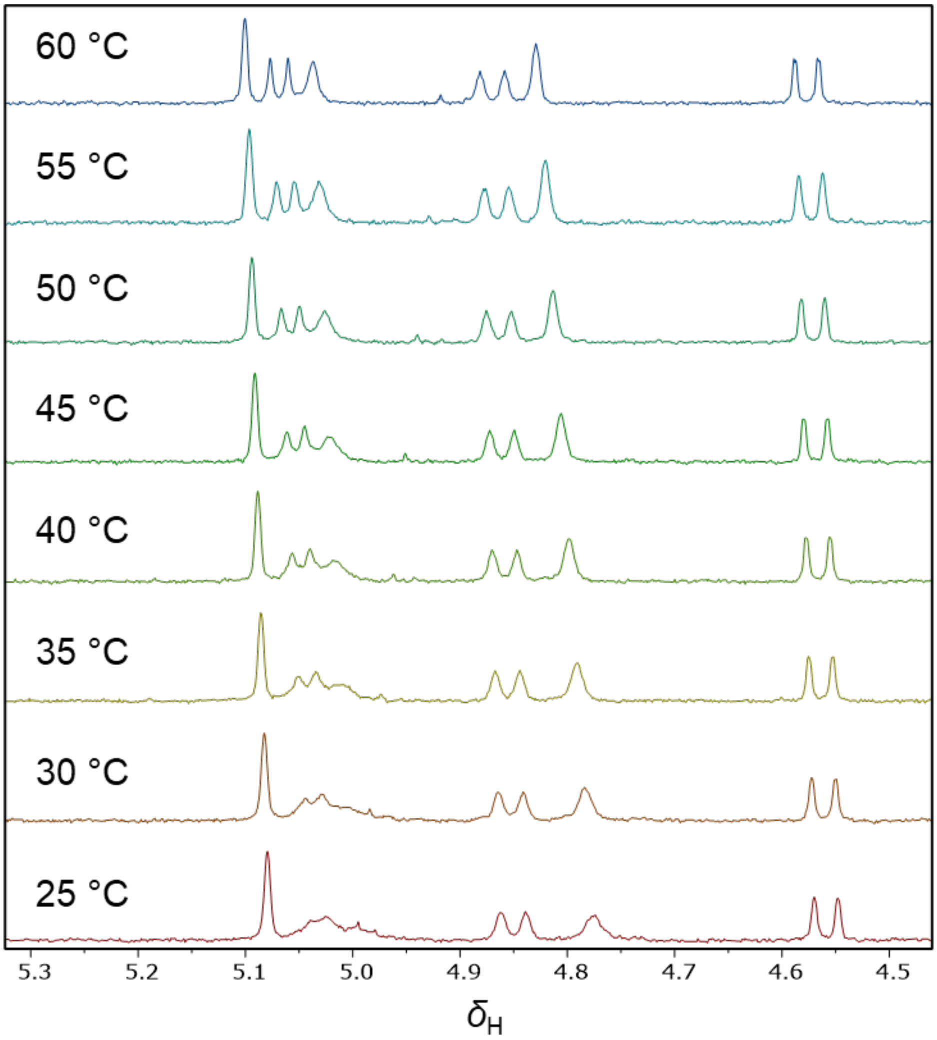


**SI Figure 15.** 500 MHz [1]H NMR spectra of compound **3** in DMSO-$d_6$ at variable temperature (25 – 60 °C).

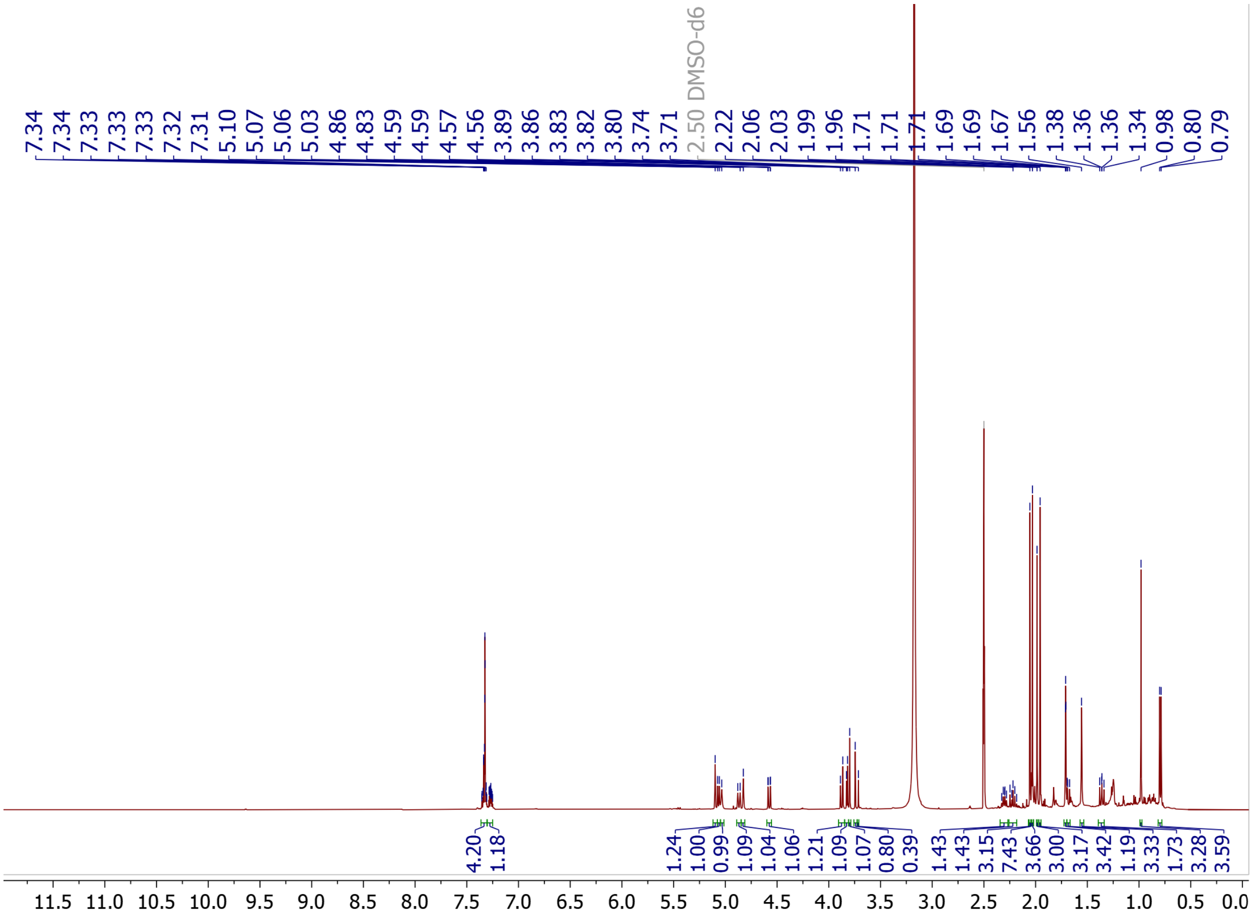

**SI Figure 16.** 500 MHz $^1$H NMR spectrum of compound **3** in DMSO-$d_6$. The spectrum was acquired over 64 scans at 60°C.

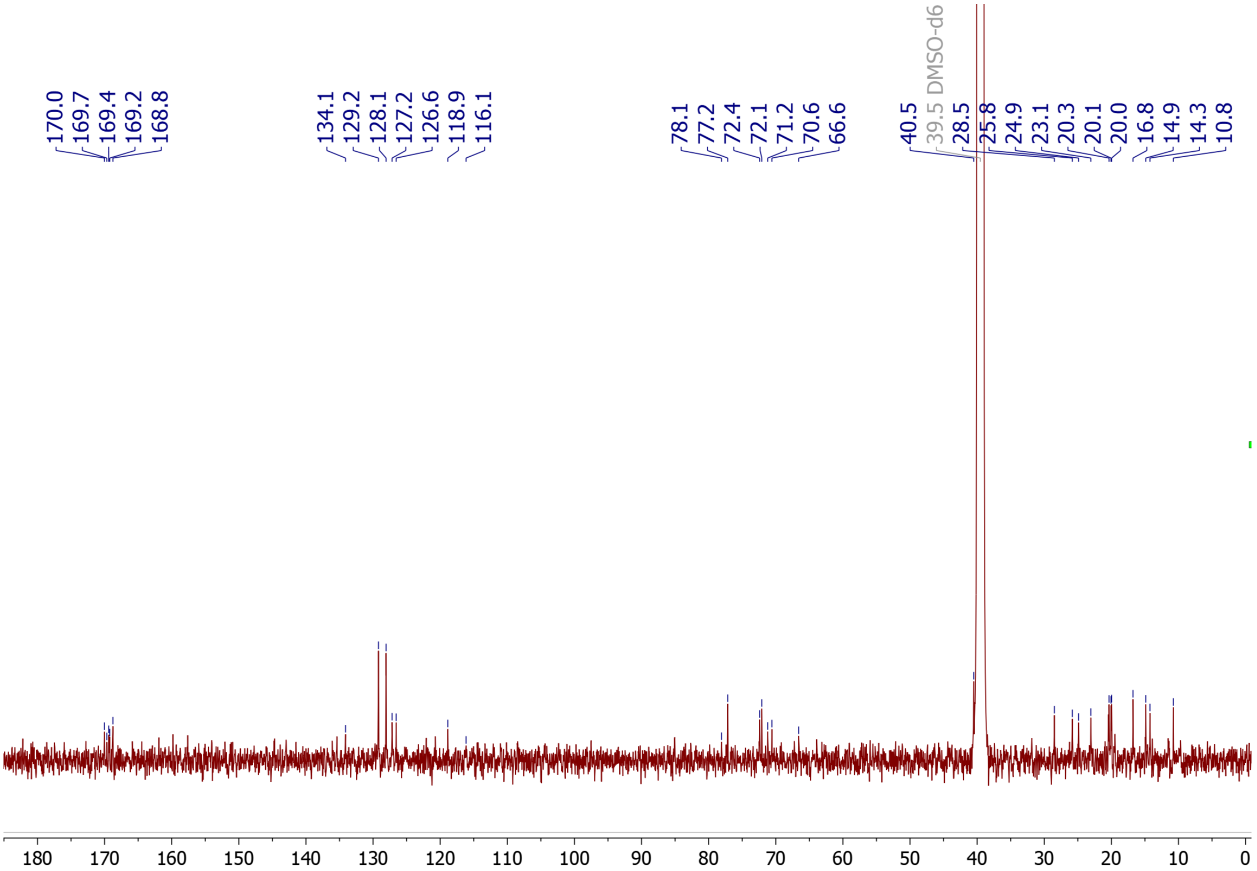


**SI Figure 17.** 125.8 MHz $^{13}C$ NMR spectrum of compound **3** in DMSO-$d_6$. The spectrum was acquired over 3017 scans at 60°C.

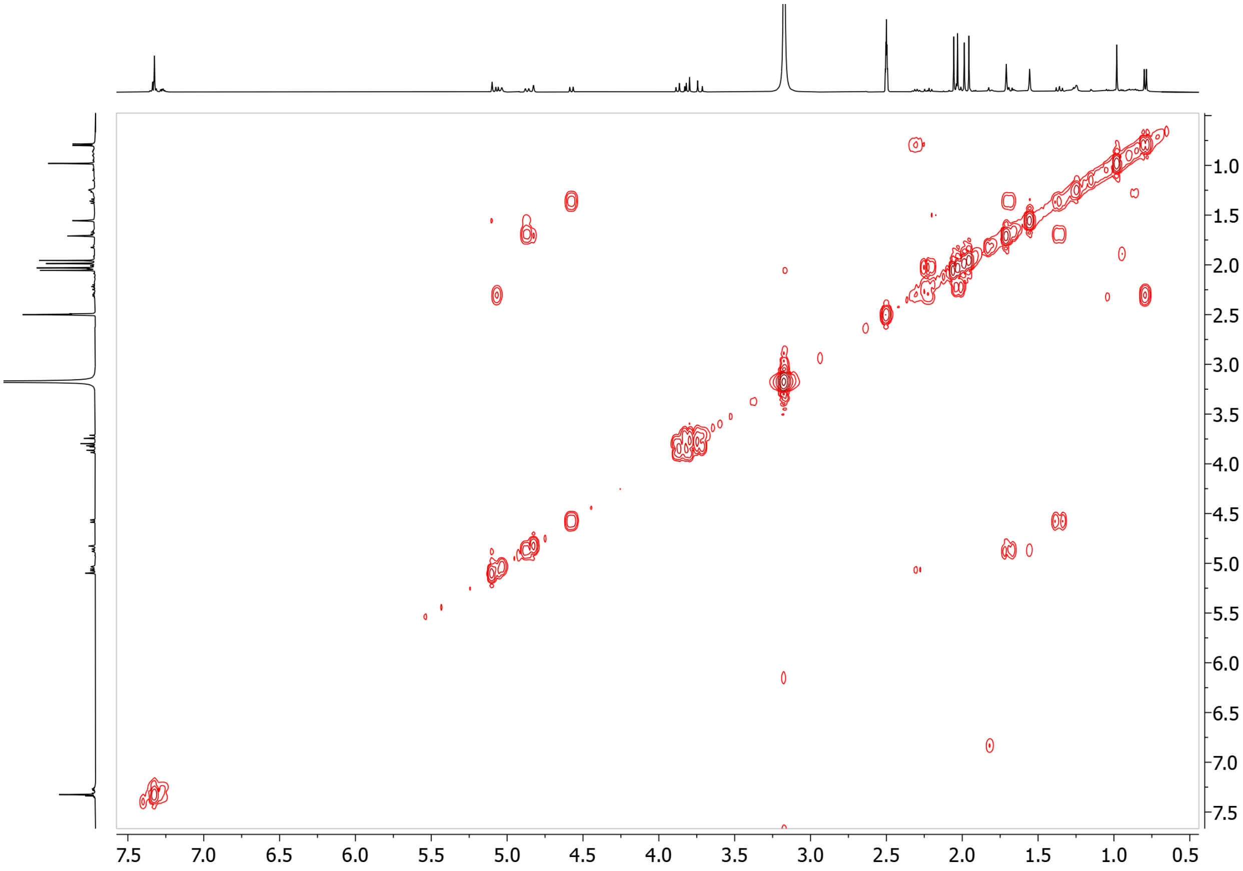


**SI Figure 18.** $^{1}H$–$^{1}H$ COSY NMR spectrum of compound **3** in DMSO-$d_6$. The spectrum was acquired over 8 scans at 60 °C.

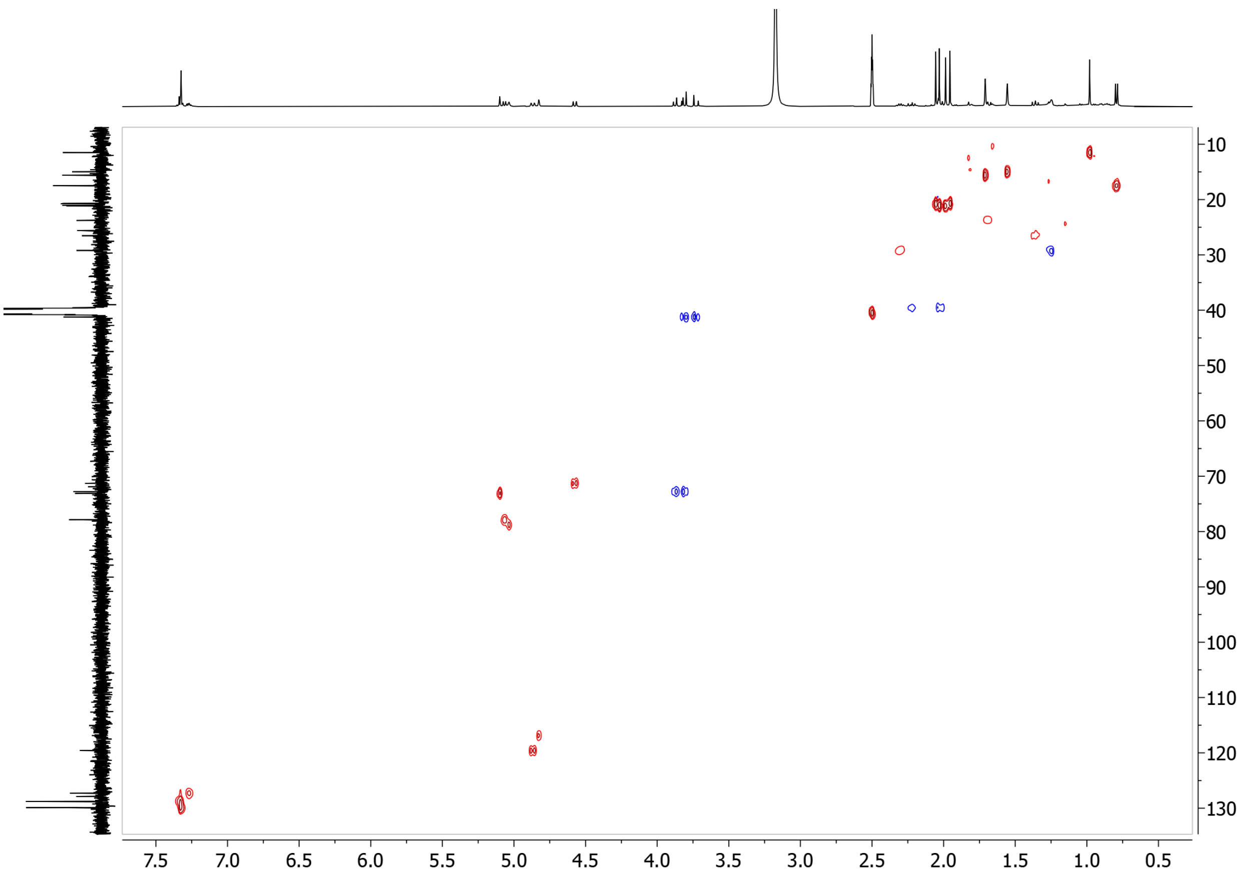


**SI Figure 19.** $^1H$–$^{13}C$ HSQC NMR spectrum of compound **3** in DMSO-$d_6$. The spectrum was acquired over 32 scans at 60 °C.

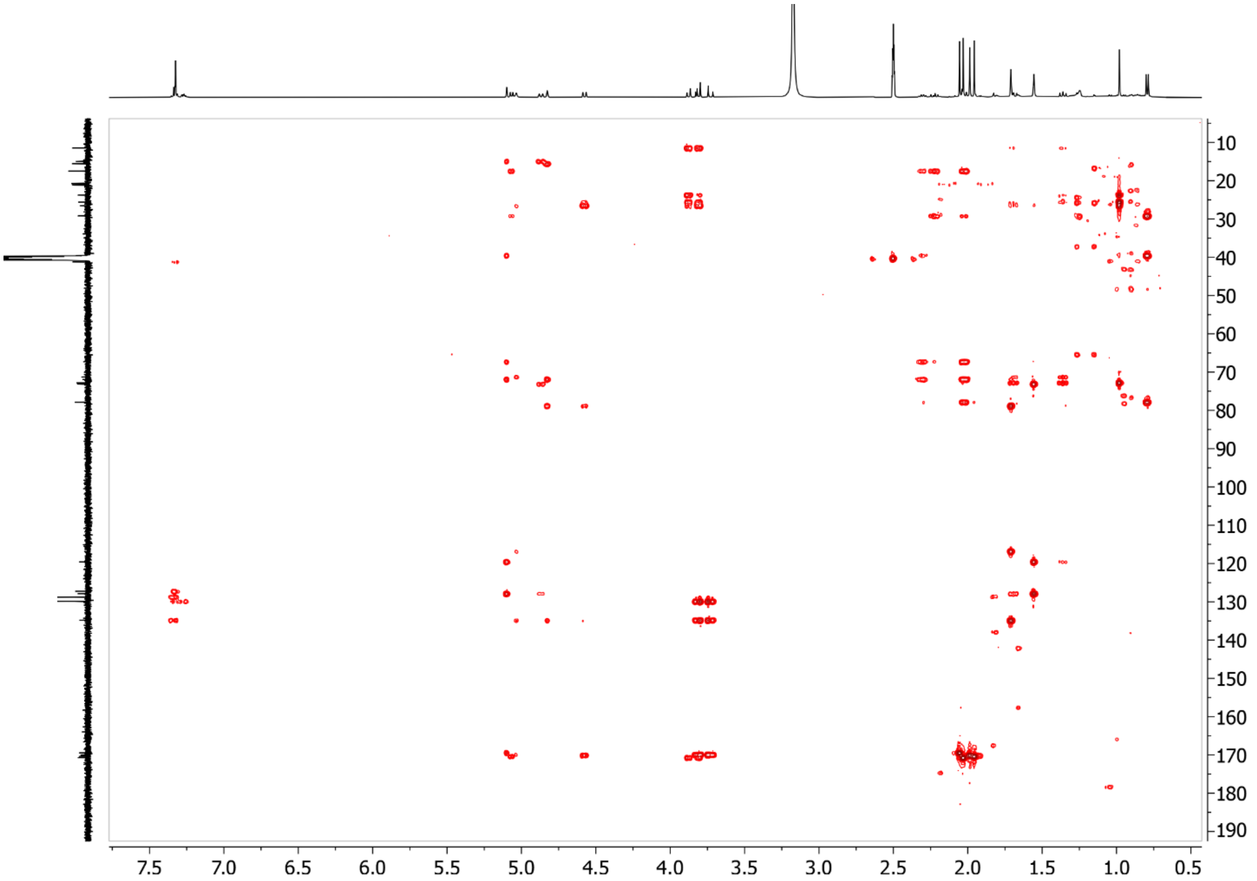


**SI Figure 20.** [1]H–[13]C HMBC NMR spectrum of compound **3** in DMSO-$d_6$. The spectrum was acquired over 32 scans at 60 °C.

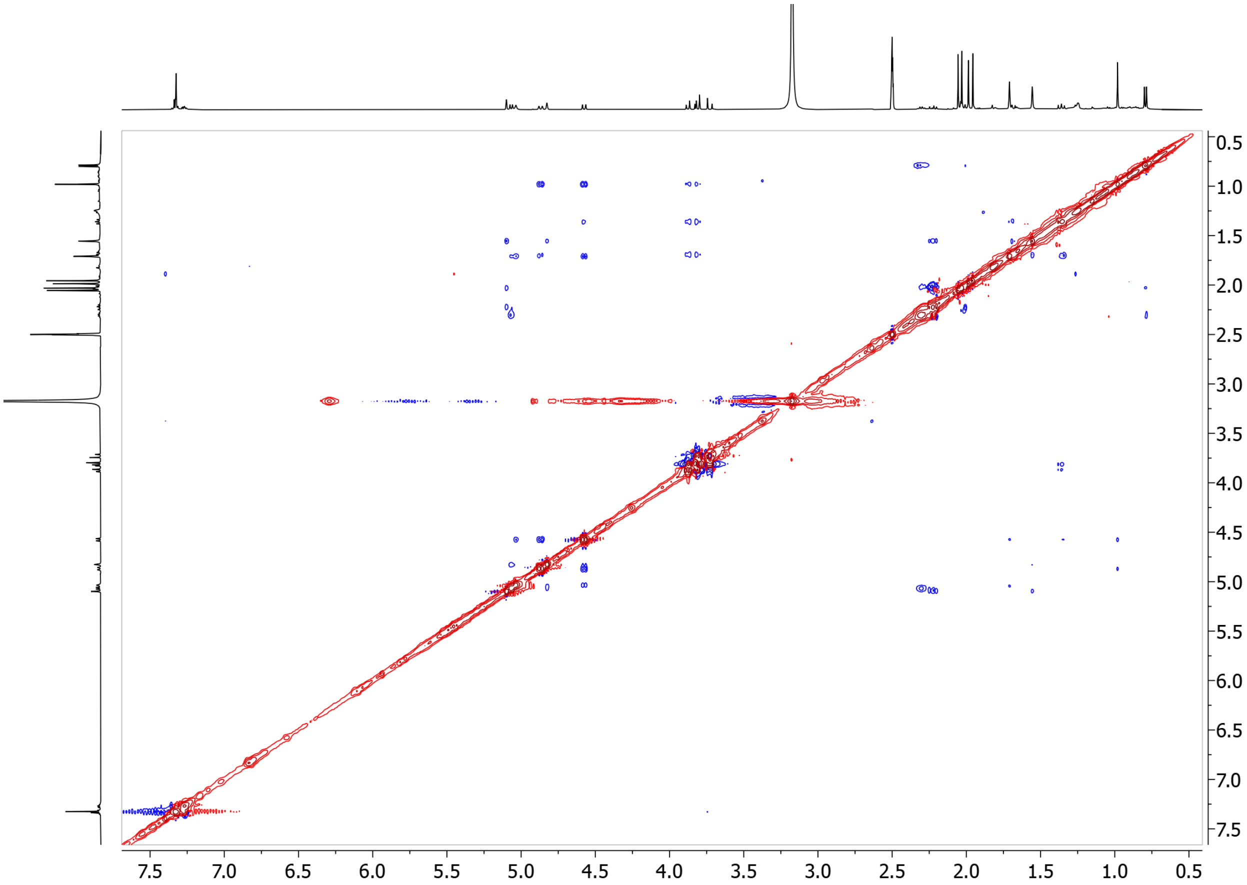


**SI Figure 21.** 2D NOESY NMR spectrum of compound **3** in DMSO-$d_6$. The spectrum was acquired over 16 scans at 60 °C.

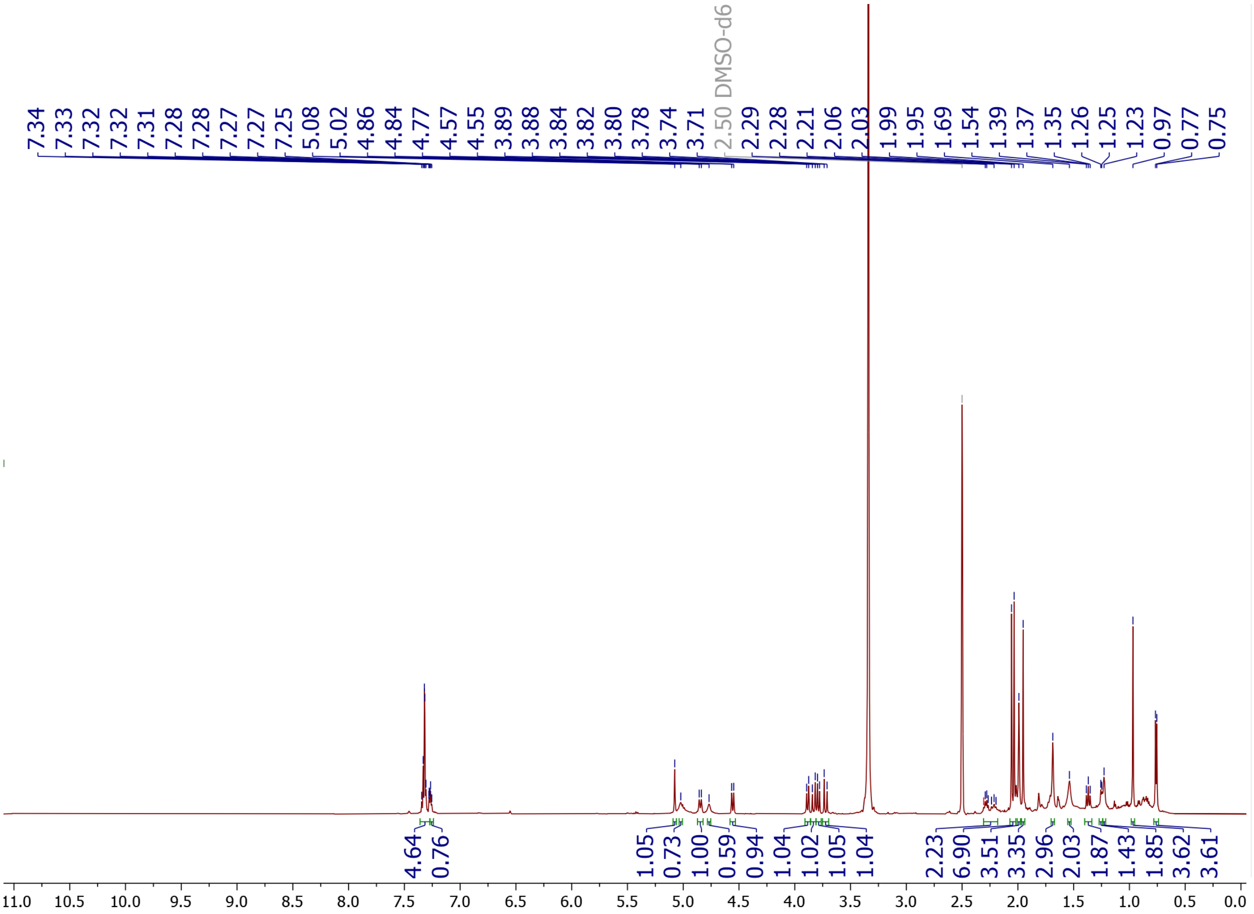

**SI Figure 22.** 600 MHz $^1$H NMR spectrum of compound **3** in DMSO-$d_6$. The spectrum was acquired over 4 scans at 25 °C.

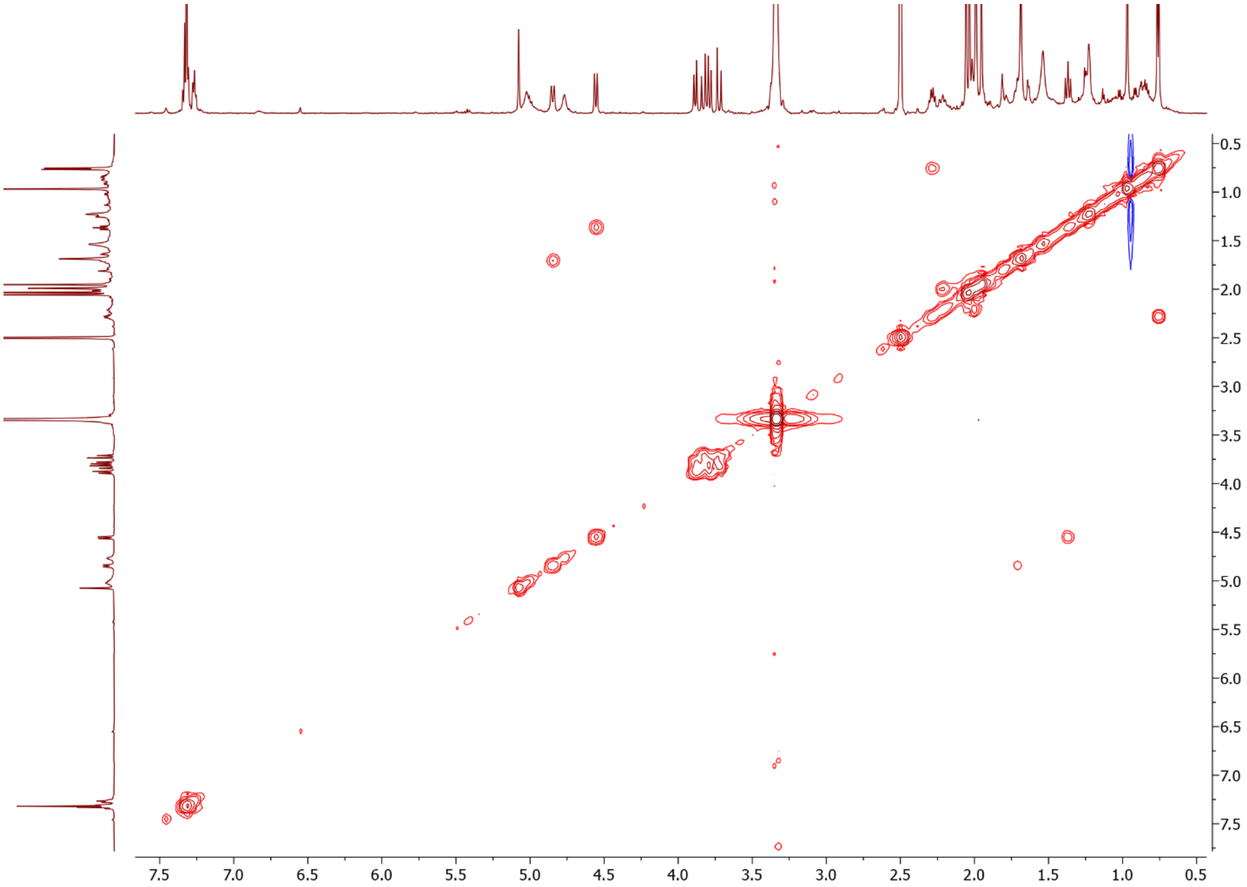


**SI Figure 23.** $^{1}H$–$^{1}H$ COSY NMR spectrum of compound **3** in DMSO-$d_6$. The spectrum was acquired over 96 scans at 25 °C.

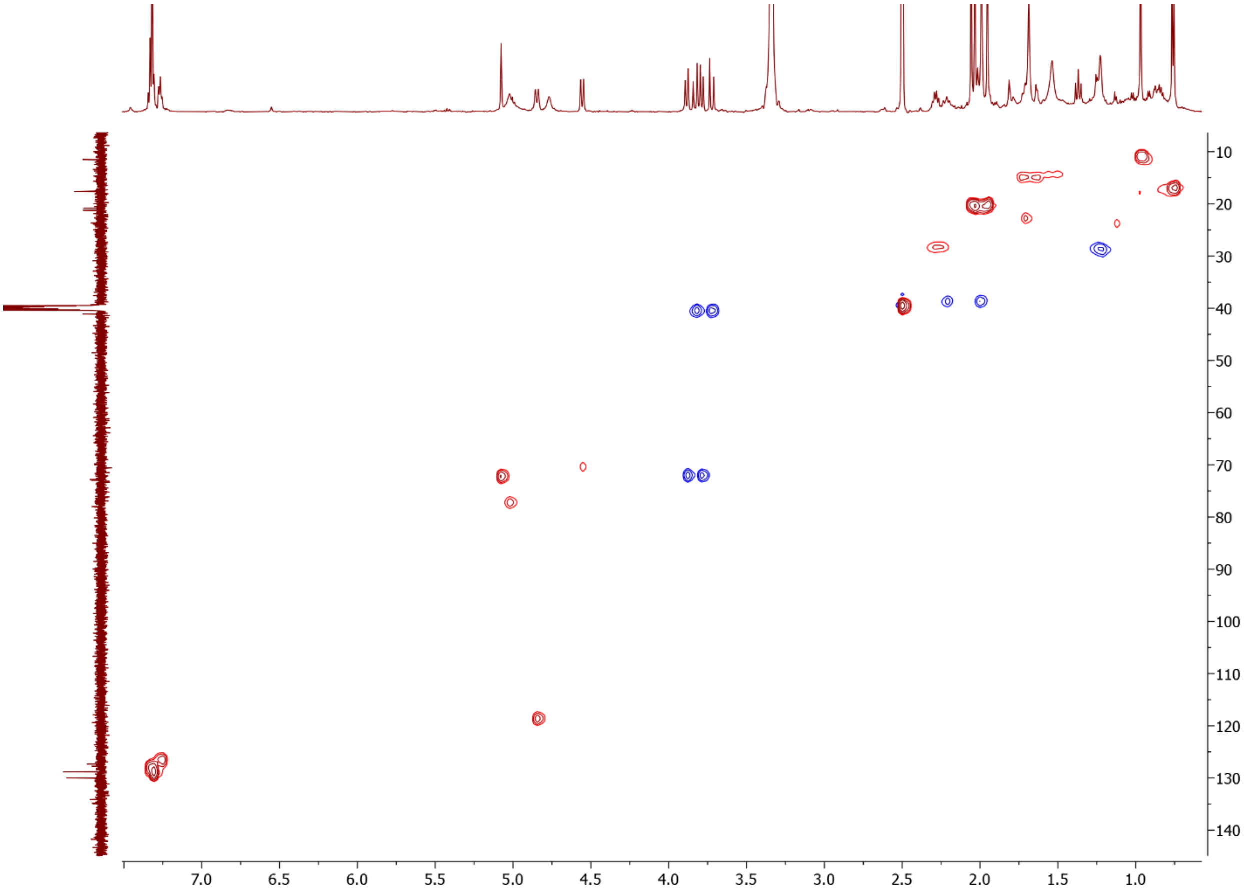


**SI Figure 24.** $^{1}H$–$^{13}C$ HSQC NMR spectrum of compound **3** in DMSO-$d_6$. The spectrum was acquired over 96 scans at 25 °C.

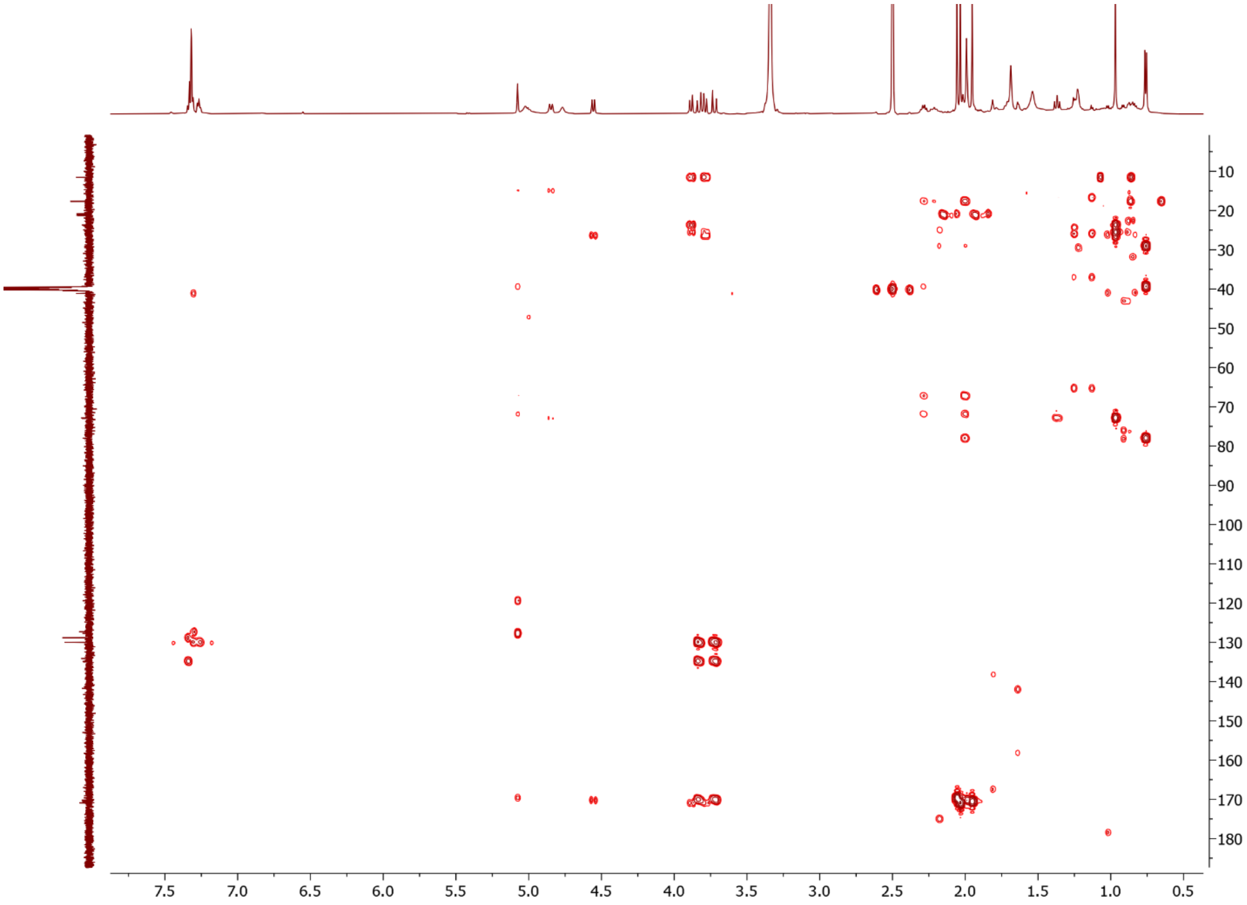


**SI Figure 25.** $^{1}H$–$^{13}C$ HMBC NMR spectrum of compound **3** in DMSO-$d_6$. The spectrum was acquired over 960 scans at 25 °C.

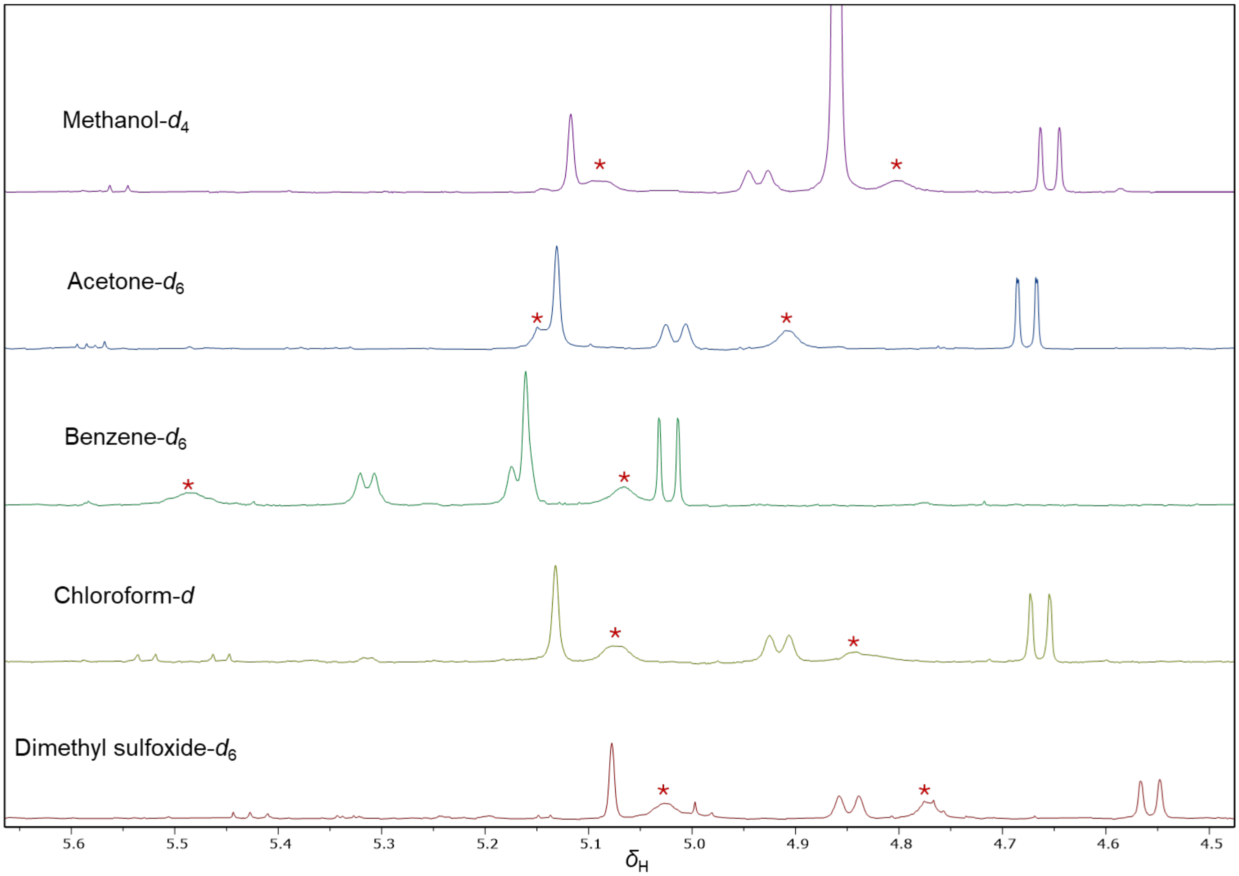


**SI Figure 26.** $^1$H NMR spectra comparison of compound **3** in different NMR solvents.

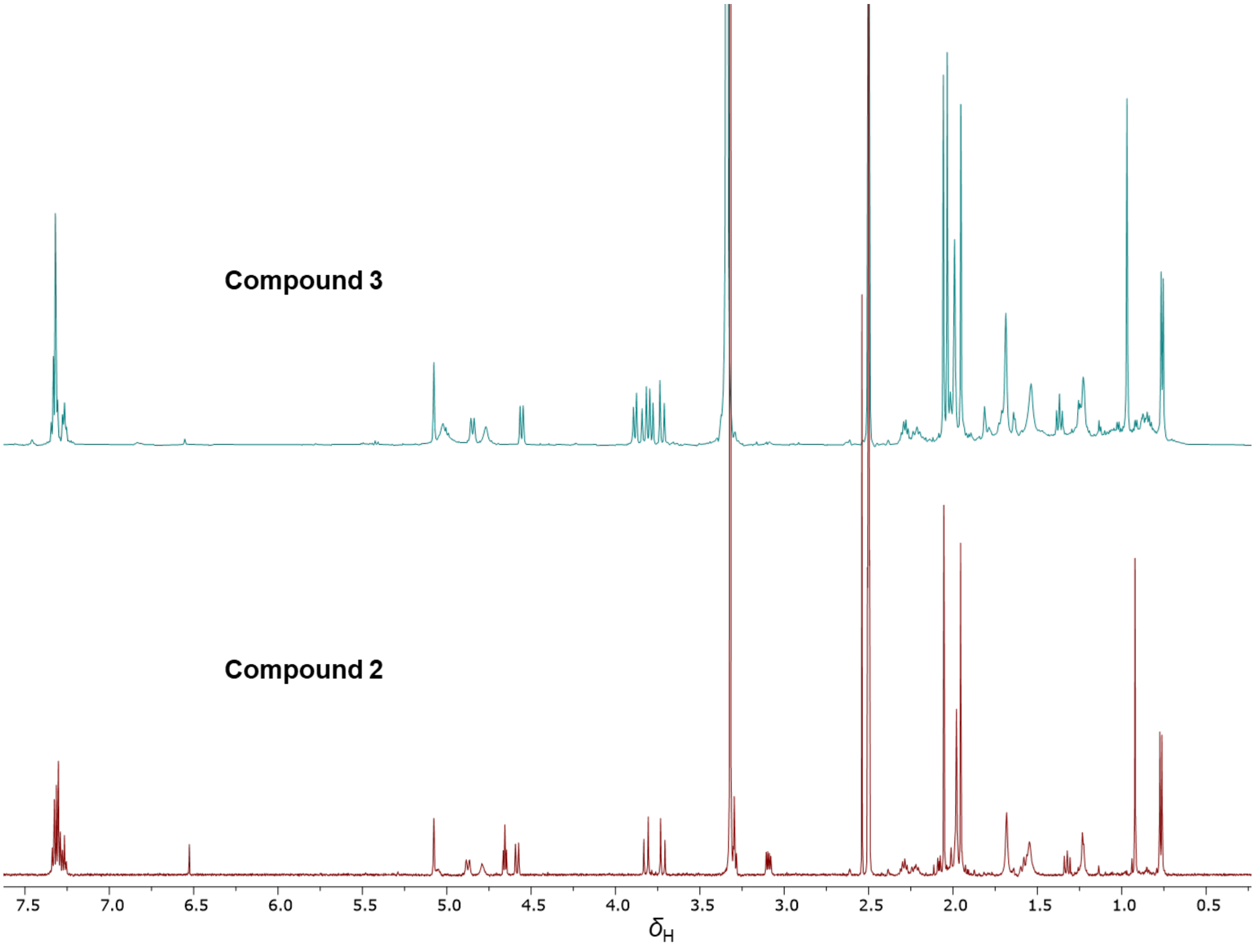


**SI Figure 27.** Comparison of [1]H NMR spectrum of compound **3** and compound **2** in DMSO-$d_6$ at 25°C.

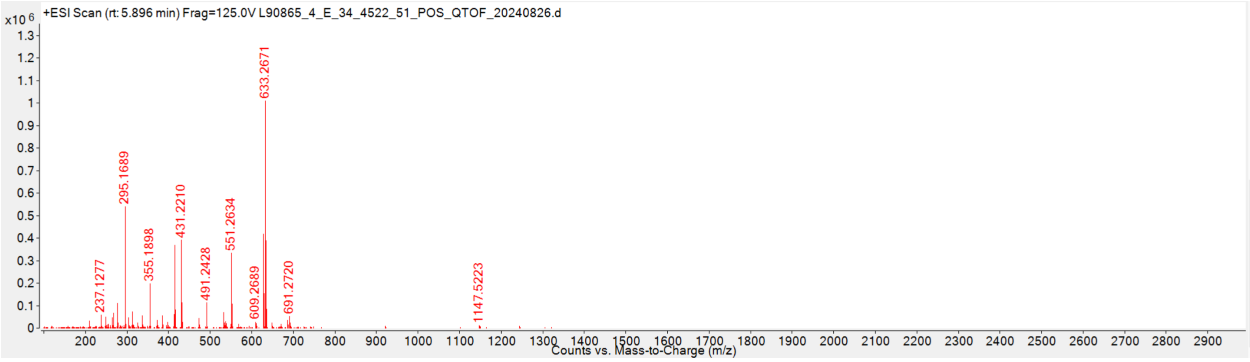


**SI Figure 28.** HRESIMS for compound **2**.

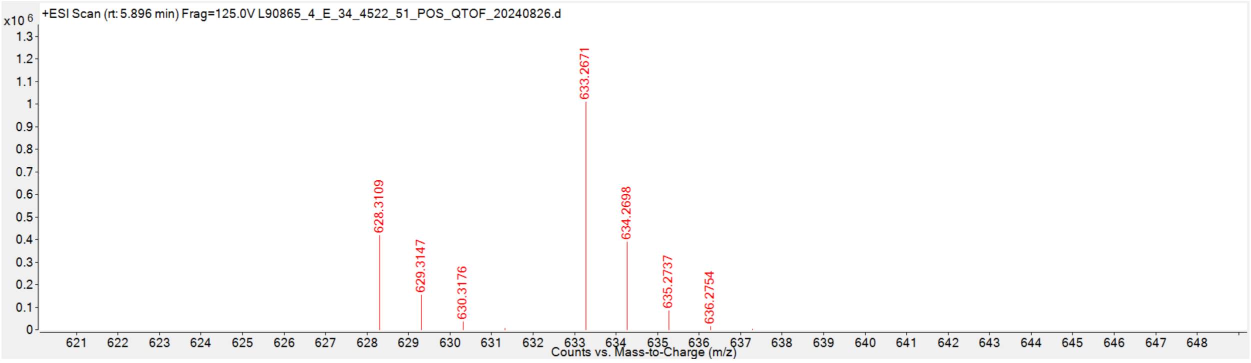


**SI Figure 29.** Expansion of HRESIMS for compound **2**.

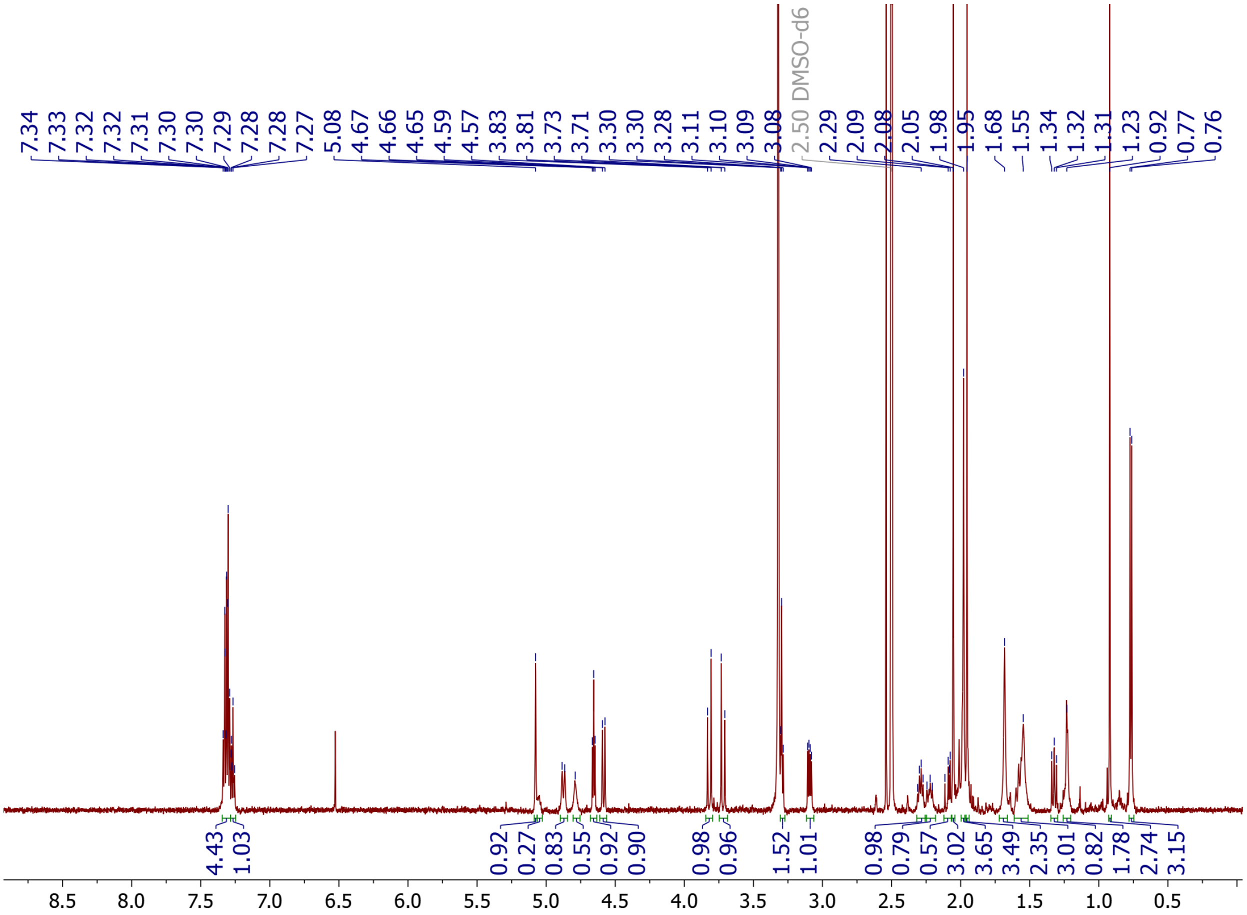


**SI Figure 30.** 600 MHz $^{1}H$ NMR spectrum of compound **2** in DMSO-$d_6$. The spectrum was acquired over 4 scans at 25°C.

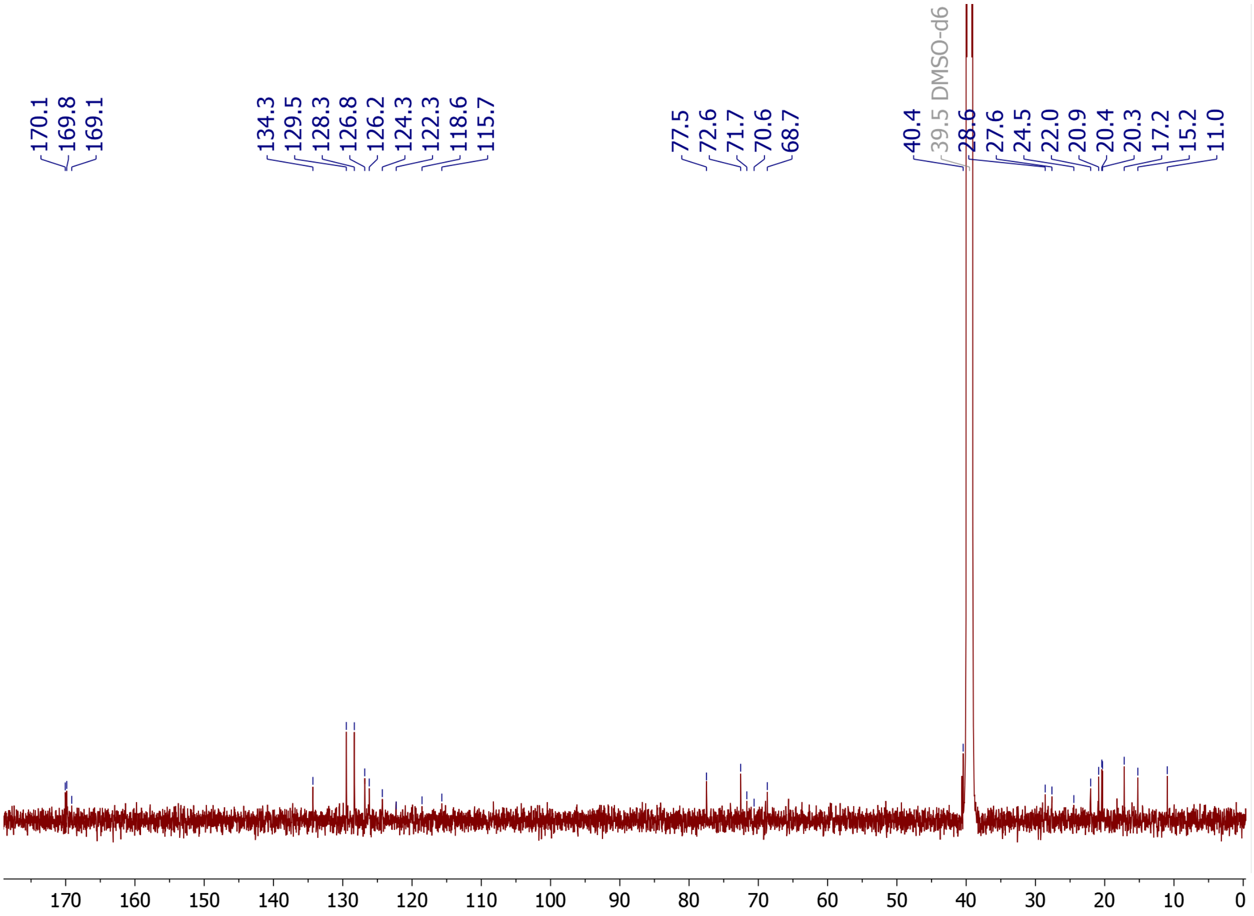


**SI Figure 31.** 151 MHz $^{13}C$ NMR spectrum of compound **2** in DMSO-$d_6$. The spectrum was acquired over 50000 scans at 25°C.

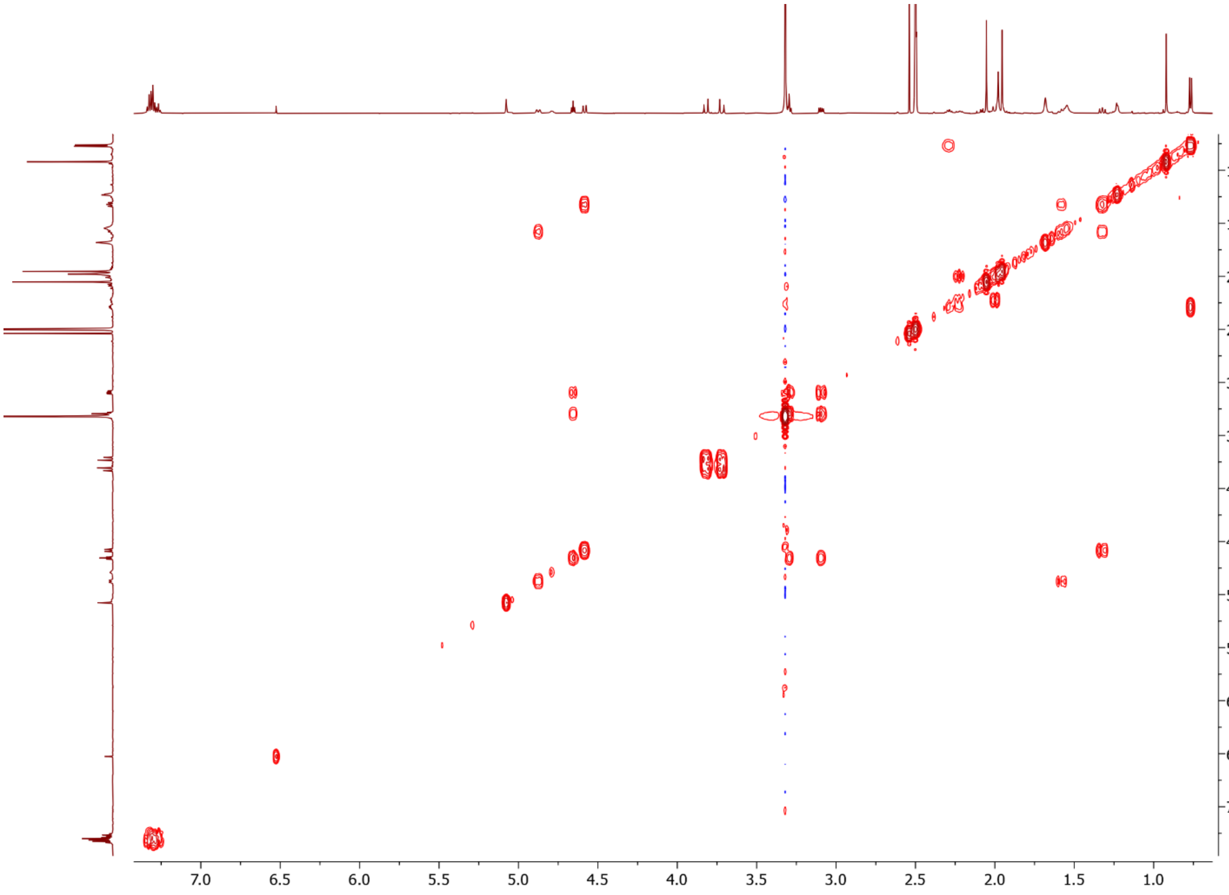


**SI Figure 32.** $^1H$–$^1H$ COSY NMR spectrum of compound **2** in DMSO-$d_6$. The spectrum was acquired over 4 scans at 25°C.

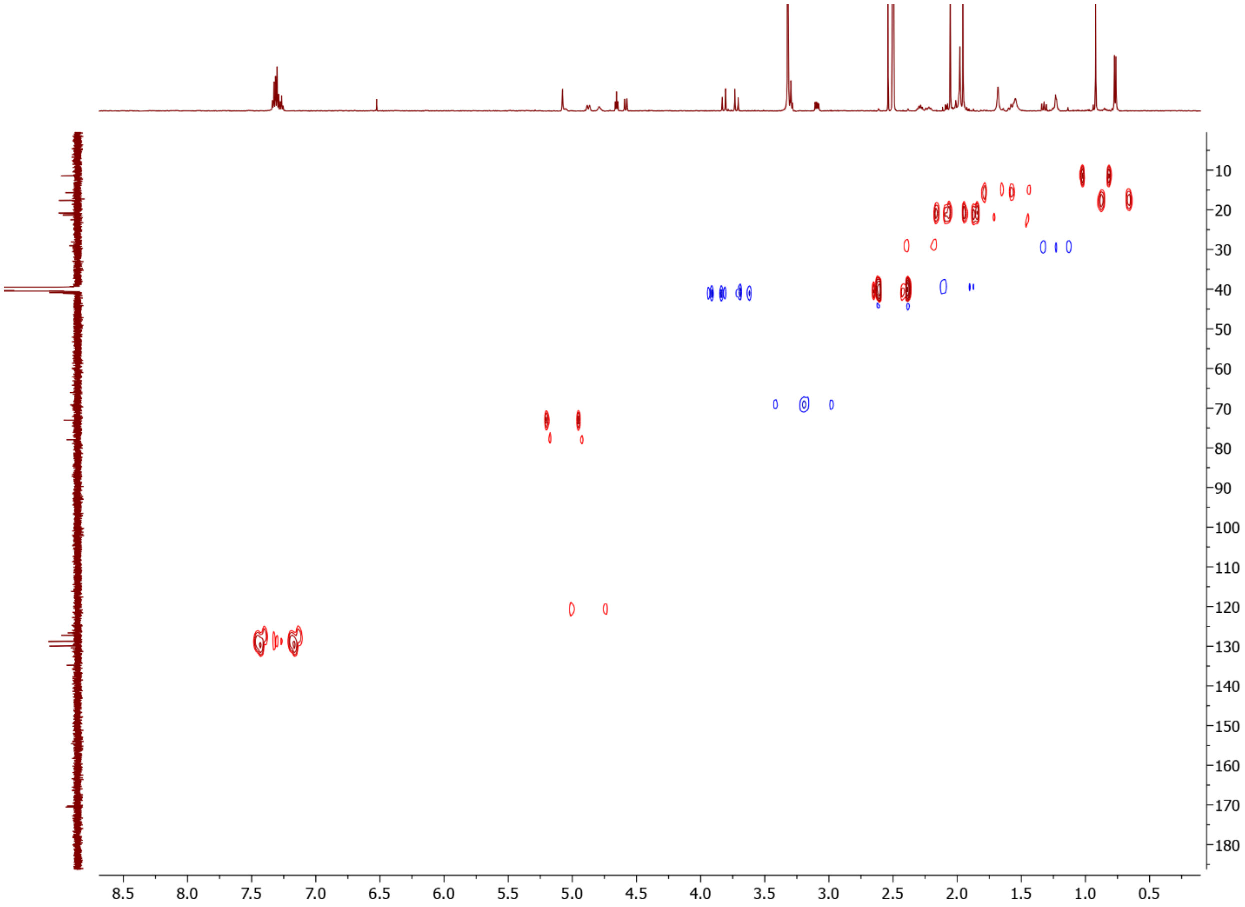


**SI Figure 33.** $^1H$–$^{13}C$ HSQC coupled NMR spectrum of compound **2** in DMSO-$d_6$. The spectrum was acquired over 128 scans at 25 °C with the decoupler turned off.

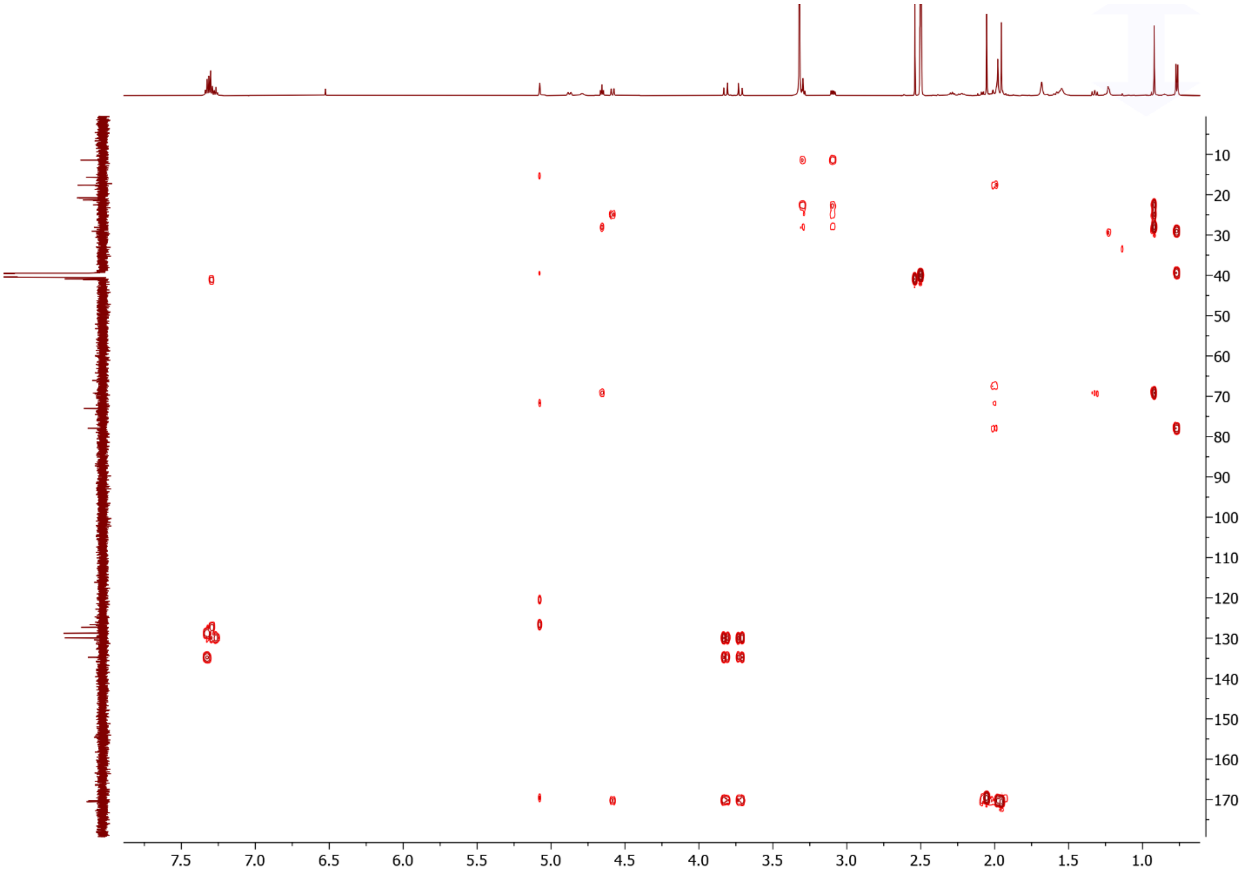


**SI Figure 34.** $^{1}H$–$^{13}C$ HMBC coupled NMR spectrum of compound **2** in DMSO-$d_6$. The spectrum was acquired over 256 scans at 25 °C.

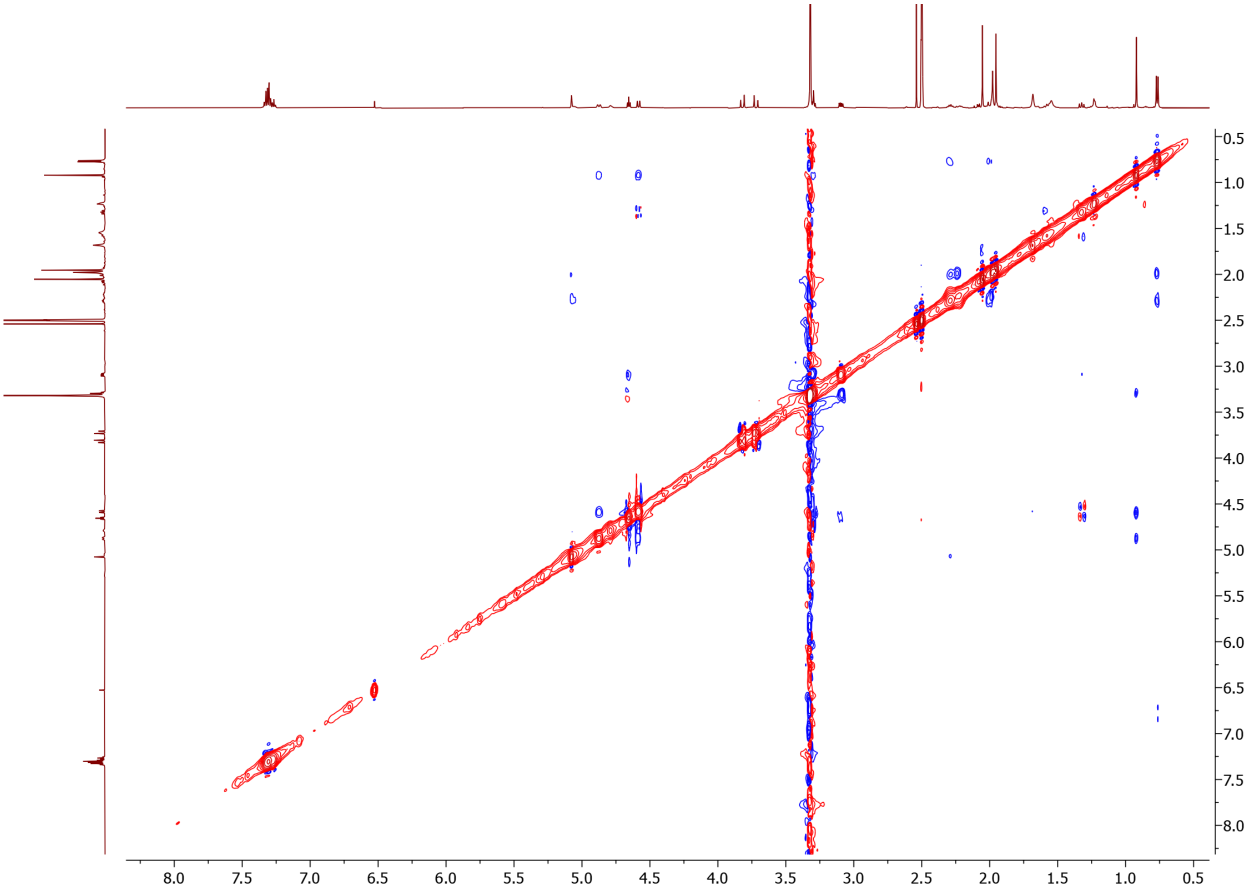


**SI Figure 35.** 2D ROESY NMR spectrum of compound **2** in DMSO-$d_6$. The spectrum was acquired over 64 scans at 25 °C.

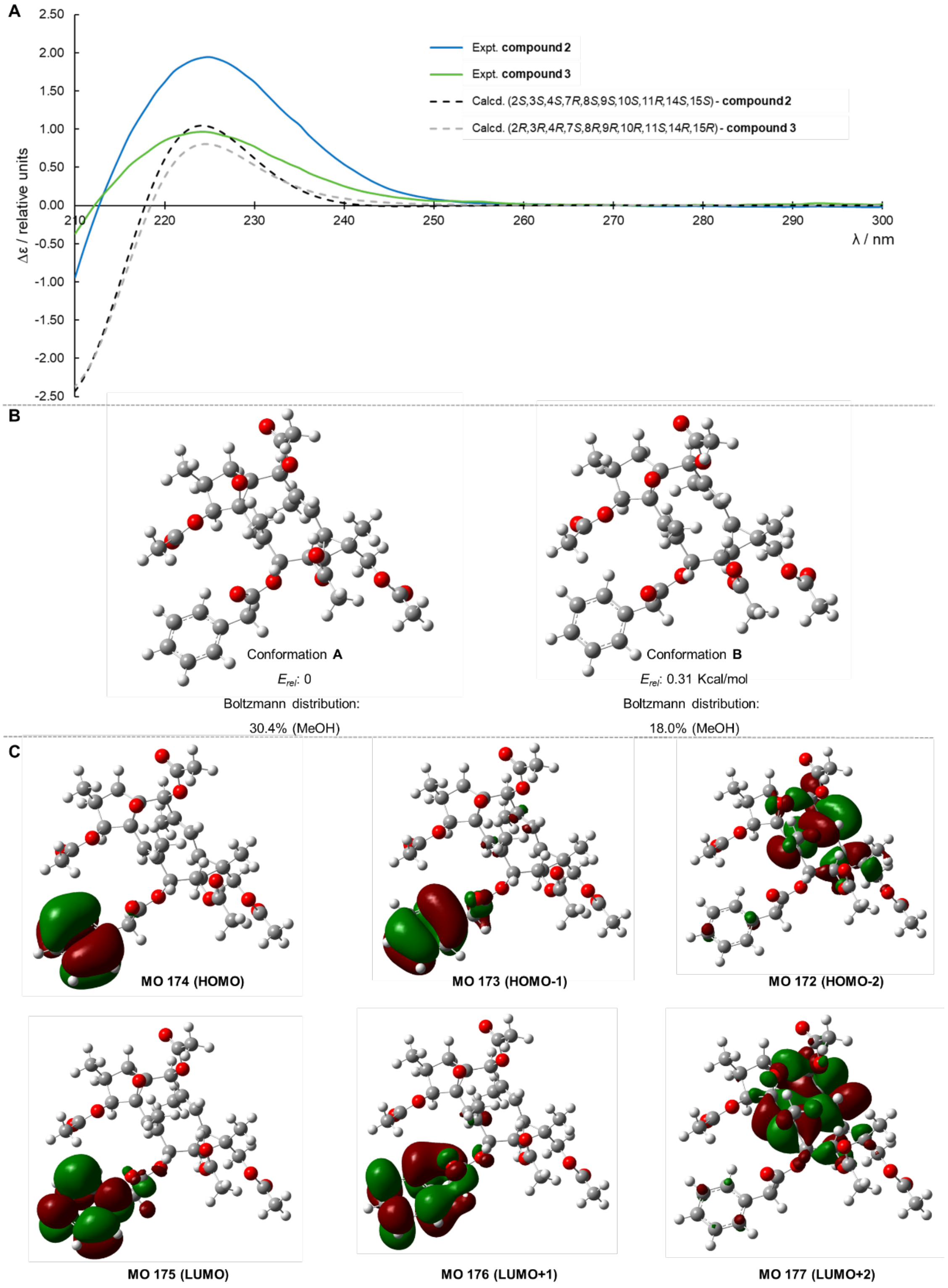


**SI Figure 36.** (**A**) Calculated (dashed lines) and experimental ECD spectra of compound **3** (green line) and compound **2** (blue line). (**B**) Lowest-energy conformers for compound **3**. (**C**) Key molecular orbitals involved in important transitions regarding the ECD spectra of the dominant conformer of compound **3** in methanol at the CAM-B3LYP/dgtzvp level.

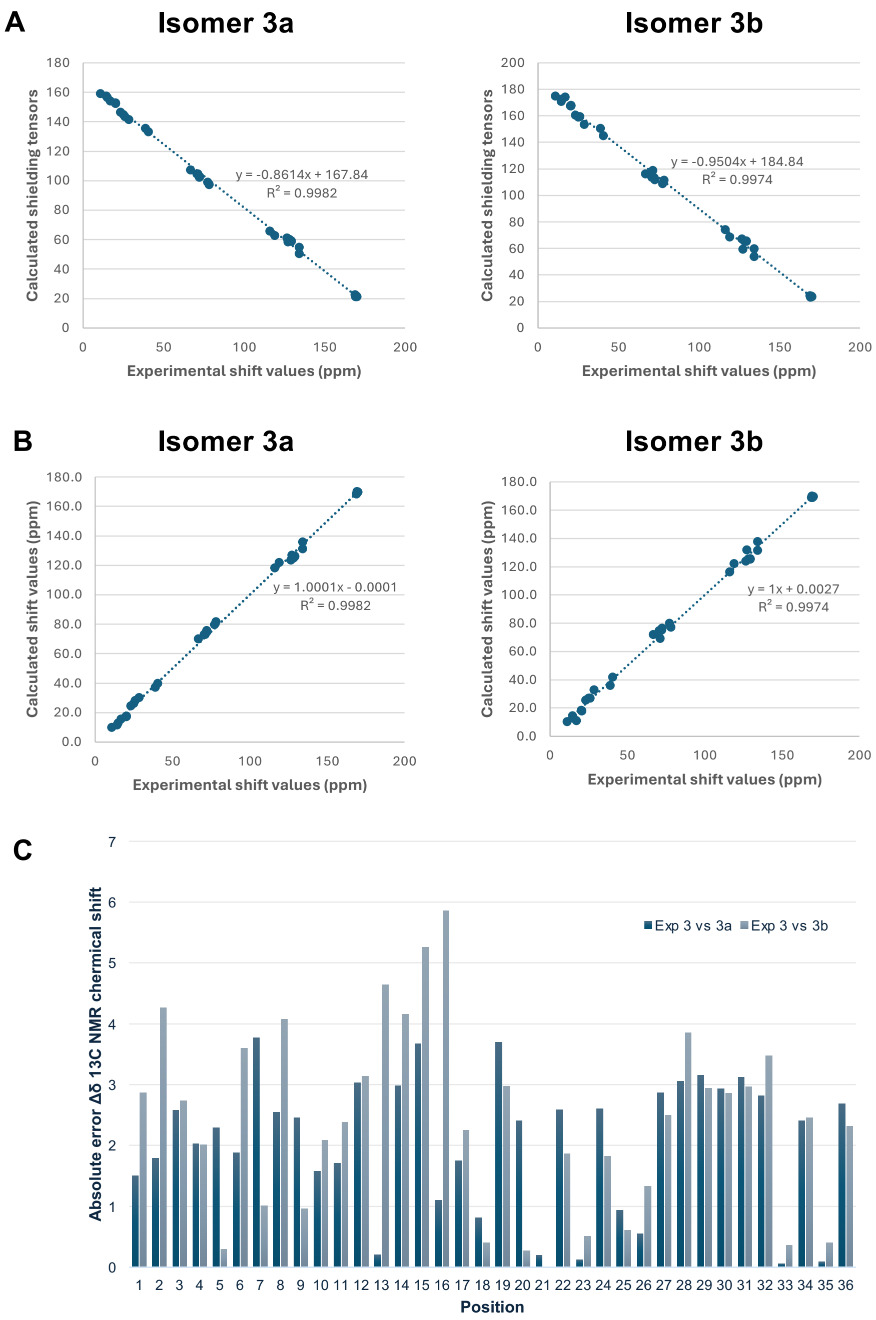


**SI Figure 37.** (**A**) Correlation between the experimental $^{13}C$ NMR chemical shifts for compound **3** and isotropic shielding tensors obtained using B3LYP hybrid functional and 6-31G(d) basis set in DMSO. (**B**) Correlation between experimental and calculated $^{13}C$ NMR chemical shifts obtained through linear regression between the experimental $^{13}C$ NMR chemical shifts and the isotropic shielding tensors of the isomers **3a** and **3b** respectively. (**C**) Experimental $^{13}C$ NMR and DFT-calculated data absolute error for isomers **3a** and **3b**.

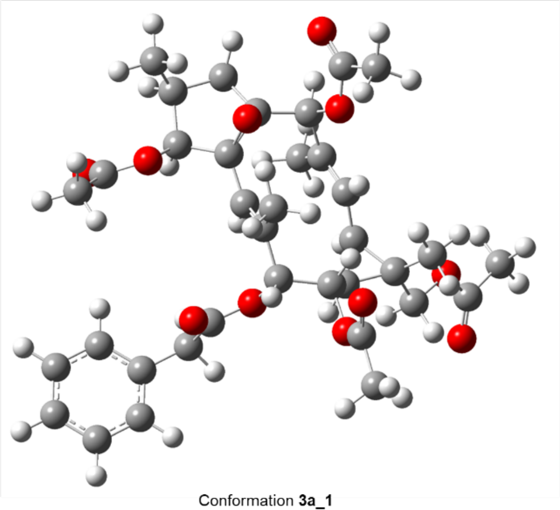

Conformation **3a_1**

$E_{rel}$: 0

Boltzmann distribution:

20.8%

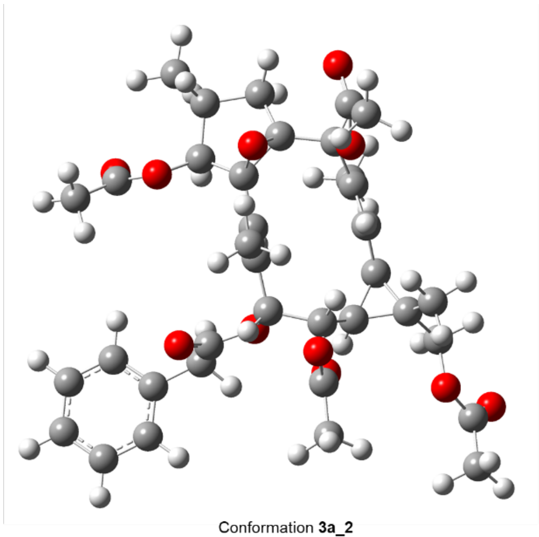

Conformation **3a_2**

$E_{rel}$: 0.10

Boltzmann distribution:

18.3%

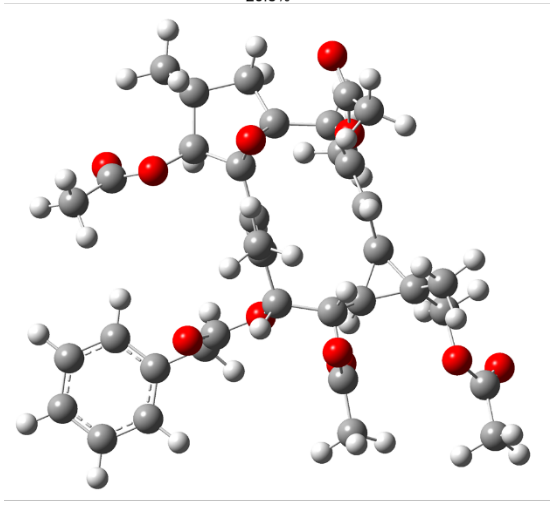

Conformation **3a_3**

$E_{rel}$: 0.10 Kcal/mol

Boltzmann distribution:

17.4%

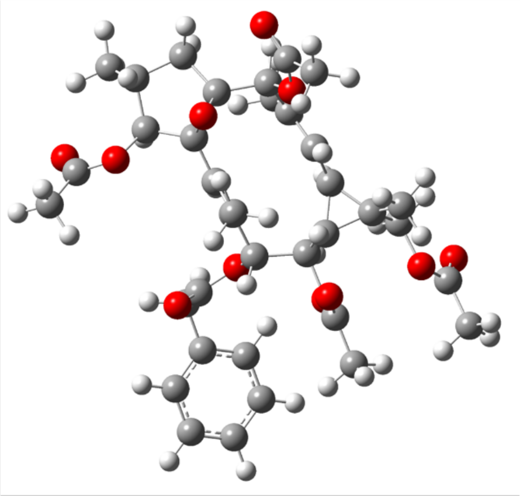

Conformation **3a_4**

$E_{rel}$: 0.63 Kcal/mol

Boltzmann distribution:

7.1%

**SI Figure 38.** The four most relevant conformers of isomer **3a** with (2*S*,3*S*,4*S*,7*R*,8*S*,7*R*,8*S*,9*S*,10*S*,11*R*,14*S*,15*S*) configuration for compound **3** modeled in DMSO.

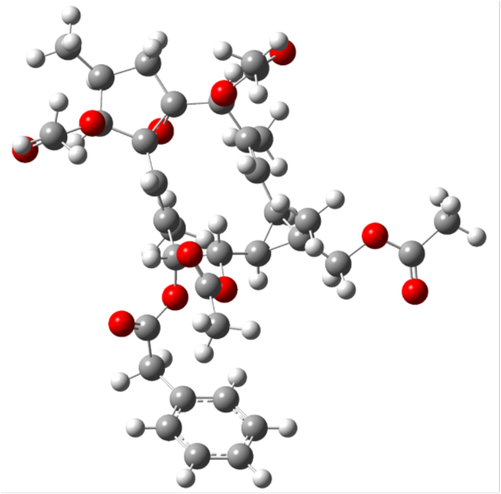

Conformation **3b_1**
$E_{rel}$: 0
Boltzmann distribution:
57.4%

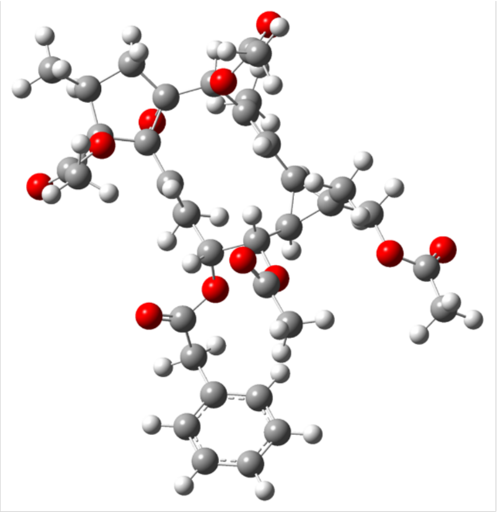

Conformation **3b_2**
$E_{rel}$: 1.14
Boltzmann distribution:
8.3%

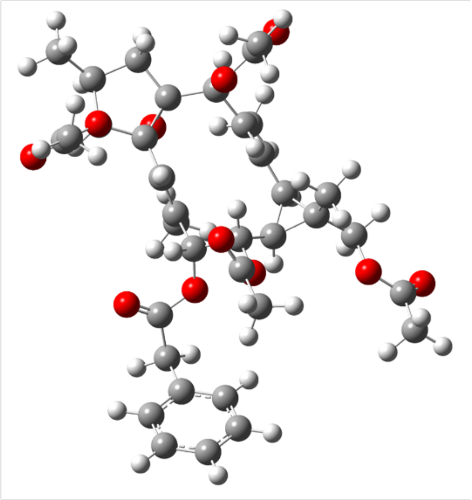

Conformation **3c_3**
$E_{rel}$: 1.16 Kcal/mol
Boltzmann distribution:
8.1%

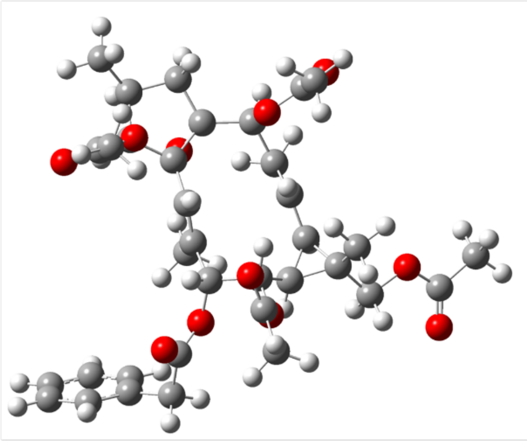

Conformation **3d_4**
$E_{rel}$: 1.28 Kcal/mol
Boltzmann distribution:
6.6%

**SI Figure 39.** The four most relevant conformers of isomer 3b with (2*S*,3*S*,4*R*,7*R*,8*S*,7*R*,8*S*,9*S*,10*S*,11*R*,14*S*,15*R*) configuration for compound **3** modeled in DMSO.

### Supplemental Data Table Legends

**Table S1**: List of confirmed hit fractions, their taxonomic information, and their locational origin.

### Supplemental References


1. V. Koduri *et al.*, Targeting oncoproteins with a positive selection assay for protein degraders. *Sci Adv* **7**, (2021).
2. S. Hirai *et al.*, Ras-dependent signal transduction is indispensable but not sufficient for the activation of AP1/Jun by PKC delta. *EMBO J* **13**, 2331-2340 (1994).
3. W. C. Chan *et al.*, Accelerating inhibitor discovery for deubiquitinating enzymes. *Nat Commun* **14**, 686 (2023).
4. T. Grkovic *et al.*, National Cancer Institute (NCI) Program for Natural Products Discovery: Rapid Isolation and Identification of Biologically Active Natural Products from the NCI Prefractionated Library. *ACS Chem Biol* **15**, 1104-1114 (2020).
5. K. A. Donovan *et al.*, Thalidomide promotes degradation of SALL4, a transcription factor implicated in Duane Radial Ray syndrome. *Elife* **7**, (2018).
6. V. Demichev, C. B. Messner, S. I. Vernardis, K. S. Lilley, M. Ralser, DIA-NN: neural networks and interference correction enable deep proteome coverage in high throughput. *Nature methods* **17**, 41-44 (2020).
7. K. Baek *et al.*, Unveiling the hidden interactome of CRBN molecular glues. *Nature Communications* **16**, 6831 (2025).
8. A. Nabbi, K. Riabowol, Rapid Isolation of Nuclei from Cells In Vitro. *Cold Spring Harb Protoc* **2015**, 769-772 (2015).
9. A. Dobin *et al.*, STAR: ultrafast universal RNA-seq aligner. *Bioinformatics* **29**, 15-21 (2013).
10. R. Patro, G. Duggal, M. I. Love, R. A. Irizarry, C. Kingsford, Salmon provides fast and bias-aware quantification of transcript expression. *Nat Methods* **14**, 417-419 (2017).
11. M. I. Love, W. Huber, S. Anders, Moderated estimation of fold change and dispersion for RNA-seq data with DESeq2. *Genome Biol* **15**, 550 (2014).
12. M. Cornwell *et al.*, VIPER: Visualization Pipeline for RNA-seq, a Snakemake workflow for efficient and complete RNA-seq analysis. *BMC Bioinformatics* **19**, 135 (2018).
13. A. Liberzon *et al.*, The Molecular Signatures Database (MSigDB) hallmark gene set collection. *Cell Syst* **1**, 417-425 (2015).
14. M. V. Kuleshov *et al.*, Enrichr: a comprehensive gene set enrichment analysis web server 2016 update. *Nucleic Acids Res* **44**, W90-97 (2016).
15. J.-Y. C. Zhu, B.; Zheng, Y.-J.; Dong, Z.; Lin, S.-L.; Tang, G.-H.; Gu, Q.; Yin, S. , Enantiomeric neolignans and sesquineolignans from Jatropha integerrima and their absolute configurations. *RSC Adv.* **5**, 12202-12208 (2015).
16. C. C. Thornburg *et al.*, NCI Program for Natural Product Discovery: A Publicly-Accessible Library of Natural Product Fractions for High-Throughput Screening. *ACS Chem Biol* **13**, 2484-2497 (2018).
17. Y. Gao *et al.*, Phonerilins A–K, cytotoxic ingenane and ingol diterpenoids from Euphorbia neriifolia. *Tetrahedron* **123**, (2022).
18. W. Y. Qi *et al.*, Ingol-type diterpenes from Euphorbia antiquorum with mouse 11beta-hydroxysteroid dehydrogenase type 1 inhibition activity. *J Nat Prod* **77**, 1452-1458 (2014).
19. J. A. Marco, J. F. Sanz-Cervera, F. J. Ropero, J. Checa, B. M. Fraga, Ingenane and lathyrane diterpenes from the latex of euphorbia acrurensis. *Phytochemistry* **49**, 1095-1099 (1998).

20. N. Grimblat, M. M. Zanardi, A. M. Sarotti, Beyond DP4: an Improved Probability for the Stereochemical Assignment of Isomeric Compounds using Quantum Chemical Calculations of NMR Shifts. *J Org Chem* **80**, 12526-12534 (2015).
21. Z. S. Safi, N. Wazzan, DFT calculations of (1)H- and (13)C-NMR chemical shifts of 3-methyl-1-phenyl-4-(phenyldiazenyl)-1H-pyrazol-5-amine in solution. *Sci Rep* **12**, 17798 (2022).
